# The mutational constraint spectrum quantified from variation in 141,456 humans

**DOI:** 10.1101/531210

**Authors:** Konrad J. Karczewski, Laurent C. Francioli, Grace Tiao, Beryl B. Cummings, Jessica Alföldi, Qingbo Wang, Ryan L. Collins, Kristen M. Laricchia, Andrea Ganna, Daniel P. Birnbaum, Laura D. Gauthier, Harrison Brand, Matthew Solomonson, Nicholas A. Watts, Daniel Rhodes, Moriel Singer-Berk, Eleina M. England, Eleanor G. Seaby, Jack A. Kosmicki, Raymond K. Walters, Katherine Tashman, Yossi Farjoun, Eric Banks, Timothy Poterba, Arcturus Wang, Cotton Seed, Nicola Whiffin, Jessica X. Chong, Kaitlin E. Samocha, Emma Pierce-Hoffman, Zachary Zappala, Anne H. O’Donnell-Luria, Eric Vallabh Minikel, Ben Weisburd, Monkol Lek, James S. Ware, Christopher Vittal, Irina M. Armean, Louis Bergelson, Kristian Cibulskis, Kristen M. Connolly, Miguel Covarrubias, Stacey Donnelly, Steven Ferriera, Stacey Gabriel, Jeff Gentry, Namrata Gupta, Thibault Jeandet, Diane Kaplan, Christopher Llanwarne, Ruchi Munshi, Sam Novod, Nikelle Petrillo, David Roazen, Valentin Ruano-Rubio, Andrea Saltzman, Molly Schleicher, Jose Soto, Kathleen Tibbetts, Charlotte Tolonen, Gordon Wade, Michael E. Talkowski, Genome Aggregation Database (gnomAD) Consortium, Benjamin M. Neale, Mark J. Daly, Daniel G. MacArthur

## Abstract

Genetic variants that inactivate protein-coding genes are a powerful source of information about the phenotypic consequences of gene disruption: genes critical for an organism’s function will be depleted for such variants in natural populations, while non-essential genes will tolerate their accumulation. However, predicted loss-of-function (pLoF) variants are enriched for annotation errors, and tend to be found at extremely low frequencies, so their analysis requires careful variant annotation and very large sample sizes^1^. Here, we describe the aggregation of 125,748 exomes and 15,708 genomes from human sequencing studies into the Genome Aggregation Database (gnomAD). We identify 443,769 high-confidence pLoF variants in this cohort after filtering for sequencing and annotation artifacts. Using an improved human mutation rate model, we classify human protein-coding genes along a spectrum representing tolerance to inactivation, validate this classification using data from model organisms and engineered human cells, and show that it can be used to improve gene discovery power for both common and rare diseases.

---

The physiological function of most genes in the human genome remains unknown. In biology, as in many engineering and scientific fields, breaking the individual components of a complex system can provide valuable insight into the structure and behavior of that system. For discovery of gene function, a common approach is to introduce disruptive mutations into genes and assay their effects on cellular and physiological phenotypes in mutant organisms or cell lines^2^. Such studies have yielded valuable insight into eukaryotic physiology and have guided therapeutic design^3^. However, while model organism and human cell studies have been crucial in deciphering the function of many human genes, they remain imperfect proxies for human physiology.

Obvious ethical and technical constraints prevent the large-scale engineering of LoF mutations in humans. However, recent exome and genome sequencing projects have revealed a surprisingly high burden of natural pLoF variation in the human population, including stop-gained, essential splice, and frameshift variants^1,4^, which can serve as natural models for human gene inactivation. Such variants have already revealed much about human biology and disease mechanisms, through many decades of study of the genetic basis of severe Mendelian diseases^5^, most of which are driven by disruptive variants in either the heterozygous or homozygous state. These variants have also proved valuable in identifying potential therapeutic targets: confirmed LoF variants in *PCSK9* have been causally linked to low LDL cholesterol levels^6^, leading ultimately to the development of multiple PCSK9 inhibitors now in clinical use for the reduction of cardiovascular disease risk. A systematic catalog of pLoF variants in humans and classification of genes along a spectrum of tolerance to inactivation would provide a valuable resource for medical genetics, identifying candidate disease-causing mutations, potential therapeutic targets, and windows into the normal function of many currently uncharacterized human genes.

A number of challenges arise when assessing LoF variants at scale. LoF variants are on average deleterious, and are thus typically maintained at very low frequencies in the human population. Systematic genome-wide discovery of these variants requires whole exome or whole genome sequencing of very large numbers of samples. Additionally, LoF variants are enriched for false positives compared to synonymous or other benign variants, including mapping, genotyping (including somatic variation), and particularly, annotation errors^1^, and careful filtering is required to remove such artifacts.

Population surveys of coding variation enable the evaluation of the strength of natural selection at a gene or region level. As natural selection purges deleterious variants from human populations, methods to detect selection have modelled the reduction in variation (constraint) ^7^ or shift in the allele frequency distribution^8^, compared to an expectation. For analyses of selection on coding variation, synonymous variation provides a convenient baseline, controlling for other potential population genetic forces that may influence the amount of variation as well as technical features of the local sequence. We have previously applied a model of constraint to define a set of 3,230 genes with a high probability of intolerance to heterozygous pLoF variation (pLI)^4^ and estimated the selection coefficient for variants these genes^9^. However, the ability to comprehensively characterize the degree of selection against pLoF variants is particularly limited, as for small genes, the expected number of mutations is still small, even for samples of up to 60,000 individuals^4,10^. Further, the previous dichotomization of pLI, although convenient for characterization of a set of genes, disguises variability in the degree of selective pressure against a given class of variation and overlooks more subtle levels of intolerance to pLoF variation. With larger sample sizes, a more accurate quantitative measure of selective pressure is a possibility.

Here, we describe the detection of pLoF variants in a cohort of 125,748 individuals with whole exome sequence data and 15,708 individuals with whole genome sequence data, as part of the Genome Aggregation Database (gnomAD; https://gnomad.broadinstitute.org), the successor to the Exome Aggregation Consortium (ExAC). We develop a continuous measure of intolerance to pLoF variation, which places each gene on a spectrum of LoF intolerance. We validate this metric by comparing its distribution to several orthogonal indicators of constraint, including the incidence of structural variation and the essentiality of genes as measured through mouse gene knockout experiments and cellular inactivation assays. Finally, we demonstrate that this metric improves interpretation of genetic variants influencing rare disease and provides insight into common disease biology. These analyses provide the most comprehensive catalog to date of the sensitivity of human genes to disruption.

In a series of accompanying manuscripts, we also describe other complementary analyses of this data set. Using an overlapping set of 14,237 whole genomes, we report the discovery and characterization of a wide variety of structural variants (large deletions, duplications, insertions, or other rearrangements of DNA)^11^. We explore the value of pLoF variants for the discovery and validation of therapeutic drug targets^12^, and provide a case study of the use of these variants from gnomAD and other large reference data sets to validate the safety of inhibition of LRRK2, a candidate therapeutic target for Parkinson’s disease^13^. By combining the gnomAD data set with a large collection of RNA sequencing data from adult human tissues^14^, we demonstrate the value of tissue expression data in the interpretation of genetic variation across a range of human diseases^15^. Finally, we characterize and investigate the impact of two understudied classes of human variation: multi-nucleotide variants^16^ and variants creating or disrupting open reading frames in the 5’ untranslated region of human genes^17^.

## A high-quality catalogue of variation

We aggregated whole exome sequencing data from 199,558 individuals and whole genome sequencing data from 20,314 individuals. These data were obtained primarily from case-control studies of adult-onset common diseases, including cardiovascular disease, type 2 diabetes, and psychiatric disorders. Each dataset, totaling over 1.3 and 1.6 petabytes of raw sequencing data respectively, was uniformly processed, joint variant calling was performed on each dataset using a standardized BWA-Picard-GATK pipeline, and all data processing and analysis was performed using Hail^18^. We performed stringent sample QC (Extended Data Fig. 1), removing samples with lower sequencing quality by a variety of metrics, second-degree or closer related individuals across both data types, samples with inadequate consent for release of aggregate data, and individuals known to have a severe childhood onset disease as well as their first-degree relatives. The final gnomAD release contains genetic variation from 125,748 exomes and 15,708 genomes from unique unrelated individuals with high-quality sequence data, spanning 6 global and 8 sub-continental ancestries (Fig. 1a,b), which we have made publicly available at https://gnomad.broadinstitute.org. We also provide subsets of the gnomAD datasets, which exclude individuals who are cases in case-control studies, or who are cases of a few particular disease types such as cancer and neurological disorders, or who are also aggregated in the Bravo TOPMed variant browser (https://bravo.sph.umich.edu).

**Figure 1.**
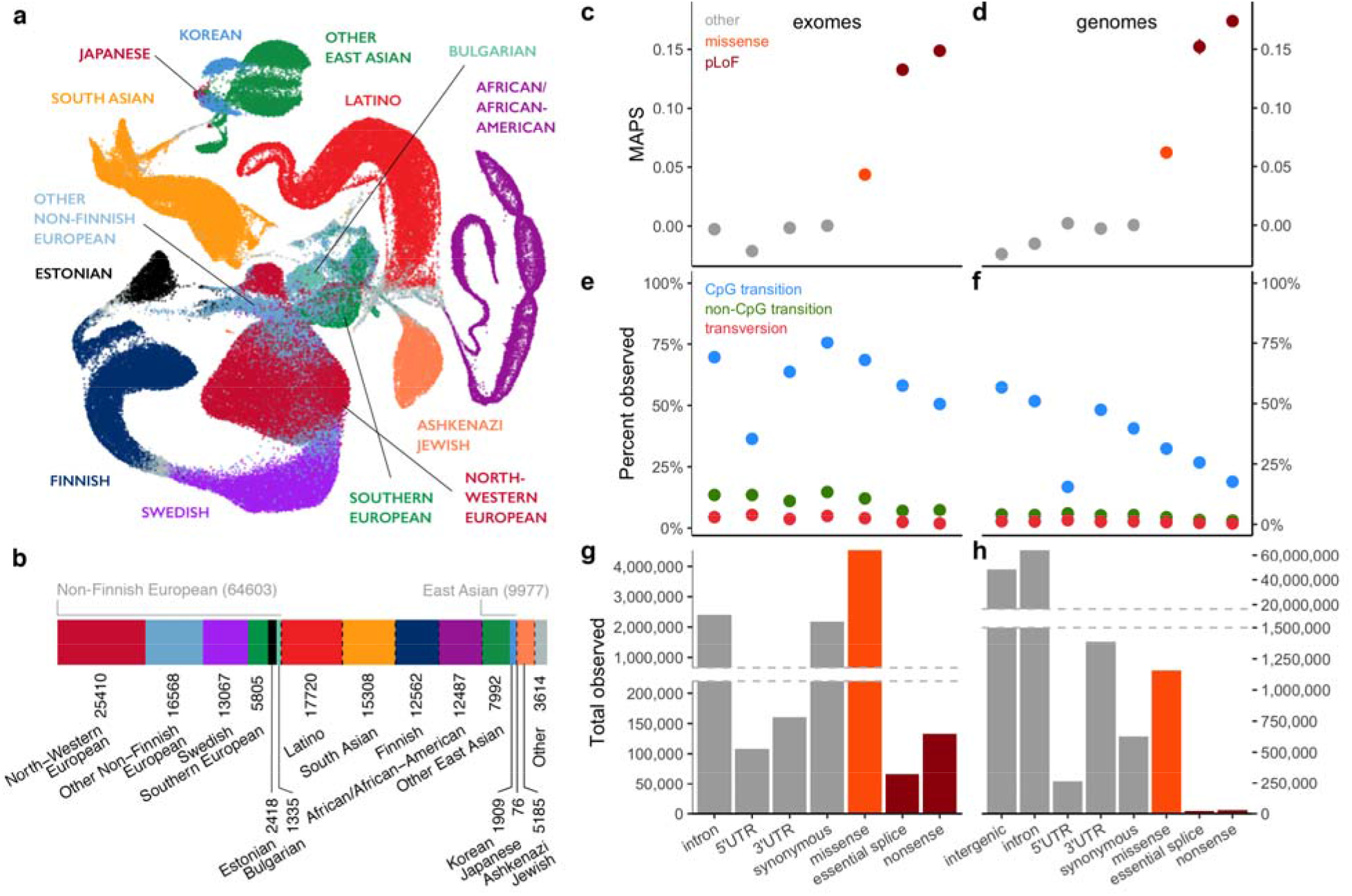
Aggregation of 141,456 exome and genome sequences. **a,** Uniform Manifold Approximation and Projection (UMAP)^19,20^ plot depicting the ancestral diversity of all individuals in gnomAD, using 10 principal components. Note that long-range distances in UMAP space are not a proxy for genetic distance. **b**, The number of individuals by population and subpopulation in the gnomAD database. Colors representing populations in **a & b** are consistent. **c-d,** The mutability-adjusted proportion of singletons^4^ (MAPS) is shown across functional categories for SNVs in exomes (**c**, x-axis shared with **e** and **g**) and genomes (**d**, x-axis shared with **f** and **h**). Higher values indicate an enrichment of lower frequency variants, suggesting increased deleteriousness. **e-f,** The proportion of possible variants observed for each functional class for each mutational type for exomes (**e**) and genomes (**f**). CpG transitions are more saturated, except where selection (e.g. pLoFs), or hypomethylation (5’UTR) decreases the number of observations. **g-h,** The total number of variants observed in each functional class for exomes (**g**) and genomes (**h**). Error bars in (**c-f**) represent 95% confidence intervals (note that in some cases these are fully contained within the plotted point).

Among these individuals, we discovered 17.2 million and 261.9 million variants in the exome and genome datasets, respectively; these variants were filtered using a custom random forest process (Supplementary Information) to 14.9 million and 229.9 million high-quality variants. Comparing our variant calls in two samples for which we had independent gold-standard variant calls, we found that our filtering achieves very high precision (>99% for single nucleotide variants (SNVs), >98.5% for indels in both exomes and genomes) and recall (>90% for SNVs and >82% for indels for both exomes and genomes) at the single sample level (Extended Data Fig. 2). In addition, we leveraged data from 4,568 and 212 trios included in our exome and genome callsets, respectively, to assess the quality of our rare variants. We found that our model retains over 97.8% of the transmitted singletons (singletons in the unrelated individuals that are transmitted to an offspring) on chromosome 20 (which was not used for model training) (Extended Data Fig. 3a-d). In addition, the number of putative *de novo* calls after filtering are in line with expectations^21^ (Extended Data Fig. 3e-h), and our model had a recall of 97.3% for *de novo* SNVs and 98% for *de novo* indels based on 375 independently validated *de novo* variants in our whole-exome trios (295 SNVs and 80 indels, Extended Data Fig. 3i-j). Altogether, these results indicate that our filtering strategy produced a callset with high precision and recall for both common and rare variants.

These variants reflect the expected patterns based on mutation and selection: we observe 84.9% of all possible consistently methylated CpG to TpG transitions that would create synonymous variants in the human exome (Supplementary Table 14), indicating that at this sample size we are beginning to approach mutational saturation of this highly mutable and weakly negatively selected variant class. However, we only observe 52% of methylated CpG stop-gained variants, illustrating the action of natural selection removing a substantial fraction of gene-disrupting variants from the population (Fig. 1c-h). Across all mutational contexts, only 11.5% and 3.7% of the possible synonymous and stop-gained variants, respectively, are observed in the exome dataset, indicating that current sample sizes remain far from capturing complete mutational saturation of the human exome (Extended Data Fig. 4).

## Identifying loss-of-function variants

Some LoF variants will result in embryonic lethality in humans in a heterozygous state, while others are benign even at homozygosity, with a spectrum of effects in between. Throughout this manuscript, we define predicted loss-of-function (pLoF) variants to be those which introduce a premature stop (stop-gained), shift reported transcriptional frame (frameshift), or alter the two essential splice-site nucleotides immediately to the left and right of each exon (splice) found in protein-coding transcripts, and ascertain their presence in the cohort of 125,748 individuals with exome sequence data. As these variants are enriched for annotation artifacts^1^, we developed the Loss-Of-Function Transcript Effect Estimator (LOFTEE) package, which applies stringent filtering criteria from first principles (such as removing terminal truncation variants, as well as rescued splice variants, that are predicted to escape nonsense-mediated decay) to pLoF variants annotated by the Variant Effect Predictor (Extended Data Fig. 5a). Despite not using frequency information, we find that this method disproportionately removes pLoF variants that are common in the population, which are known to be enriched for annotation errors^1^, while retaining rare, likely deleterious variation, as well as reported pathogenic variation (Fig. 2a). LOFTEE distinguishes high-confidence pLoF variants from annotation artifacts, and identifies a set of putative splice variants outside the essential splice site. LOFTEE’s filtering strategy is conservative in the interest of increasing specificity, filtering some potentially functional variants that display a frequency spectrum consistent with that of missense variation (Fig. 2b). Applying LOFTEE v1.0, we discover 443,769 high-confidence pLoF variants, of which 413,097 fall on the canonical transcripts of 16,694 genes. The number of pLoF variants per individual is consistent with previous reports^1^, and is highly dependent on the frequency filters chosen (Supplementary Table 17).

**Figure 2.**
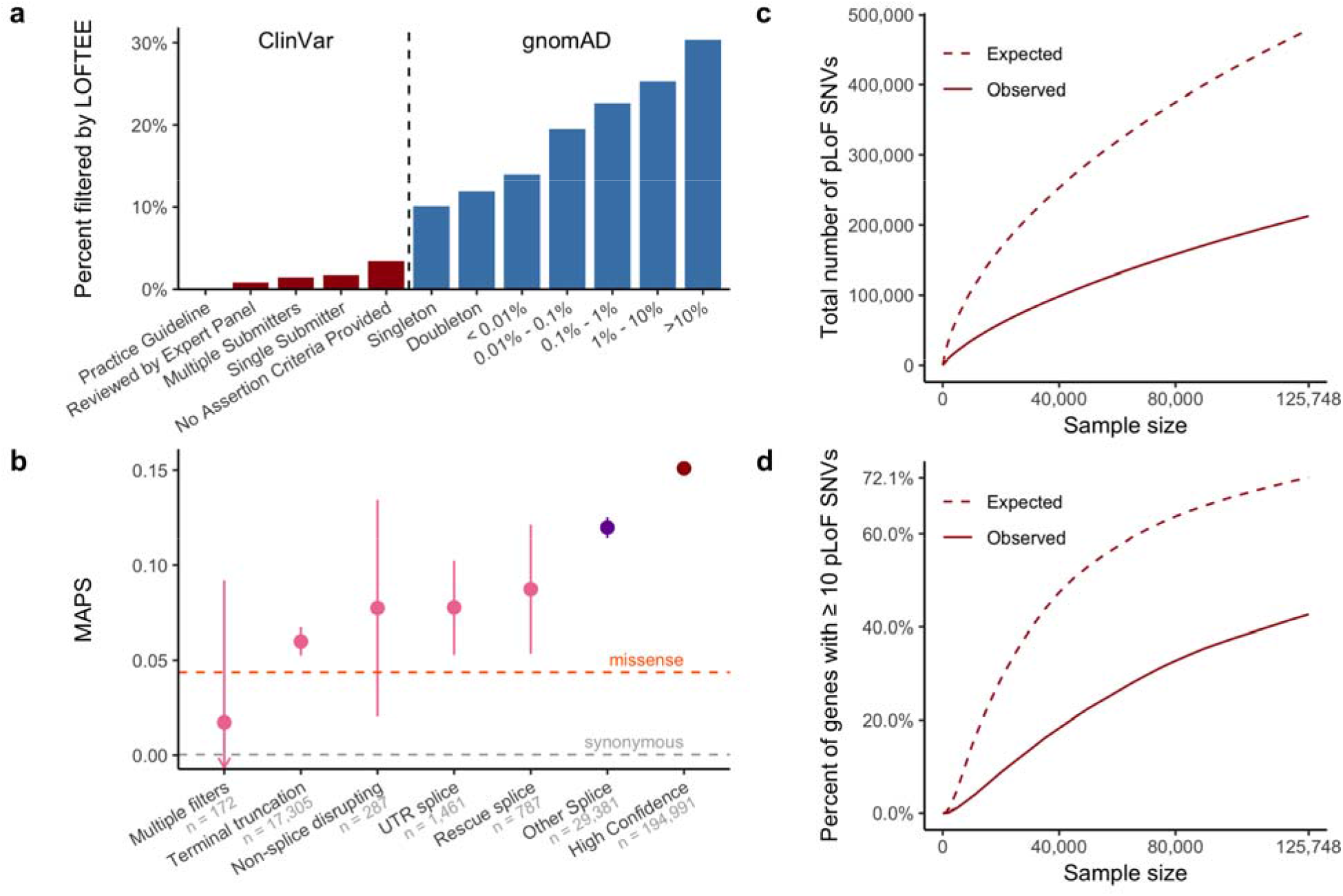
Generating a high-confidence set of predicted loss-of-function (pLoF) variants. **a,** The percent of variants filtered by LOFTEE grouped by ClinVar status and gnomAD frequency. Despite not using frequency information, LOFTEE removes a larger proportion of common variants, and a very low proportion of reported disease-causing variation. **b,** MAPS (see Fig. 1c-d) is shown by LOFTEE designation and filter. Variants filtered out by LOFTEE exhibit frequency spectra similar to those of missense variants, predicted splice variants outside the essential splice site are more rare, and high-confidence variants are very likely to be singletons. Only SNVs with at least 80% call rate are included here. Error bars represent 95% confidence intervals. **c, d,** The total number of pLoF variants (**c**), and proportion of genes with over 10 pLoF variants (**d**) observed and expected (in the absence of selection) as a function of sample size (downsampled from gnomAD). Selection reduces the number of variants observed, and variant discovery approximately follows a square-root relationship with number of samples. At current sample sizes, we would expect to identify more than 10 pLoF variants for 72.1% of genes in the absence of selection.

Aggregating across variants, we created a gene-level pLoF frequency metric to estimate the proportion of haplotypes harboring an inactive copy of each gene. We find that 1,555 genes have an aggregate pLoF frequency of at least 0.1% across all individuals in the dataset (Extended Data Fig. 5c), and 3,270 genes have an aggregate pLoF frequency of at least 0.1% in any one population. Further, we characterized the landscape of genic tolerance to homozygous inactivation, identifying 4,332 pLoF variants that are homozygous in at least one individual. Given the rarity of true homozygous LoF variants, we expected substantial enrichment of such variants for sequencing and annotation errors, and we subjected this set to additional filtering and deep manual curation before defining a set of 1,815 genes (2,636 high-confidence variants) that are likely tolerant to biallelic inactivation (Supplementary Dataset 7).

## The LoF intolerance of human genes

Just as a preponderance of pLoF variants is useful for identifying LoF-tolerant genes, we can conversely characterize a gene’s intolerance to inactivation by identifying significant depletions of predicted LoF variation^4,7^. Here, we present a refined mutational model, which incorporates methylation, base-level coverage correction, and LOFTEE (Supplementary Information, Extended Data Fig. 6), in order to predict expected levels of variation under neutrality. Under this updated model, the variation in the number of synonymous variants observed is accurately captured (r = 0.979). We then applied this method to detect depletion of pLoF variation by comparing the number of observed pLoF variants against our expectation in the gnomAD exome data from 125,748 individuals, more than doubling the sample size of ExAC, the previously largest exome collection^4^. For this dataset, we computed a median of 17.9 expected pLoF variants per gene (Fig. 2c) and found that 72.1% of genes have over 10 pLoF variants (powered to be classified into the most constrained genes; see Supplementary Information) expected on the canonical transcript (Fig. 2d), an increase from 13.2 and 62.8%, respectively, in ExAC.

In ExAC, the smaller sample size required a transformation of the observed and expected values for the number of pLoF variants in each gene into the probability of loss-of-function intolerance (pLI): this metric estimates the probability that a gene falls into the class of LoF-haploinsufficient genes (approximately ~10% observed/expected variation) and is ideally used as a dichotomous metric (producing 3,230 genes with pLI > 0.9). Here, our refined model and substantially increased sample size enabled us to directly assess the degree of intolerance to pLoF variation in each gene using the continuous metric of the observed/expected (o/e) ratio and to estimate a confidence interval around the ratio. We find that the median o/e is 48%, indicating that, as noted previously, most genes exhibit at least moderate selection against pLoF variation, and that the distribution of o/e is not dichotomous, but continuous (Extended Data Fig. 7a). For downstream analyses, unless otherwise specified, we use the 90% upper bound of this confidence interval, which we term the loss-of-function observed/expected upper bound fraction (LOEUF; Extended Data Fig. 7b-c), and bin 19,197 genes into deciles of ~1,920 genes each. At current sample sizes, this metric enables the quantitative assessment of constraint with a built-in confidence value, distinguishing small genes (e.g. those with observed = 0, expected = 2; LOEUF = 1.34) from large genes (e.g. observed = 0, expected = 100; LOEUF = 0.03), while retaining the continuous properties of the direct estimate of the ratio (see Supplementary Information). At one extreme of the distribution, we observe genes with a very strong depletion of pLoF variation (first LOEUF decile aggregate o/e = ~6%, Extended Data Fig. 7d) including genes previously characterized as high pLI (Extended Data Fig. 7e). On the other, we find unconstrained genes that are relatively tolerant of inactivation, including many that harbor homozygous pLoF variants (Extended Data Fig. 7f).

We note that the use of the upper bound means that LOEUF is a conservative metric in one direction: genes with low LOEUF scores are confidently depleted for pLoF variation, whereas genes with high LOEUF scores are a mixture of genes without depletion, and genes that are too small to obtain a precise estimate of the o/e ratio. In general, however, the scale of gnomAD means that gene length is rarely a substantive confounder for the analyses described here, and all downstream analyses are adjusted for coding sequence length or filtered to genes with at least 10 expected pLoFs (see Supplementary Information).

## Validation of LoF intolerance score

The LOEUF metric allows us to place each gene along a continuous spectrum of tolerance to inactivation. We examined the correlation of this metric with a number of independent measures of genic sensitivity to disruption. First, we found that LOEUF is consistent with the expected behavior of well-established gene sets: known haploinsufficient genes are strongly depleted of pLoF variation, while olfactory receptors are relatively unconstrained, and genes with a known autosomal recessive mechanism, for which selection against heterozygous disruptive variants tends to be present but weak^9^, fall in the middle of the distribution (Fig. 3a). Additionally, LOEUF is positively correlated with the occurrence of 6,735 rare autosomal deletion structural variants overlapping protein-coding exons identified in a subset of 6,749 individuals with whole genome sequencing data in this manuscript^11^ (Fig. 3b; r = 0.13; p = 9.8 × 10^−68^).

**Figure 3.**
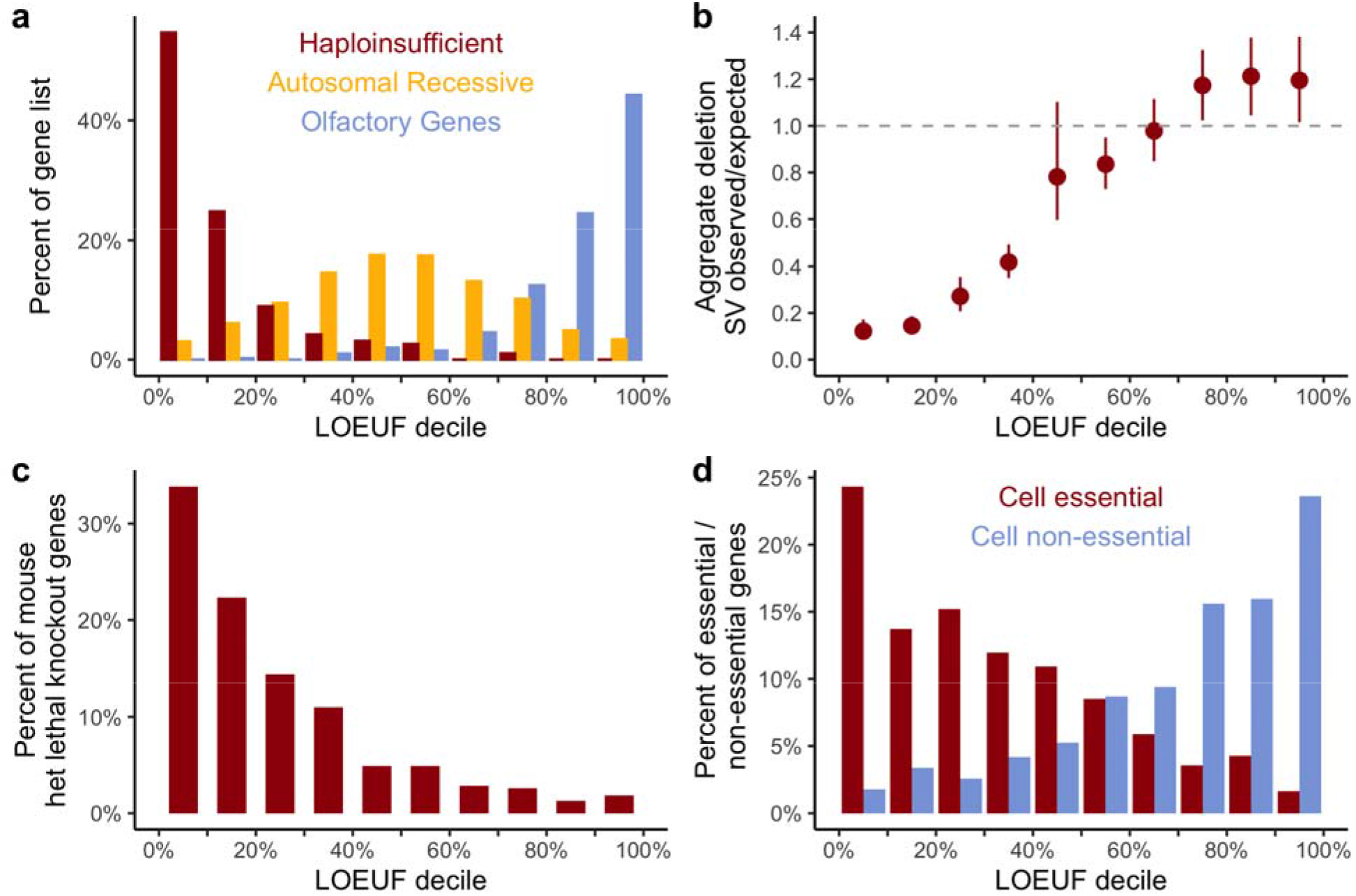
The functional spectrum of pLoF impact. **a,** The percentage of genes in a set of curated gene lists represented in each LOEUF decile. Haploinsufficient genes are enriched among the most constrained genes, while recessive genes are spread in the middle of the distribution, and olfactory receptors are largely unconstrained. **b,** The occurrence of 6,735 rare LoF deletion structural variants (SVs) is correlated with LOEUF (computed from SNVs; linear regression r = 0.13; p = 9.8 × 10^−68^). Error bars represent 95% confidence intervals from bootstrapping. **c, d,** Constrained genes are more likely to be lethal when heterozygously inactivated in mouse (**c**) and cause cellular lethality when disrupted in human cells, while unconstrained genes are more likely to be tolerant of disruption in cellular models (**d**). For all panels, more constrained genes are shown on the left.

This constraint metric also correlates with results in model systems: in 389 genes with orthologs that are embryonically lethal upon heterozygous deletion in mouse^22,23^, we find a lower LOEUF score (mean = 0.488), compared to the remaining 18,808 genes (mean = 0.962; t-test p = 10^−78^; Fig. 3c). Similarly, the 678 genes that are essential for human cell viability as characterized by CRISPR screens^24^ are also depleted for pLoF variation (mean LOEUF = 0.63) in the general population compared to background (18,519 genes with mean LOEUF = 0.964; t-test p = 9 × 10^−71^), while the 777 non-essential genes are more likely to be unconstrained (mean LOEUF = 1.34, compared to remaining 18,420 genes with mean LOEUF = 0.936; t-test p = 3 × 10^−92^; Fig. 3d).

## Biological properties of constraint

We investigated the properties of genes and transcripts as a function of their tolerance to pLoF variation (LOEUF). First, we found that LOEUF correlates with a gene’s degree of connection in protein interaction networks (r = −0.14; p = 1.7 × 10^−51^ after adjusting for gene length, Fig. 4a) and functional characterization (Extended Data Fig. 8a). Additionally, constrained genes are more likely to be ubiquitously expressed across 38 tissues in GTEx (Fig. 4b; LOEUF r = −0.31; p < 1 × 10^−100^) and have higher expression on average (LOEUF ρ = −0.28; p < 1 × 10^−100^), consistent with previous results^4^. While most results in this study are reported at the gene level, we have also extended our framework to compute LOEUF for all protein-coding transcripts, allowing us to explore the extent of differential constraint of transcripts within a given gene. In cases where a gene contained transcripts with varying levels of constraint, we found that transcripts in the first LOEUF decile were more likely to be expressed across tissues than others in the same gene (n = 1,740 genes), even when adjusted for transcript length (Fig. 4c; constrained transcripts are on average 6.34 TPM higher; p = 2.2 × 10^−14^). Additionally, we found that the most constrained transcript for each gene was typically the most highly expressed transcript in tissues with disease relevance^25^ (Extended Data Fig. 8c), supporting the need for transcript-based variant interpretation, as explored in more depth in an accompanying manuscript^15^.

**Figure 4.**
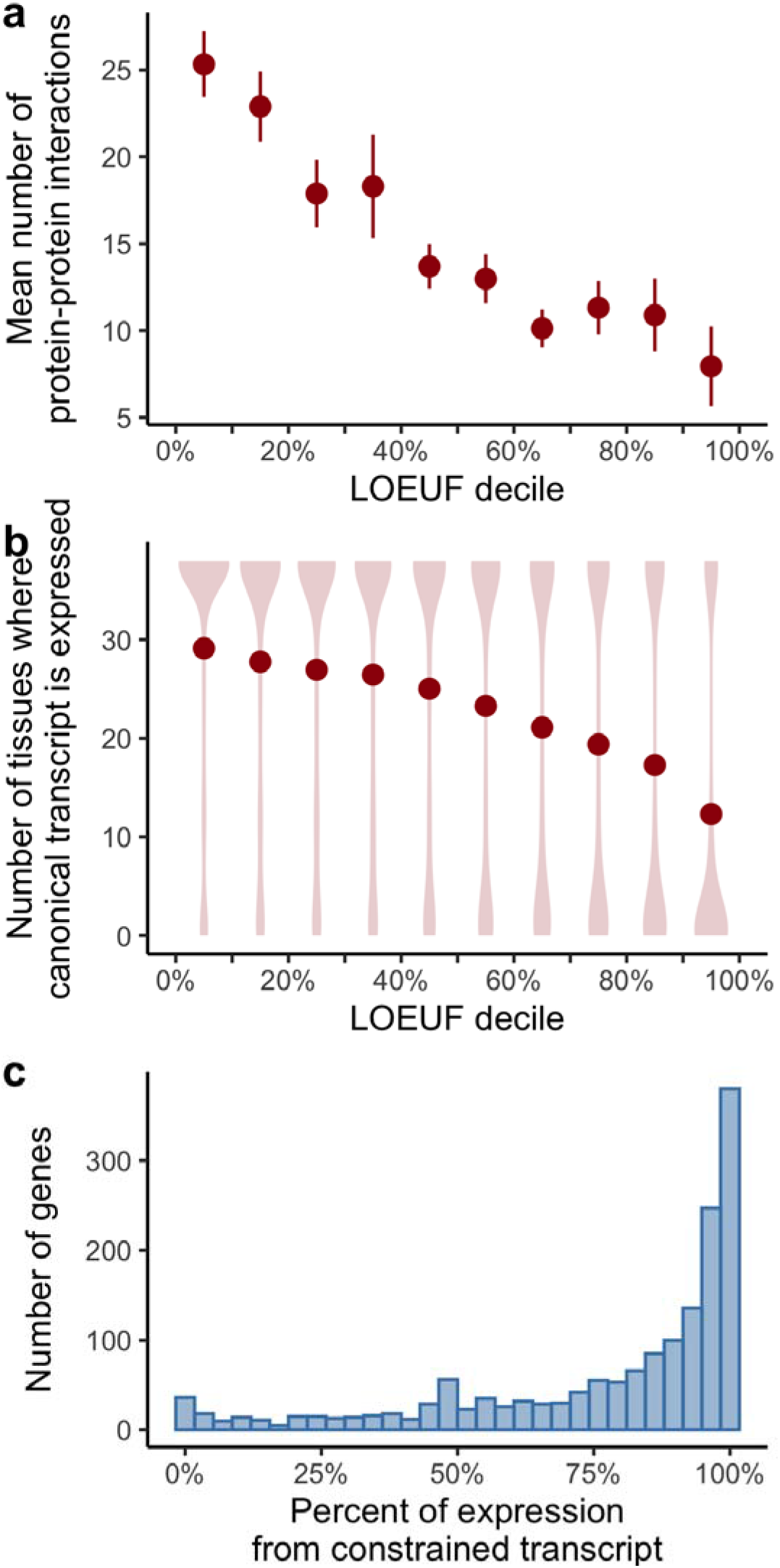
Biological properties of constrained genes and transcripts. **a,** The mean number of protein-protein interactions is plotted as a function of LOEUF decile: more constrained genes have more interaction partners (LOEUF linear regression r = −0.14; p = 1.7 × 10^−51^). Error bars correspond to 95% confidence intervals. **b,** The number of tissues where a gene is expressed (TPM > 0.3), binned by LOEUF decile, is shown as a violin plot with the mean number overlaid as points: more constrained genes are more likely to be expressed in multiple tissues (LOEUF linear regression r = −0.31; p < 1 × 10^−100^). **c,** For 1,740 genes where there exists at least one constrained and one unconstrained transcript, the proportion of expression derived from the constrained transcript is plotted as a histogram.

Finally, we investigated potential differences in LOEUF across human populations, restricting to the same sample size across all populations in order to remove bias due to differential power for variant discovery. As the smallest population in our exome dataset (African/African-American) has only 8,128 individuals, our ability to detect constraint against pLoF variants for individual genes is limited. However, for well-powered genes (expected pLoF >= 10, see Supplementary Information), we observed a lower mean o/e ratio and LOEUF across genes among African/African-American individuals, a population with a larger effective population size, compared to other populations (Extended Data Fig. 8d,e), consistent with the increased efficiency of selection in populations with larger effective population sizes^26,27^.

## Constraint informs disease etiologies

The LOEUF metric can be applied to improve molecular diagnosis and advance our understanding of disease mechanisms. Disease-associated genes, discovered by different technologies over the course of many years across all categories of inheritance and effects, span the entire spectrum of LoF tolerance (Extended Data Fig. 9a). However, in recent years, high-throughput sequencing technologies have enabled identification of highly deleterious variants that are *de novo* or only inherited in small families/trios, leading to the discovery of novel disease genes under extreme constraint against pLoF variation that could not have been identified by linkage approaches that rely on broadly inherited variation (Extended Data Fig. 9b). This result is consistent with a recent analysis which shows a post-WES/WGS era enrichment for gene-disease relationships attributable to *de novo* variants^28^.

Rare variants, which are more likely to be deleterious, are expected to exhibit stronger effects on average in constrained genes (previously shown using pLI from ExAC^29^), with an effect size related to the severity and reproductive fitness of the phenotype. In an independent cohort of 5,305 patients with intellectual disability / developmental disorders and 2,179 controls, the rate of pLoF *de novo* variation in cases is 15-fold higher in genes belonging to the most constrained LOEUF decile, compared to controls (Fig. 5a) with a slightly increased rate (2.9-fold) in the second highest decile but not in others. A similar, but attenuated enrichment (4.4-fold in the most constrained decile) is seen for *de novo* variants in 6,430 patients with autism spectrum disorder (Extended Data Fig. 9c). Further, in burden tests of rare variants (allele count across both cases and controls = 1) of patients with schizophrenia^29^, we find a significantly higher odds ratio in constrained genes (Extended Data Fig. 9d).

**Figure 5.**
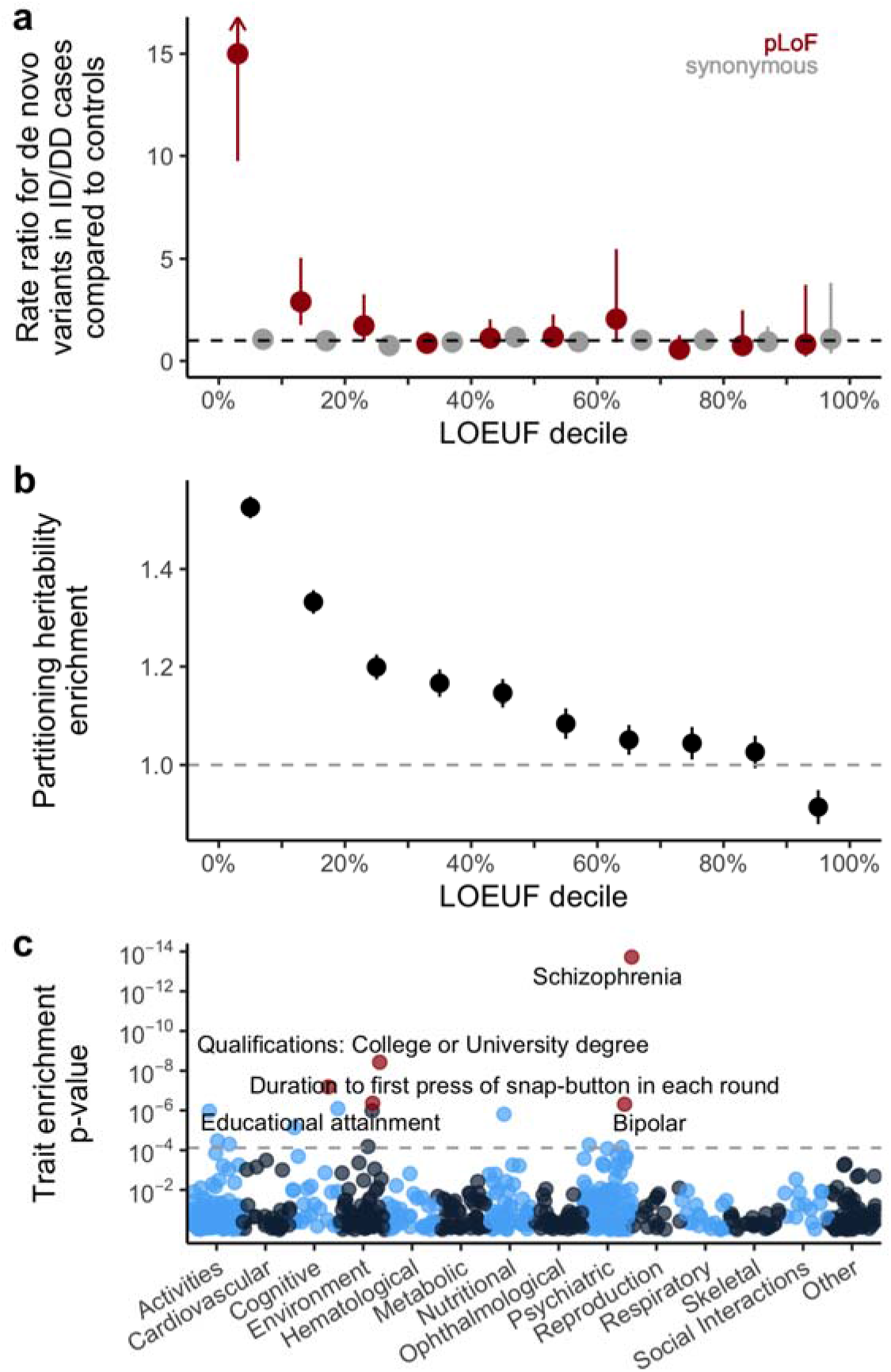
Disease applications of constraint. **a,** The rate ratio is defined by the rate of *de novo* variants (number per patient) in 5,305 intellectual disability / developmental delay (ID/DD) cases divided by the rate in 2,179 controls. pLoF variants in the most constrained decile of the genome are approximately 11-fold more likely to be found in cases compared to controls. Error bars represent 95% confidence intervals. **b,** Marginal enrichment in per-SNV heritability explained by common (MAF > 5%) variants within 100 kb of genes in each LOEUF decile, estimated by LD Score regression. Enrichment is compared to the average SNV genome-wide. The results reported here are from random effects meta-analysis of 276 independent traits (subsetted from the 658 traits with UK Biobank or large-scale consortium GWAS results). Error bars represent 95% confidence intervals. **c,** Conditional enrichment in per-SNV common variant heritability tested using LD score regression in each of 658 common disease and trait GWAS results. P-values evaluate whether per-SNV heritability scales proportional to the LOEUF of the nearest gene, conditional on 75 existing functional, linkage disequilibrium, and MAF-related genomic annotations. Colors alternate by broad phenotype category.

Finally, although pLoF variants are predominantly rare, other more common variation in constrained genes may also be deleterious, including the effects of other coding or regulatory variants. In a heritability partitioning analysis of association results for 658 traits in the UK Biobank and other large-scale GWAS efforts, we find an enrichment of common variant associations near genes that is linearly related to LOEUF decile across numerous traits (Fig. 5b). Schizophrenia and educational attainment are the most enriched traits (Fig. 5c), consistent with previous observations in associations between rare pLoF variants and these phenotypes^30–32^. This enrichment persists even when accounting for gene size, expression in GTEx brain samples, and previously tested annotations of functional regions and evolutionary conservation, and suggests that some heritable polygenic diseases and traits, particularly cognitive/psychiatric ones, have an underlying genetic architecture driven substantially by constrained genes (Extended Data Fig. 10).

## Discussion

In this paper and accompanying publications, we present the largest catalogue of harmonized variant data from any species to date, incorporating exome or genome sequence data from over 140,000 humans. The gnomAD dataset of over 270 million variants is publicly available (https://gnomad.broadinstitute.org), and has already been widely used as a resource for allele frequency estimates in the context of rare disease diagnosis (for a recent review, see Eilbeck *et al*. ^33^), improving power for disease gene discovery ^34–36^, estimating genetic disease frequencies^37,38^, and exploring the biological impact of genetic variation^39,40^. In this manuscript, we describe the application of this dataset to calculate a continuous metric describing a spectrum of tolerance to pLoF variation for each protein-coding gene in the human genome. We validate this method using known gene sets and model organism data, and explore the value of this metric for investigating human gene function and disease gene discovery.

We have focused on high-confidence, high-impact pLoF variants, calibrating our analysis to be highly specific to compensate for the increased false-positive rate among deleterious variants. However, some additional error modes may still exist, and indeed, several recent experiments have proposed uncharacterized NMD-escape mechanisms^41,42^. Further, such a stringent approach will remove some true positives. For example, terminal truncations that are removed by LOFTEE may still exert a LoF mechanism through the removal of critical C-terminal domains, despite the gene’s escape from nonsense mediated decay. Additionally, current annotation tools are incapable of detecting all classes of LoF variation and typically miss, for instance, missense variants that inactivate specific gene functions, as well as high-impact variants regulatory regions. Future work will benefit from the increasing availability of high-throughput experimental assays that can assess the functional impact of all possible coding variants in a target gene^43^, although scaling these experimental assays to all protein-coding genes represents an enormous challenge. Identifying constraint in individual regulatory elements outside coding regions will be even more challenging, and require much larger sample sizes of whole genomes as well as improved functional annotation^44^. We discuss one class of high-impact regulatory variants in a companion manuscript^17^, but many remain to be fully characterized.

While the gnomAD dataset is of unprecedented scale, it has important limitations. At this sample size, we remain far from saturating all possible pLoF variants in the human exome; even at the most mutable sites in the genome (methylated CpG dinucleotides) we observe only half of all possible stop-gained variants. A substantial fraction of the remaining variants are likely to be heterozygous lethal, while others will exhibit an intermediate selection coefficient; much larger sample sizes (in the millions to hundreds of millions of individuals) will be required for comprehensive characterization of selection against all individual LoF variants in the human genome. Such future studies would also benefit substantially from increased ancestral diversity beyond the European-centric sampling of many current studies, which would provide opportunities to observe very rare and population-specific variation, as well as increase power to explore population differences in gene constraint. In particular, current reference databases including gnomAD have a near-complete absence of representation from the Middle East, central and southeast Asia, Oceania, and the vast majority of the African continent^45^, that must be addressed if we are to fully understand the distribution and impact of human genetic variation.

It is also important to understand the practical and evolutionary interpretation of pLoF constraint. In particular, it should be noted that these metrics primarily identify genes undergoing selection against heterozygous variation, rather than strong constraint against homozygous variation^46^. In addition, the power of the LOEUF metric is affected by gene length, with ~30% of the coding genes in the genome still insufficiently powered for detection of constraint even at the scale of gnomAD (Fig. 2d). Substantially larger sample sizes and careful analysis of individuals enriched for homozygous pLoFs (see below) will be useful for distinguishing these possibilities. Further, selection is largely blind to phenotypes emerging after reproductive age, and thus genes with phenotypes that manifest later in life, even if severe or fatal, may exhibit much weaker intolerance to inactivation. Despite these caveats, our results demonstrate that pLoF constraint segments protein-coding genes in a way that correlates usefully with their probability of disease impact and other biological properties, confirming the value of constraint in prioritizing candidate genes in studies of both rare and common disease.

Examples such as *PCSK9* demonstrate the value of human pLoF variants for identifying and validating targets for therapeutic intervention across a wide range of human diseases. As we discuss in more detail in an accompanying manuscript^12^, careful attention must be paid to a variety of complicating factors when using pLoF constraint to assess candidates. More valuable information comes from directly exploring the phenotypic impact of LoF variants on carrier humans, both through “forward genetics” approaches such as Mendelian disease gene mapping, as well as “reverse genetics” approaches leveraging large collections of sequenced humans to find and clinically characterize individuals with disruptive mutations in specific genes. While clinical data are currently available for only a small subset of gnomAD individuals, future efforts integrating sequencing and deep phenotyping of large biobanks will provide valuable insight into the biological implications of partial disruption of specific genes. This is illustrated in a companion manuscript exploring the clinical correlates of heterozygous pLoF variants in the *LRRK2* gene, demonstrating that life-long partial inactivation of this gene is likely to be safe in humans^13^.

Such examples, and the sheer scale of pLoF discovery in this dataset, suggest the near-future feasibility and considerable value of a “human knockout project” - a systematic attempt to discover the phenotypic consequences of functionally disruptive mutations, in either the heterozygous or homozygous state, for all human protein-coding genes. Such an approach will require cohorts of millions of sequenced and deeply, consistently phenotyped individuals and, for the discovery of “complete knockouts”, would benefit substantially from the targeted inclusion of large numbers of samples from populations that have either experienced strong demographic bottlenecks, or high levels of recent parental relatedness (consanguinity)^12^. Such a resource would allow the construction of a comprehensive map directly linking gene-disrupting variation to human biology.

## Acknowledgments

We would like to thank the many individuals whose sequence data are aggregated in gnomAD for their contributions to research, and the users of gnomAD for their helpful and collaborative feedback. We would also like to thank David Altshuler for his important contributions to the development of the gnomAD resource, and Alicia Martin, Eric Fauman, Jon Bloom, Daniel King, and the Hail team for helpful discussions. The results published here are in part based upon data: 1) generated by The Cancer Genome Atlas managed by the NCI and NHGRI (accession: phs000178.v10.p8). Information about TCGA can be found at http://cancergenome.nih.gov, 2) generated by the Genotype-Tissue Expression Project (GTEx) managed by the NIH Common Fund and NHGRI (accession: phs000424.v7.p2), 3) generated by the Exome Sequencing Project, managed by NHLBI, 4) generated by the Alzheimer’s Disease Sequencing Project (ADSP), managed by the NIA and NHGRI (accession: phs000572.v7.p4). KJK was supported by NIGMS F32 GM115208. LCF was supported by the Swiss National Science Foundation (Advanced Postdoc.Mobility 177853). JXC was supported by NHGRI and NHLBI grants UM1 HG006493 and U24 HG008956. Analysis of the Genome Aggregation Database was funded by NIDDK U54 DK105566, NHGRI UM1 HG008900, BioMarin Pharmaceutical Inc., and Sanofi Genzyme Inc. Development of LOFTEE was funded by NIGMS R01 GM104371. The complete acknowledgments can be found in Supplementary Information. We have complied with all relevant ethical regulations.

## Data availability

The gnomAD 2.1.1 dataset is available for download at http://gnomad.broadinstitute.org, where we have developed a browser for the dataset and provide files with detailed frequency and annotation information for each variant. There are no restrictions on the aggregate data released.

## Code availability

All code to perform quality control is provided at https://github.com/broadinstitute/gnomad_qc, and the code to perform all analyses and regenerate all the figures in this manuscript is provided at https://github.com/macarthur-lab/gnomad_lof. LOFTEE is available at https://github.com/konradjk/loftee. All code and software to reproduce figures are available in a Docker image at konradjk/gnomad_lof_paper:0.2.

## Author Contributions

K.J.K, L.C.F, G.T, B.B.C, J.A, Q.W, R.L.C, K.M.L, A.G, M.S, D.R, M.S-B, B.M.N, M.J.D, D.G.M contributed to the writing of the manuscript and generation of figures. K.J.K, L.C.F, G.T, B.B.C, Q.W, R.L.C, K.M.L, A.G, H.B, D.R, M.S-B, E.M.E, E.G.S, J.A.K, N.W, J.X.C, K.E.S, E.P-H, Z.Z, A.H.O’D-L, M.E.T, B.M.N, M.J.D, D.G.M contributed to the analysis of data. K.J.K, L.C.F, G.T, B.B.C, K.M.L, D.P.B, L.D.G, M.S, N.A.W, R.K.W, K.T, Y.F, E.B, T.P, A.W, C.S, K.E.S, Z.Z, A.H.O’D-L, C.V, B.M.N, M.J.D, D.G.M developed tools and methods that enabled the scientific discoveries herein. K.J.K, L.C.F, G.T, B.B.C, J.A, R.L.C, K.M.L, L.D.G, Y.F, E.B, A.H.O’D-L, E.V.M, B.W, M.L, J.S.W, C.V, I.M.A, L.B, K.C, K.M.C, M.C, S.D, S.F, S.G, J.G, N.G, T.J, D.K, C.L, R.M, S.N, N.P, D.R, V.R-R, A.S, M.S, J.S, K.T, C.T, G.W, M.E.T, B.M.N, M.J.D, D.G.M contributed to the production and quality control of the gnomAD data set. All authors listed under The Genome Aggregation Database Consortium contributed to the generation of the primary data incorporated into the gnomAD resource. All authors reviewed the manuscript.

## Extended Data Figures

**Extended Data Figure 1.**
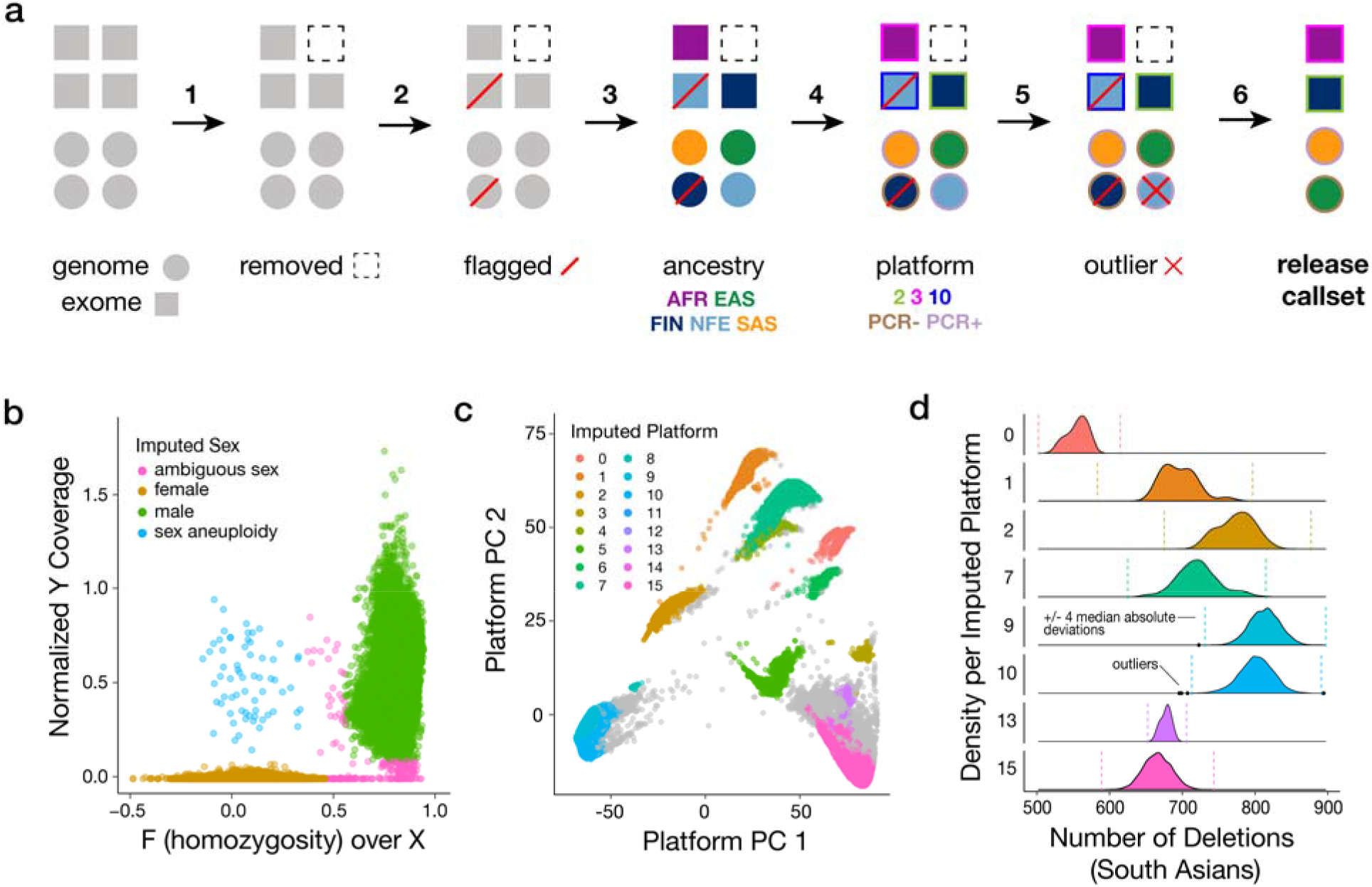
Overview of the sample QC workflow. **a,** Exome (square) and genome (circle) samples underwent quality control as described in detail in the Supplementary Information in the following stages: hard filtering (step 1), relatedness inference (step 2), ancestry inference (step 3), platform inference (step 4, for exomes only), and population- and platform-specific outlier filtering (step 5). Except for samples failing hard filters (dotted outline), all quality control analyses were applied to all samples, regardless of the presence/absence of other QC flags (e.g., relatedness, lack of release permissions, or outlier status; red diagonal bar). Assignment of ancestry labels is represented by fill color and accompanying three-letter ancestry group abbreviation. Assignment of platform labels is represented by outline color and a numbered label for exomes (corresponding to imputed platforms) and a PCR +/− label for genomes. The final set of samples included in the gnomAD release (125,748 exomes and 15,708 genomes) was defined to be the set of unrelated samples with release permissions, no hard filter flags, and no population- and platform-specific outlier metrics (step 6). **b,** In exomes, the chromosomal sex of samples was inferred based on the inbreeding coefficient on chromosome X and the coverage of chromosome Y into male (green), female (amber), ambiguous sex (pink), and sex chromosome aneuploid (blue). **c,** The top two principal components from PCA-HDBSCAN analysis of exome capture regions. Sequencing platforms were inferred for exome samples based on PCA analysis of biallelic variant call rates over all known exome capture regions, and samples were assigned a cluster label (0-15, or unknown) using HDBSCAN. **d,** We performed platform- and population-specific outlier filtering for a number of quality control metrics: here, we show the distribution of the number of deletions in South Asians across platforms. Distributions (and accordingly, median and median absolute deviations, MAD) for these metrics varied widely both by population and sequencing platform (numbered on y-axis). Outliers (black dots) were defined as samples with values outside four MAD (shown by dotted vertical lines) from the median of a given metric.

**Extended Data Figure 2.**
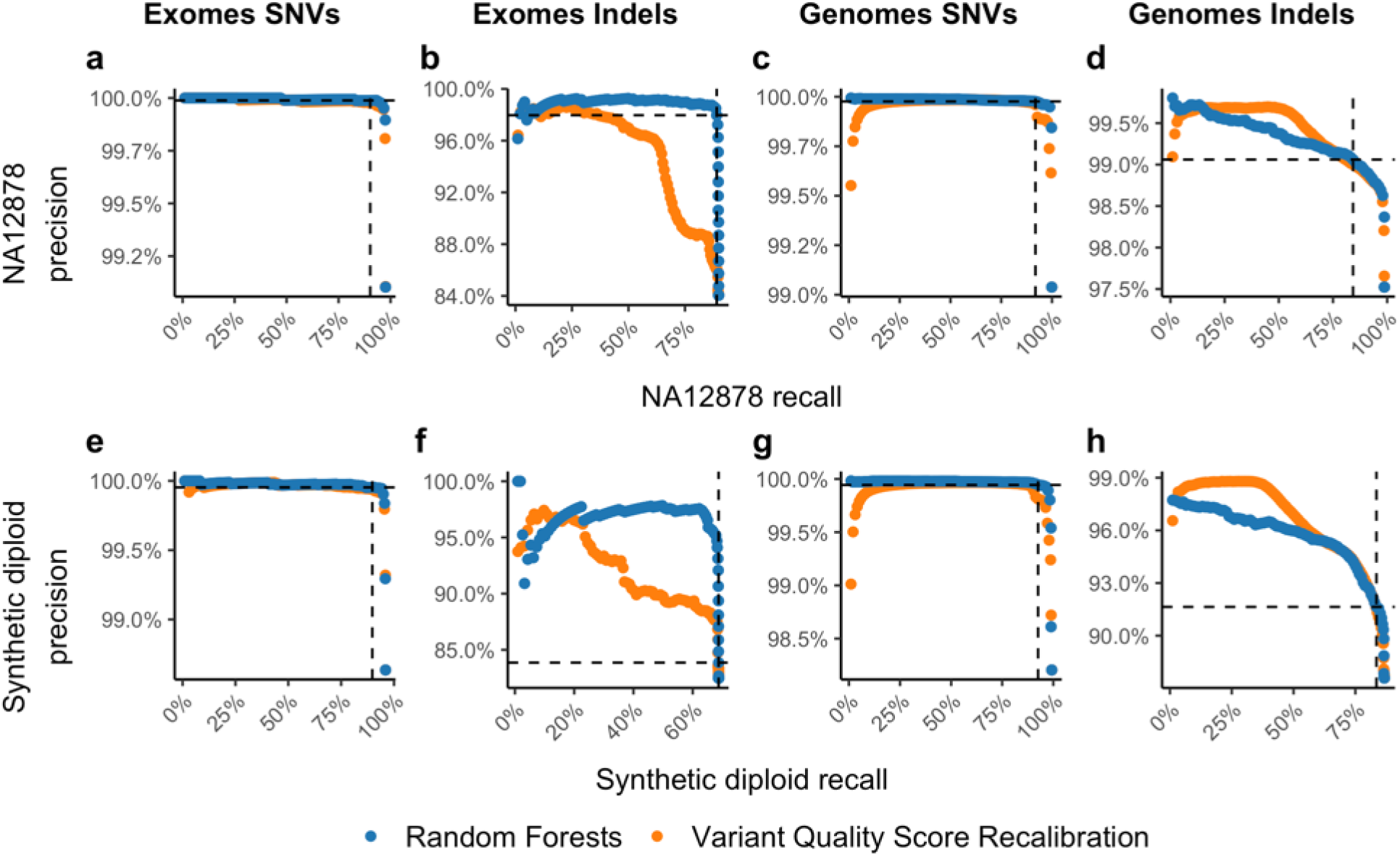
Variant calling performance for common variants. Precision-recall curves are shown for variant calls in two samples with independent gold-standard data, NA12878^47^ (**a-d**) and a synthetic diploid mixture^48^ (**e-h**). The random forest (RF, blue) approach described here is compared to the current state-of-the-art GATK Variant Quality Score Recalibration (VQSR, orange) for exome SNVs (**a, e**) and indels (**b, f**), and genome SNVs (**c, g**) and indels (**d, h**). Note that the indels presented in (**f**) and (**h**) exclude 1bp indels as they aren’t well characterized in the synthetic diploid mixture gold standard sample. In all cases, at the thresholds chosen (dashed lines representing 10% and 20% of SNVs and indels filtered, respectively), RF outperforms or is similar to VQSR.

**Extended Data Figure 3.**
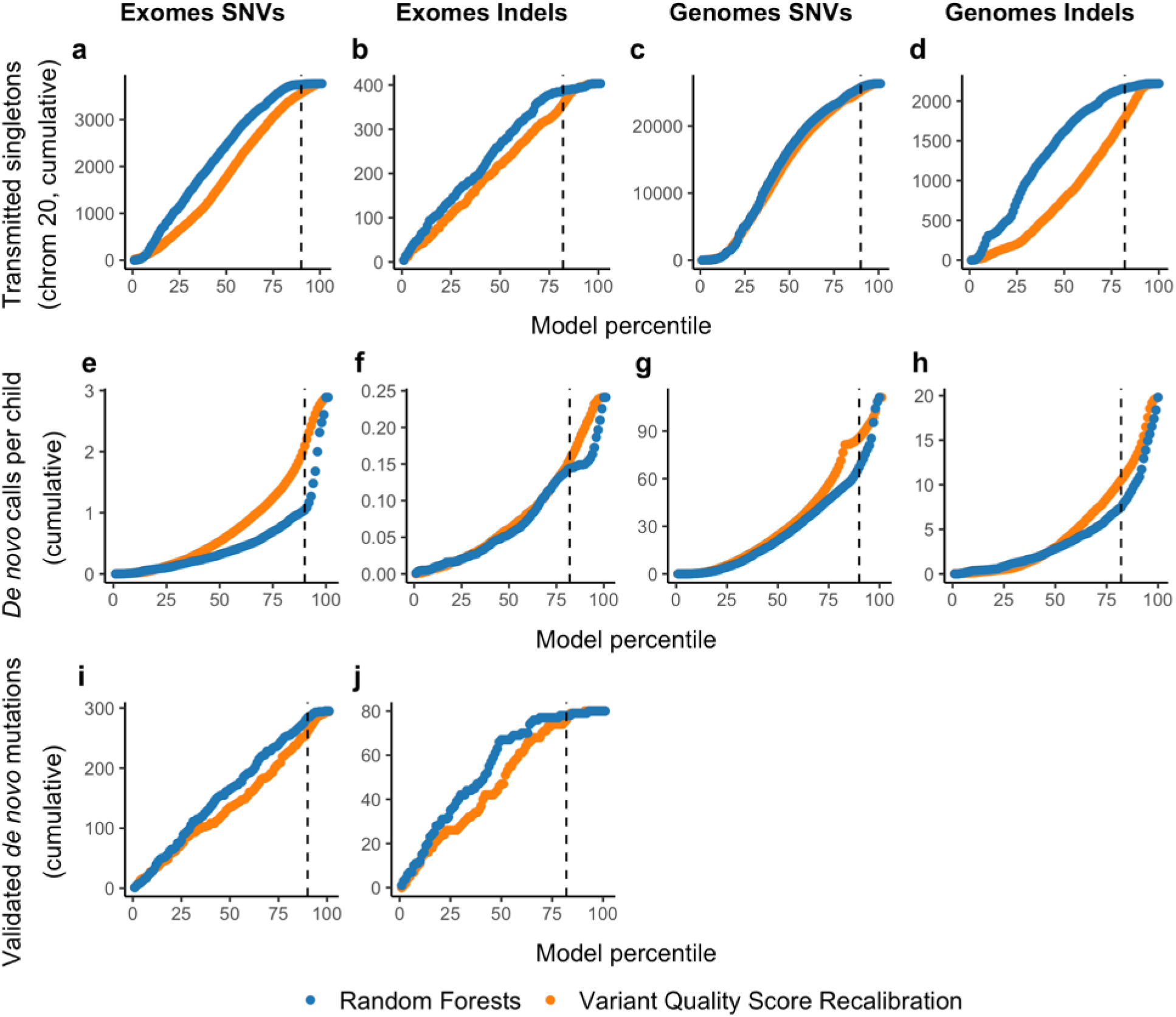
Variant calling performance for rare variants. For each of the curves in this figure, the x-axis shows the cumulative ranked percentile for our random forest (RF, blue) model and, as a comparison, for the current state-of-the-art GATK Variant Quality Score Recalibration (VQSR, orange). That is, the point at 10 shows the performance of the 10% best scored data, the point at 50, shows the performance 50% best-scored data, etc. **a-d**, The number of transmitted singletons (singletons in the unrelated individuals that are transmitted to an offspring) on chromosome 20 for exome SNVs (**a**) and indels (**b**), and genome SNVs (**c**) and indels (**d**). Chromosome 20 was not used for training our random forest model. We expect most of these to be real variants since we observe Mendelian transmission of an allele that was sequenced independently in a parent and child. **e-h,** The number of bi-allelic *de novo* calls per child (4,568 exomes, 212 genomes) outside of low-complexity regions. The expectation is that there is ~1.6 *de novo* SNV (**e**) and ~0.1 *de novo* indels per exome (**f**), and ~65 *de novo* SNVs (**g**) and ~5 *de novo* indels (**h**) per genome^21^. **i-j**, The number of independently validated *de novo* mutations, available for a subset of 331 exome samples for which *de novo* mutations were validated as part of other studies^49^. In all cases, at the thresholds chosen (dashed lines representing 10% and 20% of SNVs and indels filtered, respectively), RF outperforms or is similar to VQSR.

**Extended Data Figure 4.**
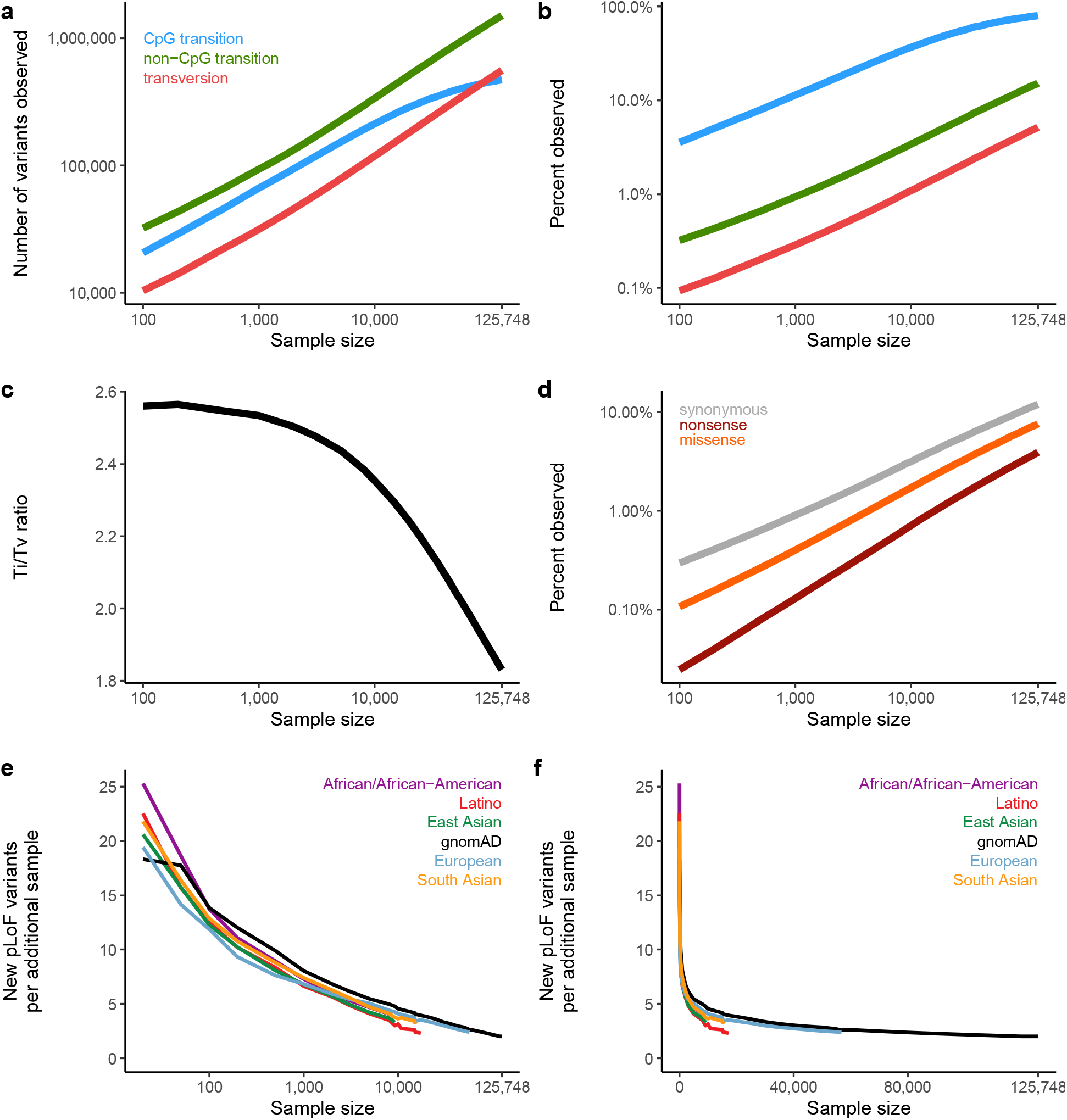
Variant discovery at large sample sizes. **a-b,**The total number of variants observed (**a**) and the proportion of possible variants observed (**b**) as a function of sample size, broken down by variant class. At large sample sizes, CpG transitions become saturated, as previously described^4^. Colours are consistent in **a-b**. **c,** This results in a decrease of the transition:transversion (Ti/Tv) ratio. **d,** When broken down by functional class, we observe the effects of selection, where synonymous variants have the highest proportion observed, followed by missense and pLoF variants. **e-f,** The number of additional pLoF variants introduced into the cohort as a function of sample size on a log (**e**) and linear (**f**) scale. Here, gnomAD (black) refers to a uniform sampling from the population distribution of the full cohort of exome equenced individuals.

**Extended Data Figure 5.**
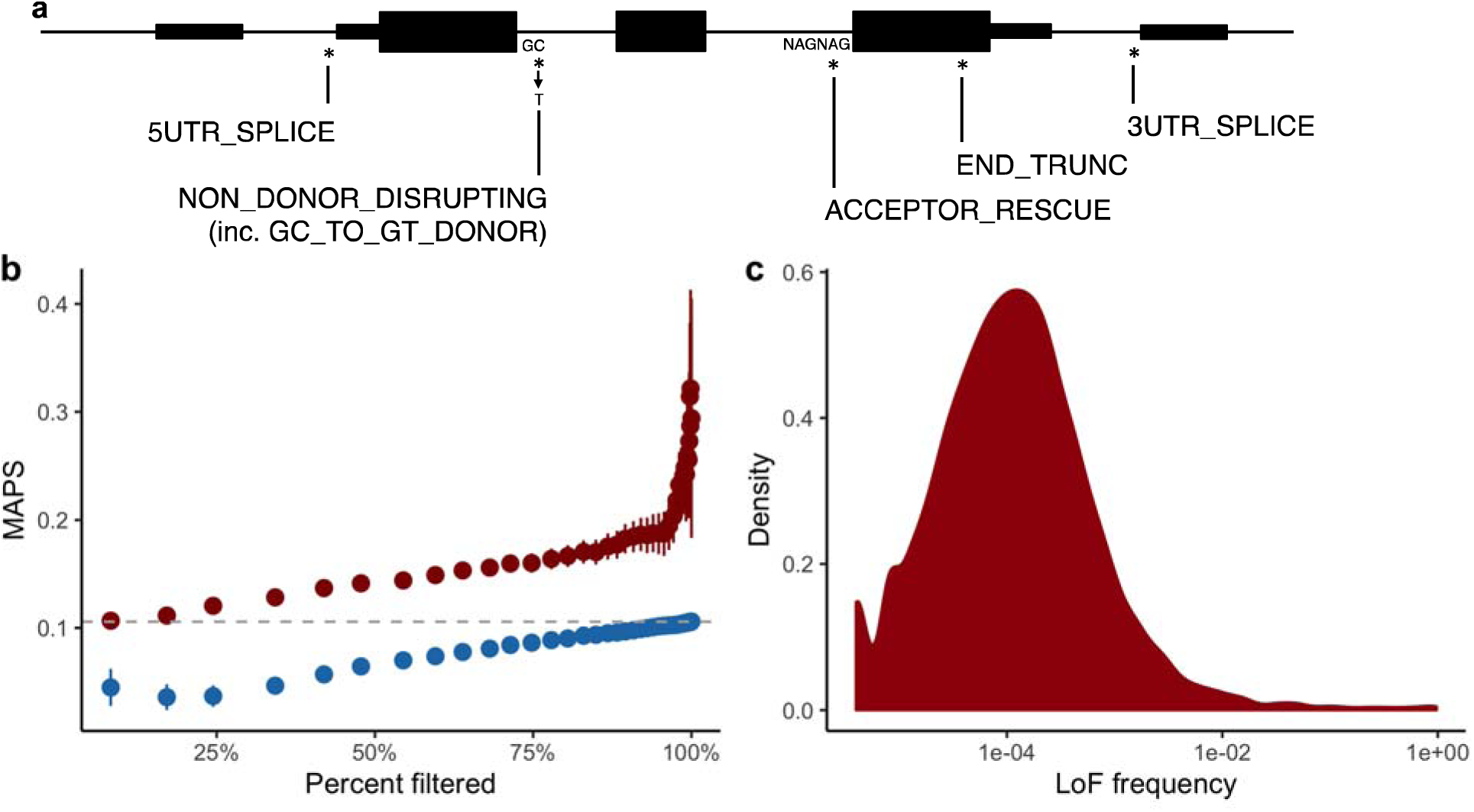
Using LOFTEE to create a high confidence set of pLoF variation. **a,** Schematic of LOFTEE filters. LOFTEE filters out putative stop-gained, essential splice, and frameshift variants based on sequence and transcript context, as well as flagging exonic features such as conservation (not shown). For instance, variants that are not predicted to disrupt splicing based on retention of a strong splice site, or rescue of a nearby splice site. Additional filters not shown include: ANC_ALLELE (the alternate allele is the ancestral allele), NON_ACCEPTOR_DISRUPTING and DONOR_RESCUE (opposite to those already shown). **b,** In order to tune the END_TRUNC filter, we retained variants that pass the 50 base pair rule (are more than 50 bp before the 3’-most splice site). The overall MAPS score for variants that fail this rule is shown in gray. For the remaining 39,072 variants, we computed the sum of the GERP score of bases deleted by the variant. At 40 bins of this score, we compute the MAPS score for those variants retained at this threshold (red) compared to variants removed at this threshold (blue), and plot this as a function of the proportion of variants filtered at this threshold. We chose the 50% point as it retains variants with a MAPS score of 0.14, while removing variants with a MAPS score of 0.06. Error bars represent 95% confidence intervals. **c,** Density plot of aggregate pLoF frequency computed from high-confidence pLoF variants discovered using LOFTEE.

**Extended Data Figure 6.**
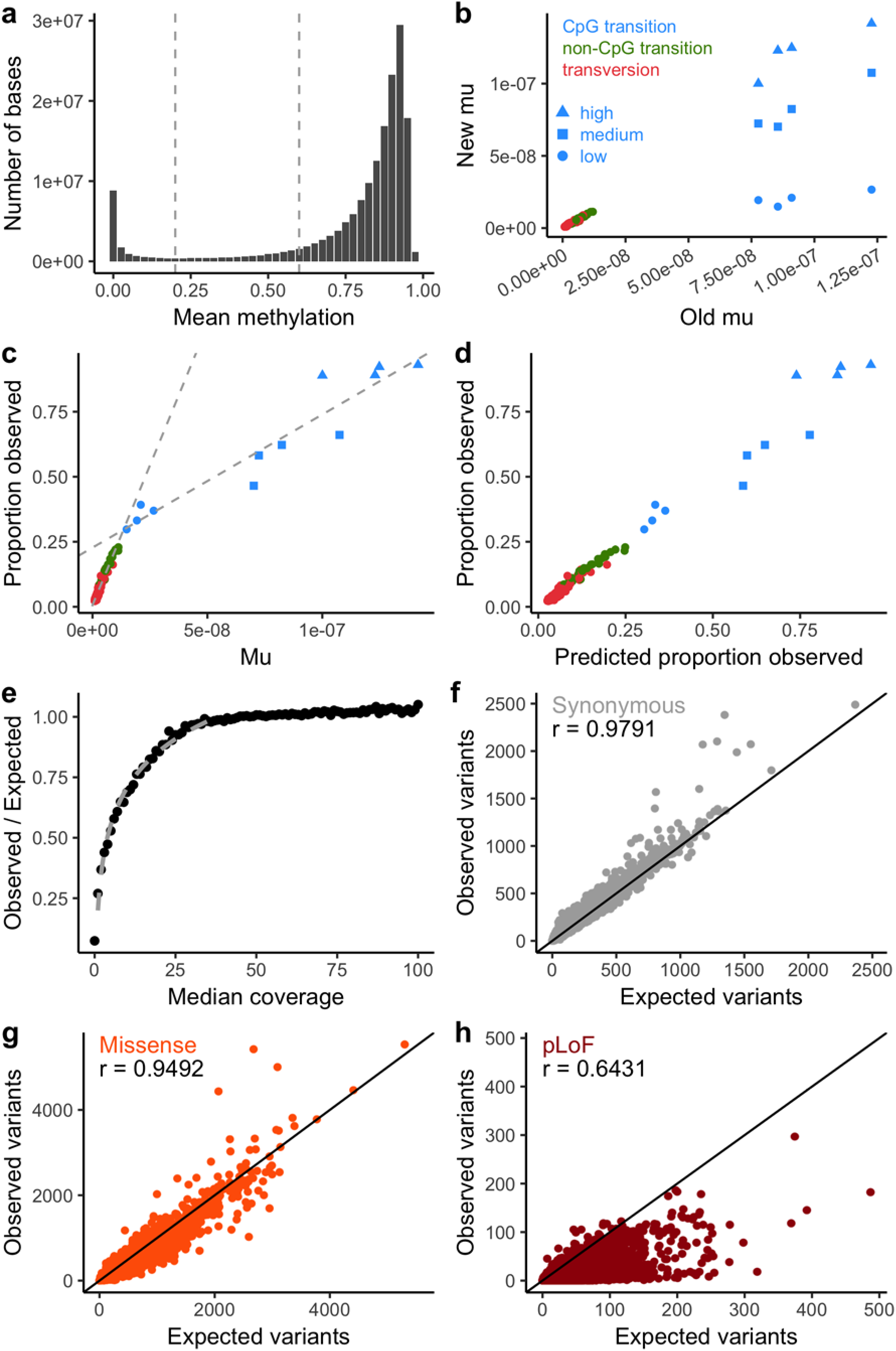
Computing the depletion of variation of functional categories. **a,** The distribution of mean methylation values across 37 tissues and across every CpG dinucleotide in the genome. We divided the genome into 3 levels (low methylation, missing or < 0.2; medium, 0.2-0.6; and high, > 0.6) and computed all ensuing metrics based on these categories. **b,** Comparison of estimates of the mutation rate with previous estimates^50^. For transversions and non-CpG transitions, we observe a strong correlation (linear regression r = 0.98; p = 2.6 × 10^−65^). For CpG transitions, the new estimates are calculated separately for the 3 levels of methylation and track with these levels. Colors and shapes are consistent in **b-d**. **c,** For panels (**c-e**), only synonymous variants are considered. The proportion of possible variants observed for each context is correlated with the mutation rate. We compute two fit lines, one for CpG transitions, and one for other contexts to calibrate our estimates. **d,** Calibration of each context to compute a predicted proportion observed after fitting the two models in (**c**), which is used to calculate an expected number of variants at high coverage. **e,** With an expectation computed from high coverage regions, the observed/expected ratio follows a logarithmic trend with the median coverage below 40X, which is used to correct low coverage bases in the final expectation model. **f-h,** For each transcript, the observed number of variants is plotted against the expected number from the model described above, for synonymous (**f**), missense (**g**), and pLoF (**h**) variants, and the linear regression coefficient is shown. Note that the expectation does not include selection, and so, pLoF and, to a lesser extent, missense variants exhibit lower observed values than expected.

**Extended Data Figure 7.**
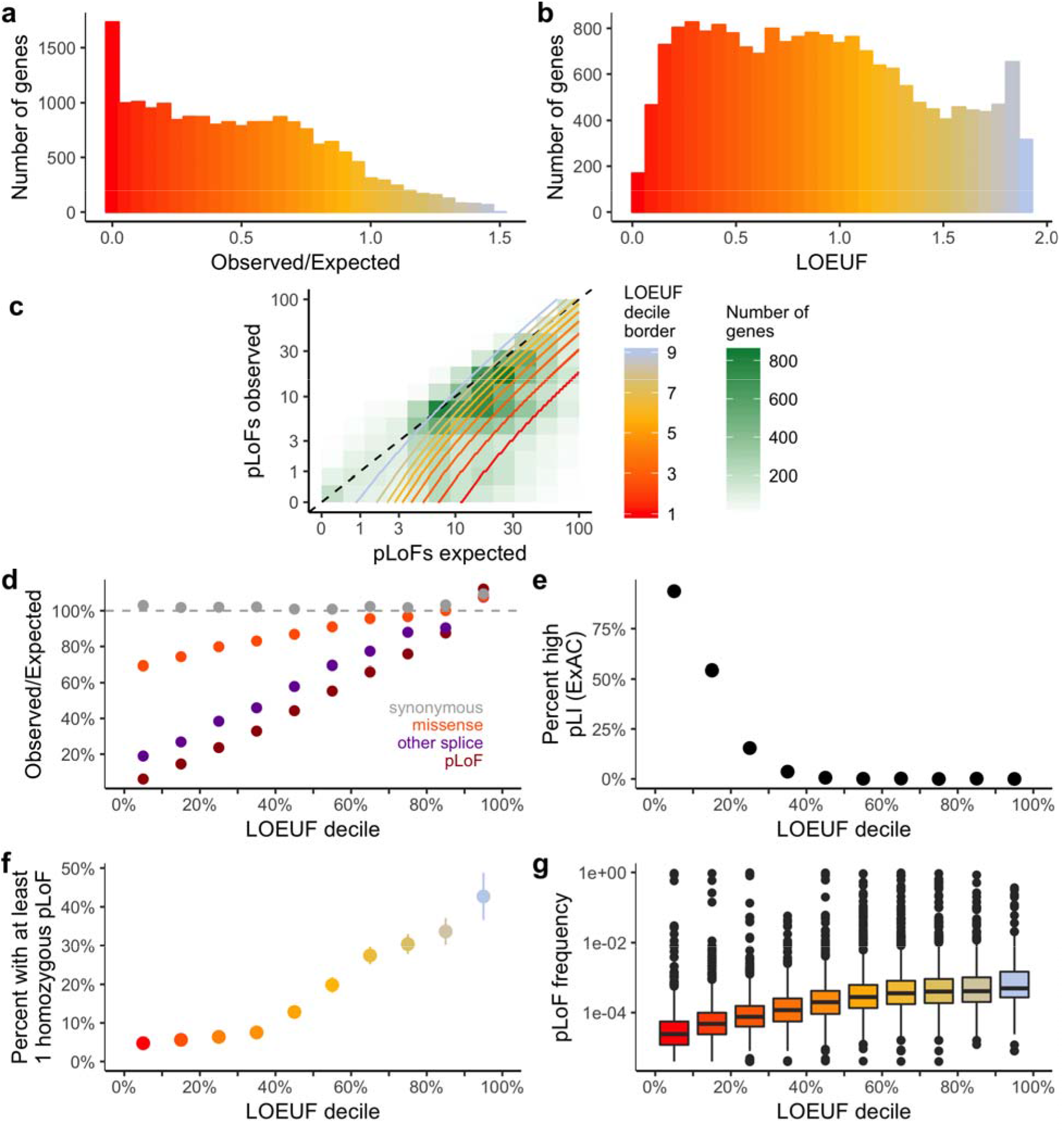
Genomic properties of constrained genes. **a-b,** Histogram of observed/expected ratio of pLoF variation (**a**) and LOEUF (**b**). Most genes have fewer observed variants than expected (median o/e = 0.48), and the genes with no observed pLoFs are distinguished between confidently constrained genes and noise by LOEUF. **c,** A 2-d density plot of the number of observed vs expected pLoF variants. The boundaries of each decile are plotted as gradients (the most constrained decile is below the lowest red line, etc.). **d,** Observed/expected ratios of various functional classes across genes within each LOEUF decile. The most constrained decile has approximately 6% of the expected pLoFs, while synonymous variants are not depleted and missense variants exhibit modest depletion. **e,** The percentage of each LOEUF decile that was described in ExAC as constrained, or pLI > 0.9^4^. **f,** The percentage of each LOEUF decile that have at least one homozygous pLoF variant. **g,** Boxplots of the aggregate pLoF frequency for each LOEUF decile (center line, median; box limits, upper and lower quartiles; whiskers, 1.5x interquartile range; points, outliers). In d-f, error bars represent 95% confidence intervals (note that in some cases these are fully contained within the plotted point).

**Extended Data Figure 8.**
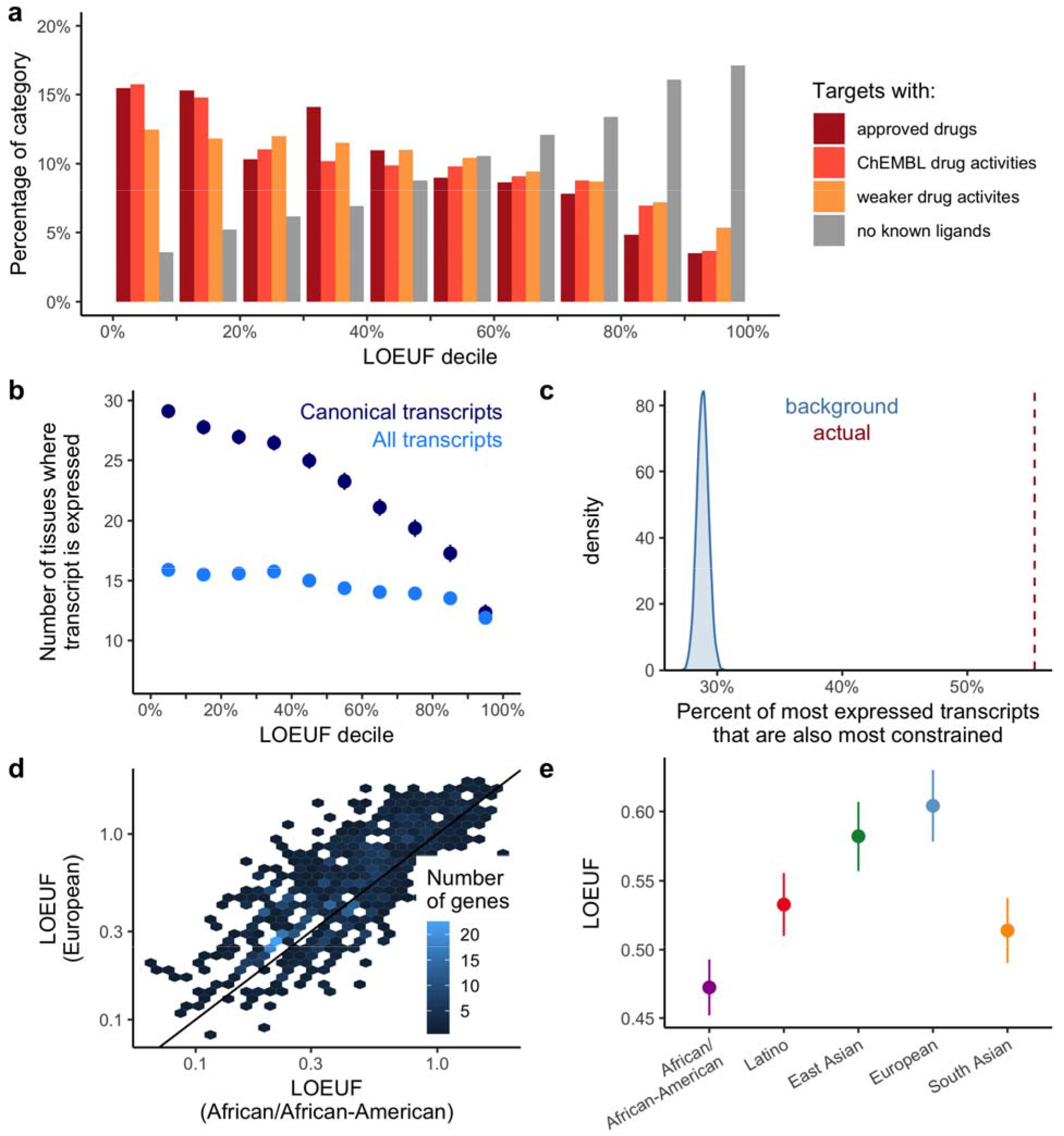
Biological properties of constrained genes. **a,** The percentage of genes in each functional category from Pharos (see Supplementary Information) is broken down by LOEUF decile. **b,** The mean number of tissues where a transcript is expressed, binned by transcript-based LOEUF decile, is shown for all transcripts and canonical transcripts. **c,** The percent of genes where the most expressed transcript is also the most constrained is plotted in red, which is enriched compared to a permuted set (blue). **d,** For 927 genes with expected pLoF >= 10 in both the African/African-American and European population subsets (n = 8,128), the LOEUF scores are highly correlated (linear regression r = 0.78, p < 10^−100^), with a lower mean score observed in the African/African-American population (0.49 vs 0.62; two-sided t-test p = 4.1 × 10^−14^), which has a higher effective population size. **e,** The mean LOEUF score for 865 genes with expected pLoF >= 10 in all populations (n = 8,128). Error bars represent 95% confidence intervals.

**Extended Data Figure 9.**
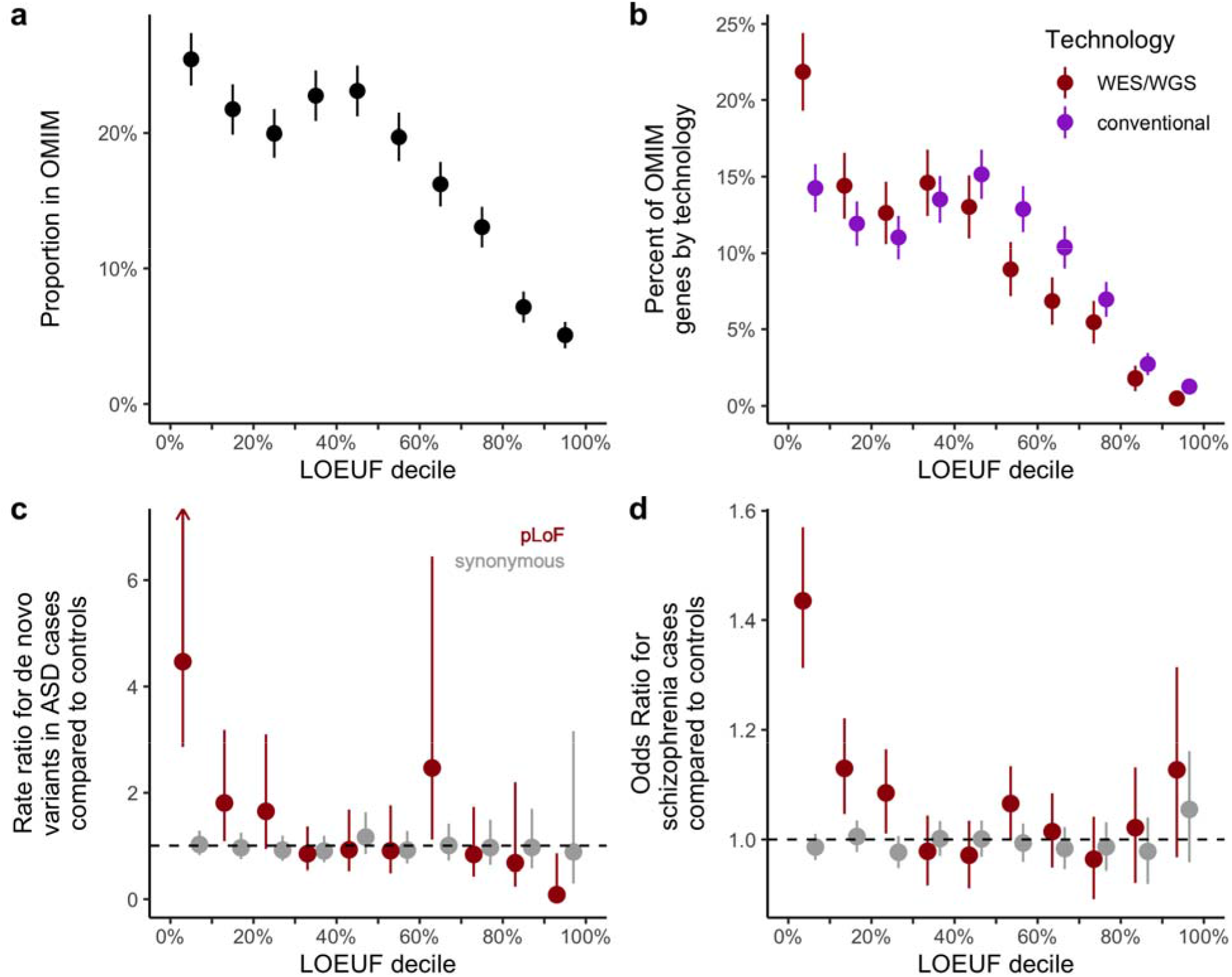
Applications of constraint metrics to rare variant analysis of disease. **a,** Proportion of each LOEUF decile found in OMIM. **b,** Proportion of disease-associated genes discovered by whole exome/genome sequencing (WES/WGS) compared to conventional (typically linkage), plotted by LOEUF decile. The latter are more constrained (LOEUF 0.674 vs 0.806, two-sided t-test p = 1.2 × 10^−16^), suggesting the effectiveness of these techniques picking up genes with a *de novo* mechanism of disease, compared to recessive genes by linkage methods. **c,** Similar to Fig. 5a, the rate ratio is defined by the rate of *de novo* variants (number per patient) in autism cases divided by the rate in controls. pLoF variants in the most constrained decile of the genome are approximately 4-fold more likely to be found in cases compared to controls. **d,** The mean odds ratio of a logistic regression of schizophrenia^29^ is plotted for each LOEUF decile. Error bars in **a-d** correspond to 95% confidence intervals.

**Extended Data Figure 10.**
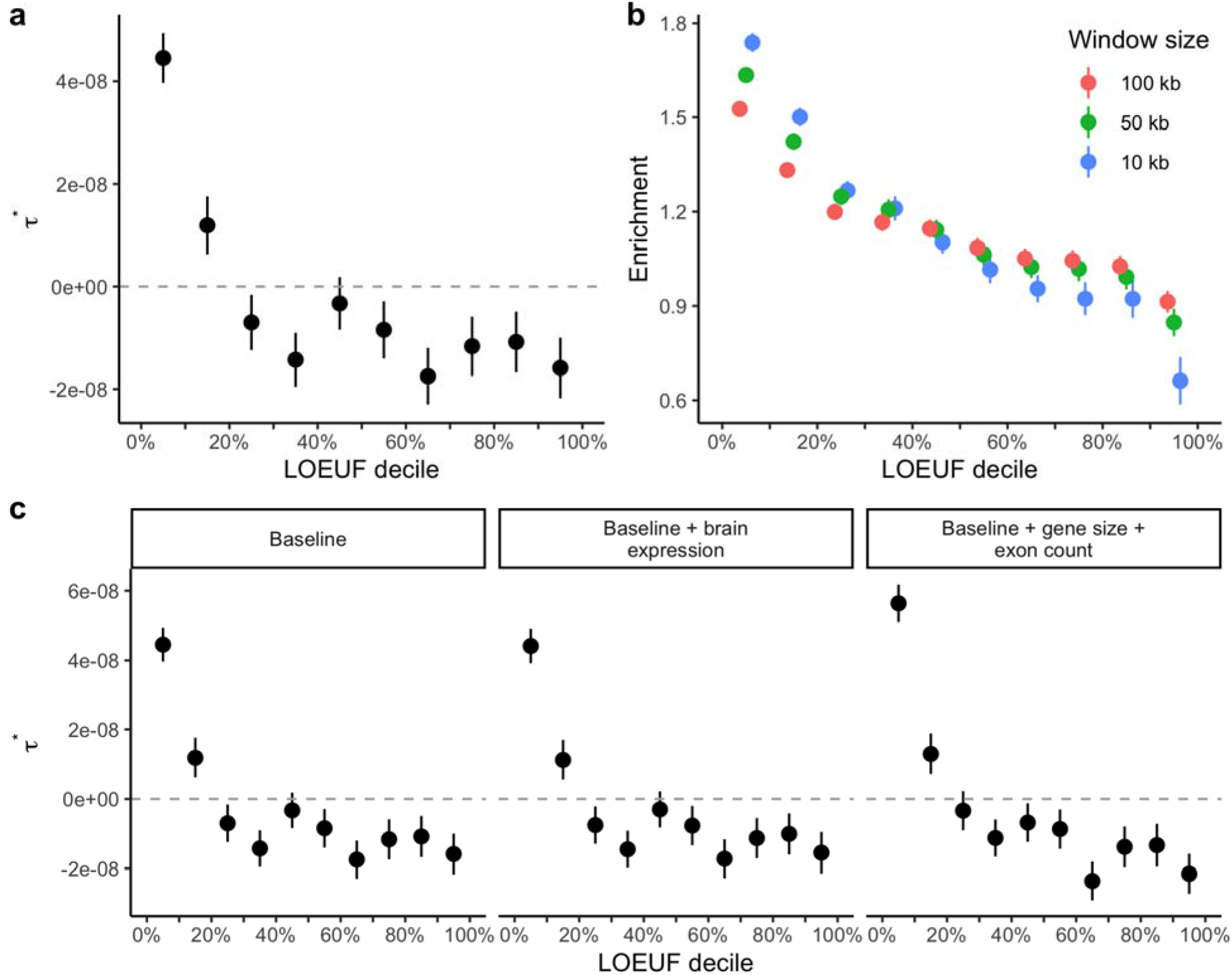
Applications of constraint metrics to common variant analysis of disease. **a,** coefficient for each LOEUF decile across 276 independent traits., unlike the enrichment measure reported in Fig. 5, is adjusted for 74 baseline genomics annotations. Positive values of indicate greater per-SNP heritability than would be expected based on the other annotations in the baseline model, while negative values indicate depleted per-SNP heritability compared to that baseline expectation. **b,** Enrichment coefficient for each LOEUF decile using different window sizes to define which SNPs to include upstream and downstream of each gene. **c,** Enrichment coefficient for each LOEUF decile across traits after controlling for brain expression and gene size. Results are consistent with those shown in Fig. 5 indicating that brain gene expression and gene size do not fully explain the enrichment of heritability observed in constrained genes. Error bars represent 95% confidence intervals.

