## Supplementary Information for "The mutational constraint spectrum quantified from variation in 141,456 humans"

### gnomAD supplement

**gnomAD supplement** 1
**Data processing** 4
Alignment and read processing 4
Variant Calling 4
Coverage information 5
Data processing 6
**Sample QC** 7
Hard filters 7
Supplementary Table 1 | Sample counts before and after hard and release filters 9
Supplementary Table 2 | Counts by data type and hard filter 9
Platform imputation for exomes 9
Supplementary Table 3 | Exome platform assignments 10
Supplementary Table 4 | Confusion matrix for exome samples with known platform labels 11
Relatedness filters 11
Supplementary Table 5 | Pair counts by degree of relatedness 12
Supplementary Table 6 | Sample counts by relatedness status 13
Population and subpopulation inference 13
Supplementary Figure 1 | Continental ancestry principal components. 14
Supplementary Table 7 | Population and subpopulation counts 16
Population- and platform-specific filters 16
Supplementary Table 8 | Summary of outliers per population and platform grouping 17
Finalizing samples in the gnomAD v2.1 release 18
Supplementary Table 9 | Sample counts by filtering stage 18
Supplementary Table 10 | Sample counts for genomes and exomes in gnomAD subsets 19
**Variant QC** 20
Hard filters 20
Random Forest model 20
Features 21
Supplementary Table 11 | Features used in final random forest model 21
Training 22
Supplementary Table 12 | Random forest training examples 22
Evaluation and threshold selection 22
Final variant counts 24
Supplementary Table 13 | Variant counts by filtering status 25
Comparison of whole-exome and whole-genome coverage in coding regions 25
**Variant annotation** 30
Frequency and context annotation 30

Functional annotation 31
Supplementary Table 14 | Variants observed by category in 125,748 exomes 33
Supplementary Figure 5 | Percent observed by methylation. 34
LOFTEE 34
Supplementary Table 15 | pLoF variants discovered in gnomAD 37
Genes affected by clonal hematopoiesis 37
Supplementary Table 16 | Genes with evidence of CHIP 39
Aggregate pLoF frequency 40
Homozygous variant curation 41
Supplementary Table 17 | pLoF alleles per individual. 43
**Constraint modeling 44**
Mutational model 44
Improvements to constraint model 45
Summary of constraint metrics 47
Supplementary Figure 7 | LOEUF summaries. 48
Supplementary Figure 8 | The sample size required for well-powered constraint calculations. 50
**Constraint assessment and implications 51**
Gene list comparisons 51
Supplementary Table 18 | Gene list membership by LOEUF decile 51
Supplementary Figure 9 (previous page) | Genes within gene lists by LOEUF decile. 53
Structural variant comparisons 53
Mouse and cell model comparisons 54
Supplementary Table 19 | Mammalian Phenotype Term lists 55
Supplementary Table 20 | Comparison of genes we observe homozygous deletion in gnomAD population with other gene lists. 57
Functional categorization 57
Network analysis 58
Expression 58
Population-specific constraint modeling 60
Comparison to previous metrics of essentiality 60
Supplementary Figure 10 | Comparison to other gene essentiality metrics. 62
Performance as a function of sample size 62
Supplementary Figure 11 | Performance of LOEUF by sample size. 62
**Disease analysis 63**
Rare disease 63
Common disease 64
Supplementary Table 21 | Phenotypes with association between heritability and constraint. 70
**Data Availability 71**
Release files 71

Code availability 71
The gnomAD Browser 72
Supplementary Datasets 73
**Acknowledgments** **78**
**References** **81**

### Data processing

Laura D. Gauthier, Konrad J. Karczewski, Ryan L. Collins, Kristen M. Laricchia, Yossi Farjoun, Laurent C. Francioli, Eric Banks, Daniel G. MacArthur

##### *Alignment and read processing*

To create a comprehensive reference panel, we integrated whole exome and genome sequence data acquired from many sources, sequenced across many labs with a variety of library prep methods (including various exome capture platforms, and for genomes, both PCR+ and PCR-) over more than five years. This study was overseen by the Broad Institute's Office of Research Subject Protection and the Partners Human Research Committee, and was given a determination of Not Human Subjects Research. Informed consent was obtained from all participants.

For data sequenced externally, we first imported FASTQ- or BAM-level data from all sources for inclusion in our pipeline, based on the Picard suite of software tools version 1.1431, as described previously[4], with any differences noted below. We mapped reads onto the human genome build 37 using bwa aln for exomes[4] and bwa mem version 0.7.7[50] for genomes. The FASTA file can be found at ftp.ncbi.nlm.nih.gov/sra/reports/Assembly/GRCh37-HG19_Broad_variant/Homo_sapiens_assembly19.fasta, which has 85 contigs including a decoy (NC_007605, 171823bp).

##### *Variant Calling*

Variants were jointly called using the Genome Analysis Toolkit (GATK) Best Practices for germline SNVs and indels[51]. Briefly, samples were called individually using local realignment by HaplotypeCaller (version nightly-2015-07-31-g3c929b0) in GVCF mode, such that every position in the genome is assigned likelihoods for discovered variants or for the reference. These per-sample GVCF genotype data, alleles, and sequence-based annotations were then merged using GenomicsDB (https://github.com/Intel-HLS/GenomicsDB), a datastore designed for genomics that takes advantage of a cohort's sparsity of variant genotypes. Samples were jointly genotyped for high confidence alleles using GenotypeGVCFs version 3.4-89-ge494930, on all autosomes and the X chromosome for genomes, and on all autosomes, as well as X and Y chromosomes for exomes. Variant call accuracy was estimated using Variant Quality Score Recalibration (VQSR) in GATK (3.6-0-g89b7209), though we have also implemented a Random Forest (RF) approach (see "Variant QC" below). In the process of manuscript revisions, we identified an issue with undercalling of homozygous variants due to low levels of contamination[52], which does not materially affect the analyses in this paper, except as noted.

Structural variants (SV) were called from WGS data (mean coverage: 32X) for a largely overlapping set of 14,891 samples. Methods for SV discovery & genotyping are covered in detail in a companion paper[11], but are also briefly summarized here. We ran an improved version of our multi-algorithm ensemble approach[53] that integrates information across several SV discovery algorithms (Manta v1.0.3, DELLY v0.7.7, MELT v2.0.5, and cn.MOPS v1.20.1) and queries evidence directly from the aligned WGS libraries (read depth, anomalous read pairs, split reads, and SNV B-allele frequencies) to maximize sensitivity for all classes of SV. Following initial SV discovery, we filtered predicted breakpoints with a series of semi-supervised machine learning methods, jointly genotyped passing breakpoints across all samples using a Bayesian consensus approach, and collapsed breakpoints into fully resolved SV alleles with a heuristic-based method. After post hoc quality control, including pruning first-degree relatives and low-quality samples, and restricting to samples overlapping those with this manuscript, we documented a total of 366,412 SV sites across 6,749 unrelated samples.

##### *Coverage information*

Coverage was calculated on a random subset of samples constituting approximately 10% of the full dataset using the depth tool from samtools (v1.4, modified to cap coverage as previously described[4] for efficiency). Metrics were generated separately for genomes and exomes across their respective calling intervals with the base quality threshold set to 10 and the mapping quality threshold set to 20. Sites with zero depth were included and coverage was capped at 100x for a given sample and base-pair.

*Data processing*

To perform quality control and analysis of the sequencing data at scale, we use an open-source, scalable software framework called Hail (https://github.com/hail-is/hail, https://hail.is), which leverages Apache Spark to distribute computational tasks across thousands of nodes to process data at the terabyte scale. We imported the SNV and indel calls into Hail 0.2 as a MatrixTable using hl.import_vcf, and loci were standardized using minimal representation (https://github.com/ericminikel/minimal_representation) using hl.min_rep (https://github.com/macarthur-lab/gnomad_qc/blob/master/load_data/import_vcf.py).

Truth datasets for random forest training (see Training below), as well as gold standard datasets for two samples for variant QC, NA12878[46] and synthetic diploid[47], were loaded using hl.import_vcf. Additionally, we loaded a set of validated *de novo* variants for variant QC, as well as ClinVar (VCF version from 10/28/2018) for assessment of LOFTEE (see Variant annotation below). Methylation data from 37 tissues from the Roadmap Epigenomics Project[54] was loaded and the mean methylation value across these tissues was computed for each base (https://github.com/macarthur-lab/gnomad_qc/blob/master/load_data/import_resources.py).

Raw coverage (per-base, per-individual) files were loaded using hl.import_matrix_table, and summary metrics were computed for each base, including mean coverage, median coverage, and the percent of samples above 1X, 5X, 10X, 15X, 20X, 25X, 30X, 50X, and 100X (https://github.com/macarthur-lab/gnomad_qc/blob/master/load_data/load_coverage.py).

### Sample QC

Grace Tiao, Kristen M. Laricchia, Konrad J. Karczewski, Laurent C. Francioli, Monkol Lek, Daniel G. MacArthur

Except for certain hard filter metrics explicitly noted below, all sample QC computations were performed on the Google Cloud Platform using a Python pipeline composed with the Hail library (https://github.com/hail-is/hail, https://hail.is/), an open-source framework for analyzing large-scale genomic datasets. The pipeline is available in its entirety at https://github.com/macarthur-lab/gnomad_qc and is summarized in Extended Data Fig. 1a, where numbered steps correspond to the following scripts in the code repository:

1. Hard filtering: apply_hard_filters.py
2. Relatedness inference: joint_sample_qc.py
3. Ancestry inference: joint_sample_qc.py, assign_subpops.py
4. Platform inference: exomes_platform_pca.py
5. Population- and platform-specific outlier filtering: joint_sample_qc.py
6. Finalizing release callset: finalize_sample_qc.py

#### *Hard filters*

Sample QC metrics were computed for each sample in the call set over a set of high-confidence variants: bi-allelic, high-call rate (> 0.99), common SNVs (allele frequency > 0.1%). The chromosomal sex of samples was inferred based on the inbreeding coefficient (F) for these common variants on chromosome X and, for exomes, the coverage of chromosome Y normalized to chromosome 20 coverage (Extended Data Fig. 1b). For exomes, samples with F > 0.6 and normalized Y coverage > 0.1 were classified as male, and samples with F < 0.5 and normalized Y coverage < 0.1 were classified as female. Samples with F values falling within standard female ranges that exhibited normalized Y values > 0.1 threshold were classified as sex aneuploid. For genomes, as no chromosome Y variants were called, samples with F > 0.8 were classified as male and samples with F < 0.5 were classified as female. For both exomes and genomes, samples with intermediate F values were classified as ambiguous sex.

Samples were flagged as failing hard filters if they exhibited high contamination (freemix > 0.05, computed using VerifyBamID 1.0.0), low sample level call rates (< 0.895), high rates of chimeric reads (> 0.05, computed using Picard 1.1431), ambiguous sex, sex aneuploidy, low coverage (for exomes, a mean chromosome 20 coverage equal to 0; for genomes, mean depth < 15), and/or low median insert sizes (genomes only, size < 250 bp).

Hematological, somatic, and pediatric cancer samples (age of onset < 30 years) were flagged for removal based on sample barcodes and age metadata from The Cancer Genome Atlas (TCGA). One TCGA sample, TCGA-06-0178-10B-01D-1491-08, was flagged for removal because it is known to have been swapped with a tumor sample. Samples with known severe pediatric disease phenotypes and samples lacking data usage permission for public release were also flagged for removal (Supplementary Tables 1,2).

At this stage, compressed (hardcalls) versions of the exome and genome datasets were created for most downstream analyses. High-quality genotypes were marked if they had depth (DP) >= 10, genotype quality (GQ) >= 20, and minor allele fraction >= 0.2 for all non-reference alleles of heterozygous genotypes. For non-PAR regions of sex chromosomes, all female genotypes and male heterozygous genotypes were set to missing, and male homozygous variants were converted to haploid. The multi-allelic variants in the datasets were then decomposed using hl.split_multi_hts.

**Supplementary Table 1 | Sample counts before and after hard and release filters**

| **Description** | **Genomes** (% remaining from previous stage) | **Exomes** (% remaining from previous stage) |
|---|---|---|
| Before filters | 20,314 | 164,332 |
| After hard filters | 20,120 (99.04%) | 160,064 (97.40%) |
| After hard + release permission filters | 17,016 (84.57%) | 141,748 (88.56%) |

**Supplementary Table 2 | Counts by data type and hard filter**

| **Data type** | **Hard filter** | **Number of samples** | **% of call set** |
|---|---|---|---|
| Exomes | High contamination | 3,300 | 2.01% |
| | Ambiguous sex | 896 | 0.55% |
| | Excess chimeric reads | 649 | 0.39% |
| | Sex chromosome aneuploidy | 94 | 0.06% |
| | Low call rate | 61 | 0.04% |
| | Low coverage | 19 | 0.01% |
| Genomes | Low median insert size | 70 | 0.34% |
| | High contamination | 53 | 0.26% |
| | Excess chimeric reads | 43 | 0.21% |
| | Ambiguous sex | 34 | 0.17% |
| | Low coverage | 16 | 0.08% |
| | Low call rate | 4 | 0.02% |

*Platform imputation for exomes*

Capture and sequencing platform metadata were only available for only a fraction of the samples in the exome call set. For the remaining exome samples, we performed platform imputation by compiling a list of known exome capture regions across multiple exome capture products and considering the set of biallelic variants that fell within these regions. For each sample passing hard filters, we computed the biallelic variant call rate for each exome capture interval on the list. We then discretized the per-sample, per-interval call rates into two categories, called and not-called (based on a per-interval call rate threshold of 0.25) and performed principal components analysis (PCA) on these thresholded values. The first two principal components are plotted in Extended Data Fig. 1c. Samples were then clustered using the top 9 principal components using HDBSCAN with a 'min_cluster_size' parameter setting of 100. Generic platform labels were assigned to each cluster discovered by the PCA-HDBSCAN analysis (Supplementary Table 3). For the samples with known platforms, we find that these imputed labels generally correspond with a single known platform, suggesting the utility of this approach (Supplementary Table 4). Aside from the subdivision of Illumina Nextera samples into two distinct sub-platforms of unknown validity, the rate of inconsistent classification using our platform imputation approach was 0.89% (87+89+106 samples).

**Supplementary Table 3 | Exome platform assignments**

| Imputed platform label | Count | % overall |
|---|---:|---:|
| Unassigned | 4,244 | 2.65% |
| 0 | 1,170 | 0.73% |
| 1 | 7,241 | 4.52% |
| 2 | 3,195 | 2.00% |
| 3 | 152 | 0.09% |
| 4 | 1,279 | 0.80% |
| 5 | 1,906 | 1.19% |
| 6 | 527 | 0.33% |
| 7 | 5,928 | 3.70% |
| 8 | 158 | 0.10% |
| 9 | 28,655 | 17.90% |
| 10 | 47,251 | 29.52% |
| 11 | 152 | 0.09% |
| 12 | 415 | 0.26% |
| 13 | 447 | 0.28% |
| 14 | 574 | 0.36% |
| 15 | 56,772 | 35.47% |

**Supplementary Table 4 | Confusion matrix for exome samples with known platform labels**

| Imputed Platform Label | Agilent SureSelect v1 | Agilent SureSelect v2 | Agilent SureSelect v3 | Illumina Nextera | NimbleGen SeqCap v2 |
|---|---|---|---|---|---|
| Unassigned | 0 | 87 | 0 | 106 | 0 |
| 0 | 0 | 0 | 0 | 0 | 0 |
| 1 | 0 | 0 | 0 | 0 | 0 |
| 2 | 0 | 0 | 0 | 0 | 0 |
| 3 | 114 | 0 | 0 | 0 | 0 |
| 4 | 0 | 0 | 0 | 0 | 0 |
| 5 | 0 | 0 | 585 | 0 | 0 |
| 6 | 0 | 0 | 0 | 0 | 0 |
| 7 | 0 | 0 | 0 | 0 | 509 |
| 8 | 0 | 0 | 0 | 0 | 0 |
| 9 | 0 | 0 | 0 | 8,796 | 0 |
| 10 | 0 | 89 | 0 | 6,544 | 0 |
| 11 | 0 | 0 | 0 | 0 | 0 |
| 12 | 0 | 0 | 0 | 0 | 0 |
| 13 | 0 | 0 | 0 | 0 | 0 |
| 14 | 0 | 0 | 0 | 0 | 0 |
| 15 | 0 | 14,937 | 0 | 0 | 0 |

*Relatedness filters*

We used the PC-Relate[55] method as implemented in Hail (https://hail.is/docs/0.2/methods/genetics.html#hail.methods.pc_relate) to compute pairwise relatedness between all samples passing hard filters in the joint exome and genome call set. Variant allele frequencies were recomputed over the joint exome and genome call set, and only bi-allelic variants with joint allele frequencies > 0.001 and high joint call rates (> 0.99) were considered (n=94,177). Variants were LD-pruned ($r^2 = 0.1$) and alternate allele counts from the pruned variant set were used for an initial PCA. The top 10 components from the PCA were then used in the PC-Relate computation, along with a 'min_individual_maf' parameter setting of 0.05.

Pairwise kinship coefficients from PC-Relate were then used to group samples into clusters of related individuals. Sample pairs with kinship coefficients greater than 0.0883 (corresponding to second-degree or closer) were considered related (Supplementary Table 5). We used the Hail maximal_independent_set() function to prune clusters of related individuals in such a way as to maximize the number of unrelated samples retained for each familial cluster. To resolve cases for which there were multiple possible pruning options, we created an ordered list that assigned a rank to each sample in the call set, with more desired samples receiving a lower rank than less desirable samples, and based on the ranking, we flagged less desirable samples for removal before more desirable samples.

**Supplementary Table 5 | Pair counts by degree of relatedness**

| Degree of relatedness | Number of pairs in joint callset |
|---|---:|
| Duplicates or twins | 5,012 |
| 1st degree | 23,939 |
| 2nd degree | 6,611 |
| 3rd degree or unrelated | 10,457 |

Samples with release permissions were preferred over samples without release permissions. Genome samples were preferred over exome samples. Among the genome samples, PCR-free samples were preferred over PCR-plus samples, parent samples in several trio studies were chosen over child samples, and samples with a higher average depth of coverage were preferred over those with lower coverage. For exomes, samples were ranked first according to preference for internally- over externally-sequenced samples; then in preference for more recently sequenced samples over less recently sequenced samples; subsequently in preference for parents over children in several designated trios; and lastly in preference for greater fraction of bases covered above 20x depth over lower fraction of bases covered above 20x depth.

We defined the maximal set of unrelated individuals after pruning familial clusters as the set of unrelated individuals in gnomAD, retaining the number of samples as indicated in Supplementary Table 6.

**Supplementary Table 6 | Sample counts by relatedness status**

| | **Genomes** (% total) | **Exomes** (% total) |
|---|---|---|
| Related samples | 1,806 (8.98%) | 17,308 (10.81%) |
| Unrelated samples | 18,314 (91.02%) | 142,758 (89.19%) |

*Population and subpopulation inference*

We then inferred continental ancestry on the set of unrelated samples using a PCA and random forest approach. Restricting our analysis to unrelated samples, we computed the top 20 principal components on the alternate allele counts for the same set of variants used in the PC-Relate PCA and projected the remaining related samples onto these principal components (Supplementary Fig. 1). Next, we trained a random forest model on a set of samples with known continental ancestry and used this model to assign continental ancestry labels to samples for which the random forest probability > 0.9.

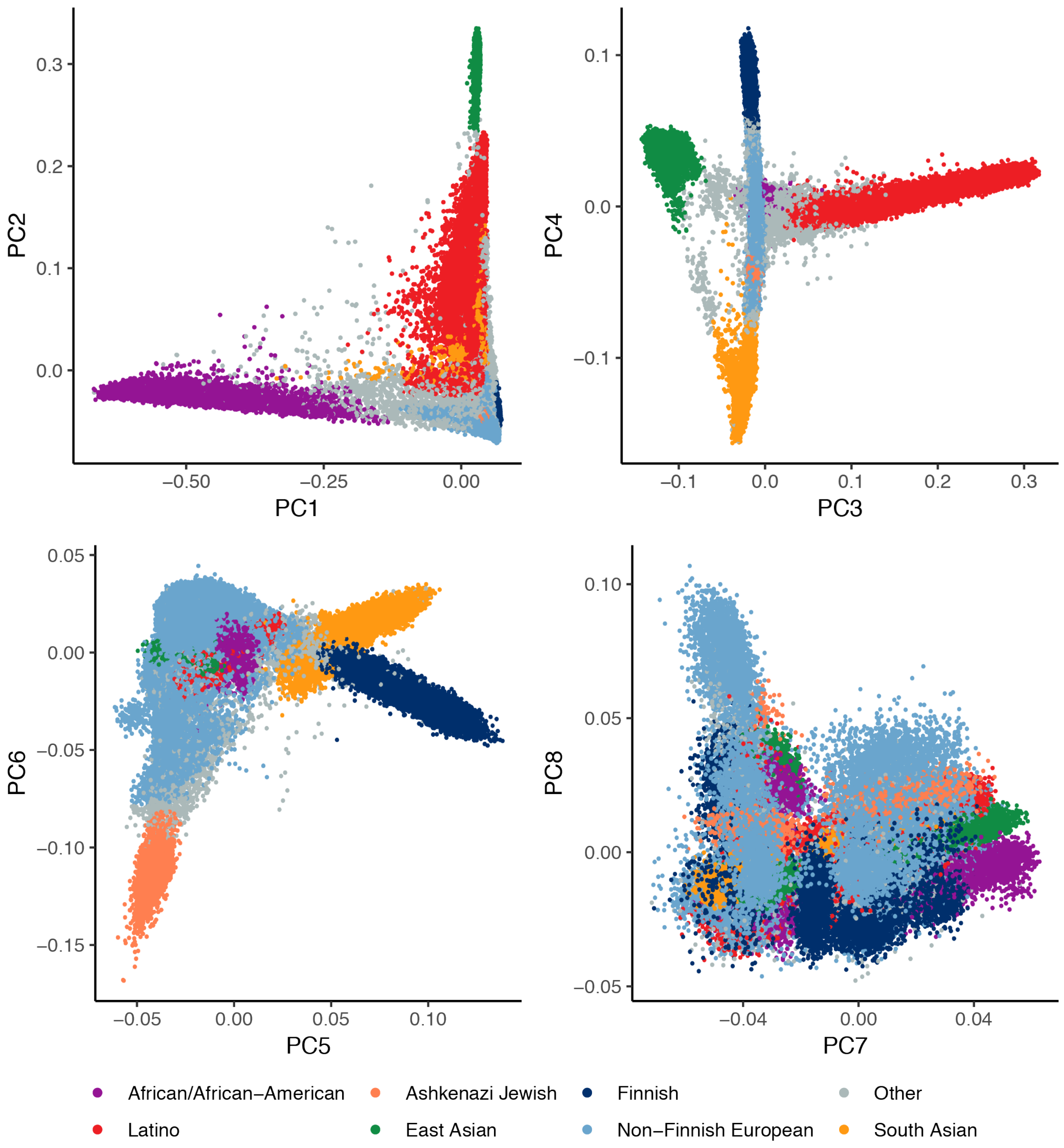


**Supplementary Figure 1 | Continental ancestry principal components.** The top ancestry principal components are shown, such that a dot represents a single sample colored by its inferred continental ancestry. The top 6 principal components were used to infer continental ancestry for 141,456 samples.

There were sufficient sample sizes of European and East Asian continental ancestry to perform more fine-grained subpopulation inference, and for these populations we performed another round of PCA on unrelated individuals, considering only variants with a continental-specific allele frequency > 0.001 and a call rate >= 0.999 across all platforms (or no more than 1 missing sample per platform if the number of samples sequenced on a platform was less than 1000). We LD-pruned these variant sets ($r^2 = 0.1$; 11,842 variants for East Asians and 84,792 variants for Europeans) and ran PCA using the alternate allele counts for these variants, obtaining sub-continental ancestry principal components (Supplementary Fig. 2). For both European and East Asian cohorts, we then projected related samples onto the sub-continental PCs. Random forest models were then trained on samples with known sub-population labels (e.g., Bulgarian and Estonian for Europeans; Korean for East Asian) and applied to samples without sub-population labels. Samples that were assigned sub-population labels by the random forest models with probabilities < 0.9 were collected into a generic “other” sub-population category.

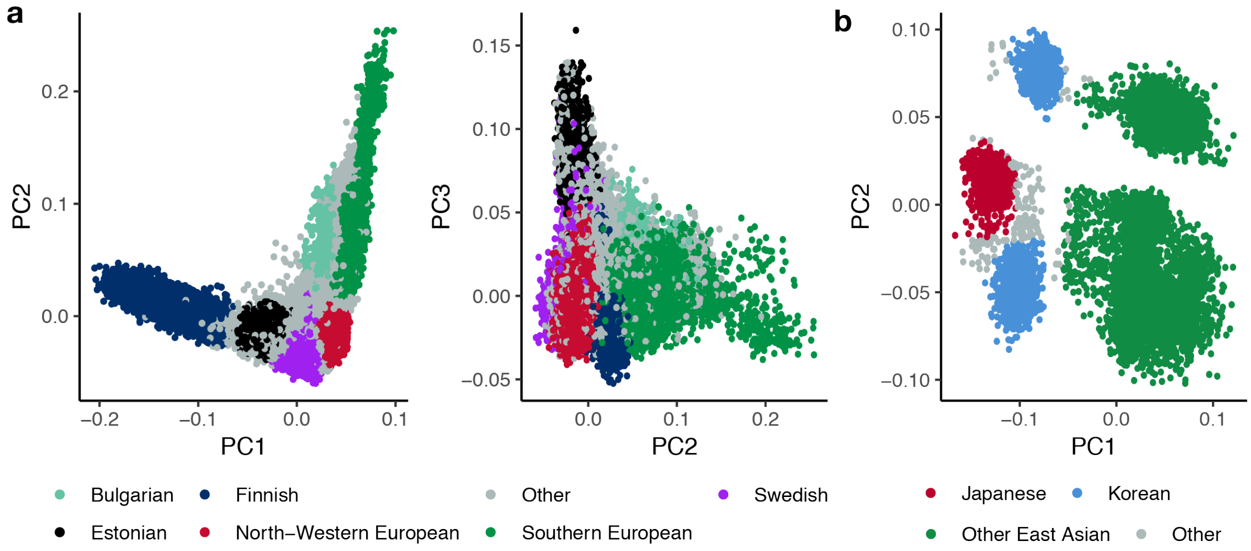


**Supplementary Figure 2 | Sub-continental ancestry principal components.** The top sub-continental ancestry principal components are shown, such that a dot represents a single sample colored by its inferred sub-continental ancestry. For 64,603 European samples (a), three principal components were used for sub-population inference; for 9,977 East Asian samples (b), two principal components were used for sub-population inference.

The final count of samples per-population and sub-population (after final population- and platform-specific filtering, see below) is shown in Supplementary Table 7. A two-dimensional visualization of the sub-populations present in gnomAD (Fig. 1a) was created by applying the

UMAP algorithm[20] to seven ancestry PCs with a min_dist parameter of 0.5 and an n_neighbors parameter of 30.

**Supplementary Table 7 | Population and subpopulation counts**

| Population | Code | Genomes | Exomes | Total |
|---|---|---|---|---|
| African/African-American | afr | 4,359 | 8,128 | 12,487 |
| Latino/Admixed American | amr | 424 | 17,296 | 17,720 |
| Ashkenazi Jewish | asj | 145 | 5,040 | 5,185 |
| East Asian | eas | (780) | (9,197) | (9,977) |
| Koreans | kor | - | 1,909 | 1,909 |
| Japanese | jpn | - | 76 | 76 |
| Other East Asian | oea | 780 | 7,212 | 7,992 |
| Finnish | fin | 1,738 | 10,874 | 12,526 |
| Non-Finnish European | nfe | (7,718) | (56,885) | (64,603) |
| Bulgarian | bgr | 0 | 1,335 | 1,335 |
| Estonian | est | 2,297 | 121 | 2,418 |
| North-Western European | nwe | 4,299 | 21,111 | 25,410 |
| Southern European | seu | 53 | 5,752 | 5,805 |
| Swedish | swe | 0 | 13,067 | 13,067 |
| Other non-Finnish European | onf | 1,069 | 15,499 | 16,568 |
| South Asian | sas | - | 15,308 | 15,308 |
| Other (population not assigned) | oth | 544 | 3,070 | 3,614 |
| *Total* | | *15,708* | *125,748* | *141,456* |

##### *Population- and platform-specific filters*

After inferring population and platform labels, we computed a battery of sample quality control metrics on all samples in the call set passing hard filters: number of deletions, number of insertions, number of SNVs, ratio of deletions to insertions, ratio of transitions to transversions, and ratio of heterozygous to homozygous variants. These metrics were analyzed separately per continental population group and per inferred sequencing platform, as underlying distributions for these metrics varied widely both by population and by sequencing platform. Outliers were defined as samples with values outside four median absolute deviations (MAD) from the median of a given metric, with median and MAD values computed over all samples sharing the same ancestry and platform labels. A sample of the process is plotted in Extended Data Fig. 1d, and the number of samples failing per platform is summarized in Supplementary Table 8.

**Supplementary Table 8 | Summary of outliers per population and platform grouping**

| Data type | Platform (inferred for exomes, known for genomes) | Samples with at least one outlier metric | Total Samples | % of Platform |
|---|---|---|---|---|
| Exomes | Unassigned | 129 | 4,243 | 3.04% |
| | 0 | 67 | 1,170 | 5.73% |
| | 1 | 92 | 7,241 | 1.27% |
| | 2 | 59 | 3,195 | 1.85% |
| | 3 | 10 | 152 | 6.58% |
| | 4 | 39 | 1,279 | 3.05% |
| | 5 | 153 | 1,906 | 8.03% |
| | 6 | 64 | 527 | 12.14% |
| | 7 | 503 | 5,928 | 8.49% |
| | 8 | 3 | 158 | 1.90% |
| | 9 | 666 | 28,655 | 2.32% |
| | 10 | 927 | 47,251 | 1.96% |
| | 11 | 6 | 152 | 3.95% |
| | 12 | 7 | 415 | 1.69% |
| | 13 | 22 | 447 | 4.92% |
| | 14 | 8 | 574 | 1.39% |
| | 15 | 3,382 | 56,771 | 5.96% |
| Genomes | Legacy Standard High Coverage Whole Genome Sequencing (30x) | 8 | 131 | 6.11% |
| | Low Input Human WGS (Standard Coverage) | 22 | 56 | 39.29% |
| | PCR-Free Human Genome 30x | 32 | 832 | 3.85% |
| | PCR-Free Human WGS (Lite Coverage) | 192 | 9384 | 2.05% |
| | PCR-Free Human WGS (Standard Coverage) | 52 | 712 | 7.3% |
| | PCR-Free Human WGS (Standard Coverage) + Decoy | 196 | 2913 | 6.73% |
| | Standard High Coverage Whole Genome Sequencing (30x) | 3 | 10 | 30.0% |
| | Standard High Coverage Whole Genome Sequencing (30x) \| High Read Length | 306 | 5124 | 5.97% |
| | Standard High Coverage Whole Genome Sequencing (30x) \| Low Read Length | 102 | 958 | 10.65% |

*Finalizing samples in the gnomAD v2.1 release*

The final set of samples included in the gnomAD v2.1 release (125,748 exomes and 15,708 genomes, for a total of 141,456) was defined to be the set of unrelated samples with release permissions, no hard filter flags, and no population- and platform-specific outlier metrics (Supplementary Table 9), including 64,754 females and 76,702 males.

Finally, we created several subsets of samples from the release dataset requested by popular demand: controls-only, comprised of samples not designated as cases in the common disease studies from which they originated; non-topmed, comprised of samples not included in the Trans-Omics for Precision Medicine (TOPMED)-BRAVO cohort (i.e., unique to gnomAD relative to TOPMED); non-neuro, comprised of samples not ascertained for a neurological phenotype; and non-cancer, comprised of samples not included in cancer cohort studies. Global and population-level allele frequencies were recomputed for each of these cohorts and included in the release datasets.

**Supplementary Table 9 | Sample counts by filtering stage**

| **Description** | **Genomes** (% remaining from previous stage) | **Exomes** (% remaining from previous stage) |
|---|---|---|
| Before filters | 20,314 | 164,332 |
| After hard filters | 20,120 (99.04%) | 160,064 (97.40%) |
| After hard + release permission filters | 17,016 (84.57%) | 141,748 (88.56%) |
| After hard, release permission, and relatedness filters | 16,288 (95.72%) | 130,645 (92.17%) |
| Final release: After hard, release permission, relatedness, and outlier metric filters | 15,708 (96.44%) | 125,748 (96.25%) |

**Supplementary Table 10 | Sample counts for genomes and exomes in gnomAD subsets**

| | Exomes | Genomes | Total |
|---|---|---|---|
| Full release dataset | 125,748 | 15,708 | 141,456 |
| Controls only | 54,704 | 5,442 | 60,146 |
| Non-TOPMED | 122,439 | 13,304 | 135,743 |
| Non-neuro | 104,068 | 10,636 | 114,704 |
| Non-cancer | 118,479 | 15,708 | 134,187 |

### Variant QC

Laurent C. Francioli, Konrad J. Karczewski, Grace Tiao, Daniel G. MacArthur

We next sought to define a high-quality set of variation for release and downstream analysis. For variant QC, we considered the variants present in the 141,456 release samples (see above), as well as sites present in family members forming trios that passed all of the sample QC filters (212 trios in genomes, 4,568 trios in exomes), which allowed us to look at transmission and Mendelian violations for evaluation purposes. Variant QC was performed on the exomes and genomes separately but using the same pipeline (although different thresholds were used), which is available at https://github.com/macarthur-lab/gnomad_qc/tree/master/variant_qc. We excluded variants on the basis of both hard filters and a random forest (RF) model that we developed (details below).

##### *Hard filters*

We excluded (1) all variants that showed an excess of heterozygotes, defined by an inbreeding coefficient < -0.3, and (2) all variants for which no sample had a high quality genotype (as described above: depth >= 10, genotype quality >= 20 and minor allele fraction >= 0.2 for all non-reference alleles of heterozygous genotypes).

##### *Random Forest model*

We implemented a random forest (RF) model using Hail / PySpark to distinguish true genetic variants from artifacts. Our model considers SNVs and indels together and operates on each variant allele separately (as opposed to each variant site). This model outperformed the state-of-the-art Genome Analysis Toolkit (GATK) Variant Quality Score Recalibration (VQSR) based on the quality metrics we looked at (details below). All code is open-source and available at https://github.com/macarthur-lab/gnomad_hail/blob/master/utils/rf.py.

Features

The variant annotations (features) we used to train the model came from two sources: (1) site-level annotations from the GATK HaplotypeCaller (which at present, does not output any allele-level quality annotations) and (2) allele-level annotations that we computed using Hail. In addition to these variant quality annotations, we also included categorical annotations that describe the type of and context of the variant allele. Supplementary Table 11 summarizes the features we used and shows their relative importance in the final random forest model we trained. Note that since random forests do not tolerate missing data, we imputed all missing values using the median value for that feature.

**Supplementary Table 11 | Features used in final random forest model**

| | | | **Importance** | |
|---|---|---|---|---|
| **Feature** | **Description** | **Type** | **Exomes** | **Genomes** |
| InbreedingCoeff | Deviation from Hardy-Weinberg expectation of observed heterozygotes | Site | 0.123 | 0.042 |
| StrandOddsRatio | Strand bias estimated by the Symmetric Odds Ratio test | Site | 0.100 | 0.266 |
| ReadPosRankSum | Rank Sum Test for relative positioning of REF versus ALT alleles within reads | Site | 0.061 | 0.060 |
| MappingQualityRankSum | Rank Sum Test for mapping qualities of REF versus ALT reads | Site | 0.019 | 0.031 |
| qd (quality by depth) | Sum of the non-reference genotype likelihoods divided by the total depth in all carriers of the allele | Allele | 0.618 | 0.470 |
| pab_max | Highest p-value for sampling the observed allele balance under a binomial model in any heterozygote | Allele | 0.065 | 0.106 |
| allele_type | SNV or indel | Allele | 0.0003 | 0.001 |
| n_alleles | Number of alleles at the site | Site | 0.006 | 0.006 |
| mixed_site | True if both SNVs and indels are present at the site | Site | 0.0005 | 0.0005 |
| spanning_deletion | True if one or more deletions at other sites overlap the site | Site | 0.002 | 0.012 |

Training

We used alleles that were previously genotyped or discovered with high confidence as positive training sites (Supplementary Table 12). Because these variants are mostly common, we also included transmitted singletons, which are alleles that were found in only a single individual in the unrelated gnomAD samples and for which we observed Mendelian transmission to an offspring in one of the gnomAD trios. For negative training examples, we used alleles that fail the traditional GATK hard filters.

**Supplementary Table 12 | Random forest training examples**

| Name | Description | Class | Number of alleles | |
|---|---|---|---|---|
| | | | Exomes | Genomes |
| Omni 2.5 | SNVs present on the Omni 2.5 genotyping array and found in 1000 genomes (from the GATK bundle) | TP | 95K | 2.3M |
| Mills/Devine | Indels present in the Mills and Devine data (from the GATK bundle) | TP | 12K | 1.3M |
| 1000 Genomes high-quality sites | Sites discovered in 1000 Genomes with high confidence (from the GATK bundle) | TP | 560K | 29M |
| Transmitted singletons | Singletons in gnomAD unrelated samples that are transmitted to an offspring excluded from the gnomAD release | TP | 106K | 116K |
| Failing hard filers | Variants failing traditional GATK hard filters: QD < 2 \|\| FS > 60 \|\| MQ < 30 | FP | 789K | 31M |

To train our model, we randomly subsetted the training examples to get a balanced set of positive and negative training examples. Chromosome 20 was entirely left out of the training for evaluation purposes. We trained models using 500 trees with a maximum depth of 5, for exomes and genomes separately.

Evaluation and threshold selection

While using a balanced set of training data allowed us to train our model without a prior on either positive or negative class, it is also not representative of the true positive rate in our data. For this reason, we evaluated our model using metrics that consider both common and rare variants and adapted our filtering threshold for SNVs and indels and for exomes and genomes. In addition, we also compared the performance of our model against the state-of-the-art GATK Variant Quality Score Recalibration (VQSR).

The threshold cutoffs were chosen manually based on the evaluation metrics presented below and they were set independently for SNVs (90% SNVs retained) and indels (80% of indels retained). These cutoffs broadly maximized our sensitivity and specificity based on the collective evaluation of the metrics below.

To evaluate the performance of our model for filtering common variants, we investigated the precision and recall of our model on two samples for which gold standard variation is available: NA12878[46] (Extended Data Fig. 2a-d) and a synthetic diploid mixture[47] (Extended Data Fig. 2e-h). For both samples, our model was either superior or similar to GATK VQSR for both SNVs and indel at the chosen threshold (intersection of the dashed lines in the figures representing 10% and 20% of SNVs and indels filtered, respectively).

To evaluate the performance of our model for filtering rare variants, we ranked all variants by the score output by our model and then binned them in percentiles, so that every point on the plot represents the same number of alleles. We also ranked and binned our variants based on the GATK VQSR score for comparison purposes. We then evaluated the following three metrics:

1. The number of de novo calls per child (Extended Data Fig. 3a-d): For each of our evaluation trios, we counted the number of alleles that were found in a single child and no other sample in the callset (i.e. a *de novo* mutation call private to the trio). We only expect to observe ~1.6 *de novo* SNVs and ~0.1 de novo indels per exome, and ~65 *de novo* SNVs and ~5 *de novo* indels per genome[21]. The shape of the curves show that we find a relatively low number of *de novo* calls at higher confidence (presumably mostly true *de novo* variation) and then a relatively large number of *de novo* calls with low confidence (presumably errors). The sharp increase in the number of *de novo* calls begins around our chosen cutoffs (dashed lines in the figures representing 10% and

20% of SNVs and indels filtered, respectively). The number of *de novo* calls per child at our cutoff is very close to expectations for SNVs, suggesting very good specificity, and ~2x the number of expected indels, suggesting that for this class of especially challenging variants we may still have a relatively high error rate (or current rate estimates maybe too low). Note that in all cases our model outperforms GATK VQSR.

2. The number of transmitted singletons (Extended Data Fig. 3e-h): Here, we used transmitted singletons on chromosome 20, which were held out for evaluation purposes. Because these very rare variants were observed independently in a parent and a child (consistent with Mendelian transmission) but not in any other samples, we expect them to represent true rare variation. In all cases (exomes and genomes, SNVs and indels), our model outperforms GATK VQSR by classifying more of these variants as true positives. At our chosen cutoff threshold, we keep 99.5% and 97.8% of the transmitted singleton SNVs in exomes and genomes, respectively (vs 90% of all SNVs), and 95.2% and 96.9% of the transmitted singleton indels in exomes and genomes respectively (vs 80% of all indels).
3. The number of validated *de novo* mutations (exomes only; Extended Data Fig. 3i-j): We had access to 295 SNVs and 80 indels from 331 exome samples that were validated in previous studies[48]. For both SNVs and indels, we outperform GATK VQSR and at our chosen cutoff threshold, we keep 96.2% of the validated *de novo* SNVs (vs 90% of all SNVs) and 97.5% of the validated *de novo* indels (vs 80% of all indels).

##### *Final variant counts*

For exomes, our filtration process removes 12.2% of SNVs (RF probability >= 0.1) and 24.7% of indels (RF probability >= 0.2). For genomes, we filtered 10.7% of SNVs (RF probability >= 0.4) and 22.3% of indels (RF probability >= 0.4). Supplementary Table 13 shows the number of variants filtered and retained by our filtering strategy.

The 14,078,157 SNVs in the exomes span 11,999,542 genomic positions, representing

20.1% of the 59,837,395 bases where calling was performed. When filtering observed and possible sites to a median of 30X coverage, we observe 21.9% of sites with at least one SNV. The 204,063,503 SNVs in the genomes span 192,608,400 genomic positions, representing 6.8% of the 2,831,728,308 bases where calling was performed.

**Supplementary Table 13 | Variant counts by filtering status**

| | | SNVs | | Indels | |
|---|---|---|---|---|---|
| | | Passing hard filters | Failing hard filters | Passing hard filters | Failing hard filters |
| Exomes | Passing RF filter | **14,078,157** | 797,205 | **889,254** | 56,136 |
| | Failing RF filter | 525,783 | 626,685 | 148,750 | 88,002 |
| Genomes | Passing RF filter | **204,063,503** | 1,326,773 | **25,925,202** | 727,296 |
| | Failing RF filter | 18,713,041 | 4,466,895 | 5,312,957 | 1,406,669 |

*Comparison of whole-exome and whole-genome coverage in coding regions*

gnomAD 2.1 is a unique collection of sequenced samples, mixing whole-exome sequencing using many different whole-exome capture kits with whole-genome sequencing using with and without PCR amplification steps in sample preparation (PCR+ and PCR-free). To evaluate how well we captured the coding part of the genome overall and the potential for whole-genome data to complement whole-exome data in coding regions, we compared the coverage between samples sequenced on different technologies in the protein-coding part of the genome. We used 19,636 genes on the basis of all the coding regions (CDS) from the gencode v.19 GTF file. We then annotated each coding base with the coverage in a random subset of 10% of our samples (as described above) and analyzed how well each gene was covered by sequencing platform. The whole-exome sequencing platforms were inferred based on PCA (see Platform imputation for exomes) and we only show here platforms for which we computed coverage for at least 100 samples. Clusters correspond to platforms from specific vendors, but should not be regarded as representative of the overall quality of any provider due to substantial ascertainment biases and confounding by project and sequencing provider. In this analysis, Cluster 9 refers to Illumina capture platforms with 151 bp reads, Cluster 10 to Illumina capture platforms with 76 bp reads, and Cluster 15 to Agilent products. We do not have information regarding Cluster 5 and 7 products. The whole-genomes were split into PCR+ and PCR-free. We defined a base as well-covered if at least 80% of the samples sequenced were covered at a depth of at least 20x (10x for males on non-pseudoautosomal regions of the X chromosome). As can be seen in Supplementary Fig. 3, ~80% of protein-coding genes are well-captured by all technologies, whole-genome sequencing captures ~8% additional genes well and about 2.5% of the genes are not captured by either.

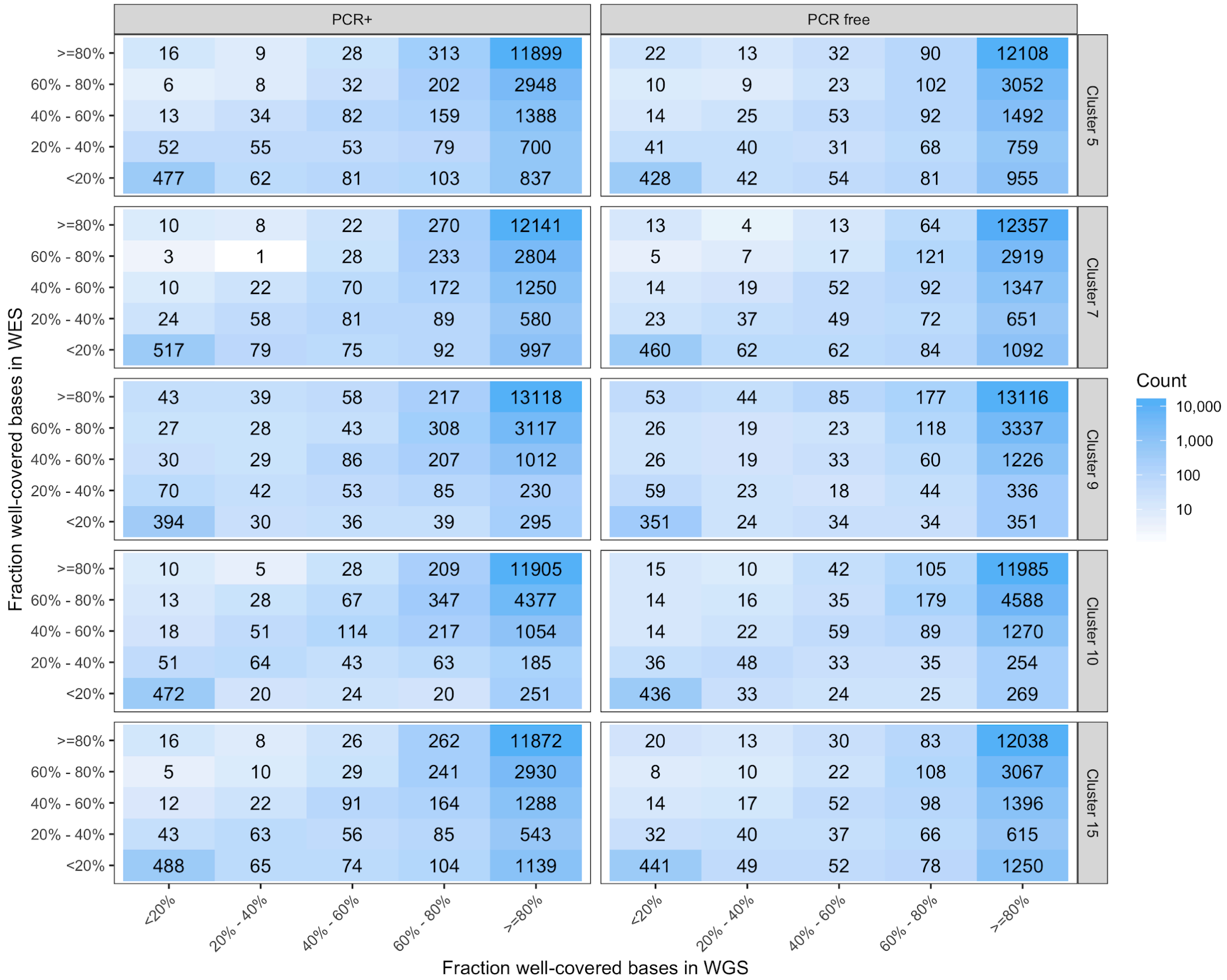


**Supplementary Figure 3 | Comparison of gene coverage by sequencing technology.** Each cell shows the number of genes that correspond to a given bin of bases well-covered by a given whole-exome sequencing platform (rows) and a whole-genome sequencing platform (columns).

In addition, ~1% of the genes in Supplementary Fig. 3 are better captured by whole-exome sequencing. We hypothesized that this may be due to differences in our alignment pipeline which uses bwa-mem for genomes and bwa-aln for exomes, leading to differences in homologous regions. To investigate this further, we computed the mean mappability for each gene using the UCSC duke35bp mappability track and compared the mappability of genes where >50% of bases are well-captured against that of genes where <50% of the bases are well-captured (Supplementary Fig. 4). We found that indeed the vast majority of the genes not well-captured by whole-genome sequencing have low mappability. We further observe that

PCR-free whole-genome sequencing well captures another 121 genes that could not be captured with PCR+ whole-genome sequencing. These genes have a high mappability score, so it is likely that they have abnormal GC contents and thus are difficult to capture using PCR+ whole-genome sequencing. Finally, while the low-mappability genes are also poorly-covered in exomes, this gene set is dominated by genes with high mappability, and are simply not captured by the exome kit.

The entire table of coverage summary for each gene and each platform can be found in Supplementary Dataset 1 and at: https://storage.googleapis.com/gnomad-public/papers/2019-flagship-lof/v1.1/summary_gene_coverage/gencode_grch37_gene_by_platform_coverage_summary.tsv.gz

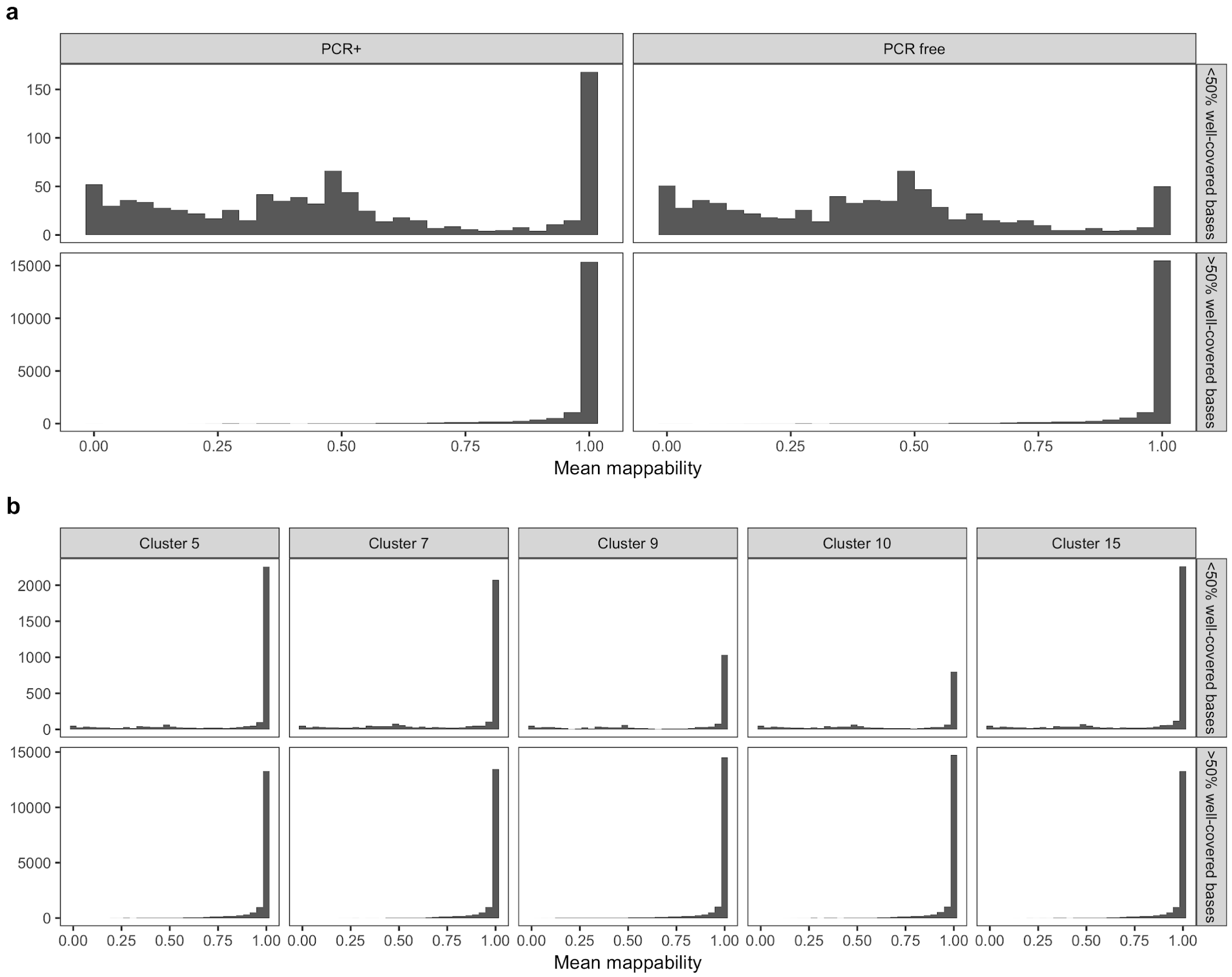


**Supplementary Figure 4 | Mappability and gene coverage.** Histogram showing the number of genes that are poorly or well-covered by mean gene mappability, broken down by platform for whole-genome sequencing (a) and whole-exome sequencing (b).

### Variant annotation

Konrad J. Karczewski, Daniel P. Birnbaum, Moriel Singer-Berk, Daniel Rhodes, Eleanor G. Seaby, Kristen M. Laricchia, Beryl B. Cummings, Laurent C. Francioli, Grace Tiao, Cotton Seed, Monkol Lek, Daniel G. MacArthur

#### *Frequency and context annotation*

All frequency annotations were calculated using a custom script written in Hail (https://github.com/macarthur-lab/gnomad_qc/blob/master/annotations/generate_frequency_data.py). Briefly, after filtering to the high quality samples with permissions for data release as described above, we applied the call_stats aggregator to compute the allele count, allele number, and number of homozygotes at every site. After filtering to high-quality genotypes (GQ >= 20, DP >= 10, and for heterozygotes, that each alternate allele has at least 20% of reads supporting the allele), these calculations are repeated separately for the full dataset, males and females, for each major population (also split by males and females), each subpopulation, and each computationally predicted capture platform (see Sample QC). Additionally, the full dataset and each population are downsampled to various sample numbers (where sufficient samples exist, 10, 20, 50, 100, 200, 500, 1000, 2000, 5000, 10000, 15000, 20000, 25000, 30000, 35000, 40000, 45000, 50000, 55000, 60000, 65000, 70000, 75000, 80000, 85000, 90000, 95000, 100000, 110000, and 120000, as well as additional downsamplings of the total numbers of samples of each population). The maximum frequency across continental populations (aside from Ashkenazi Jewish, Finnish, or Other) is stored as the “popmax” frequency. The filtering frequency described previously[56] is implemented in Hail (https://hail.is/docs/0.2/experimental/index.html#hail.experimental.filtering_allele_frequency) and computed for the full dataset and each population (separately for males and females on the sex chromosomes) at 95% and 99% confidence interval levels. Finally, we compute the histogram of ages of heterozygous and homozygous variant carriers using Hail's hist aggregator. This process is repeated for the subsets of the data as described in Supplementary Table 10.

A dataset of every possible SNV in the human genome (2,858,658,098 sites x 3 substitutions at each site = 8,575,974,294 variants) along with 3 bases of genomic context was created using the GRCh37 reference. This dataset was annotated with methylation data for all CpG variants and coverage summaries as described above, and was subsequently used to annotate the exome and genome datasets where required downstream. The number of variants observed at each downsampling, broken down by variant class, is shown in Extended Data Fig. 4a. As previously shown[4], the CpG sites begin to saturate at a sample size of about 10,000 individuals, which affects the callset-wide transition/transversion (TiTv) ratio (Extended Data Fig. 4b). In order to compute the proportion of possible variants observed, we filtered the dataset of all possible SNVs to the exome calling intervals described previously[4] and considered only bases where exome coverage was >= 30X (Extended Data Fig. 4c).

*Functional annotation*

Variants were annotated using the Variant Effect Predictor (VEP) version 85[57] against Gencode v19[58], implemented in Hail with the LOFTEE plugin, described below. The 17,209,972 and 261,942,336 variants in the gnomAD exomes and genomes, respectively, were annotated, as well the 8,575,974,294 possible SNVs in the human genome and 429,237 variants in the ClinVar dataset described above. VEP performs annotation against individual transcripts: in most downstream analyses, unless otherwise specified, variants were filtered to canonical transcripts as defined by Gencode/Ensembl. To assess these annotations, we use the mutability-adjusted proportions of singletons (MAPS) score previously described[4], implemented as a Hail module. Notably, in this manuscript, we calibrate MAPS using the mutation rates computed below (see Mutational Model), which includes methylation level as a feature. MAPS is a relative metric, and so cannot be compared across datasets, but is a useful summary metric for the frequency spectrum, indicating deleteriousness as inferred from rarity of variation (high values of MAPS correspond to lower frequency, suggesting the action of negative selection at more deleterious sites). Using this metric, we find nonsense, essential splice, and frameshift variants as the classes of variation which undergo the greatest degree of negative selection (Fig. 1c,d), as previously observed, which is also observed by lower proportion of possible variants observed (Extended Data Fig. 4d; Supplementary Table 14; Supplementary Fig. 5). We provide the number of variants observed and possible number of variants by functional class, variant type (context, reference, and alternative allele for SNVs; length for indels), and median coverage, for each of the downsamplings computed in gnomAD as Supplementary Datasets 2-5.

Further, we compute the number of pLoF variants discovered in each population for each downsampling described above, and show the number of new pLoFs added as a function of each new individual sequenced (Extended Data Fig. 4e,f): at current sample sizes, we observe ~1.87 new pLoF variants for each additional individual exome sequenced.

**Supplementary Table 14 | Variants observed by category in 125,748 exomes**

| Annotation | Variant Type | Methyl. Level | Number Observed | Number Possible | Percent Observed | Number Singletons | Percent Singleton |
|---|---|---|---|---|---|---|---|
| missense | CpG | 0 | 26,496 | 112,509 | 23.55 | 12,900 | 48.69 |
| missense | CpG | 1 | 24,966 | 53,228 | 46.90 | 7,242 | 29.01 |
| missense | CpG | 2 | 667,832 | 884,142 | 75.53 | 118,669 | 17.77 |
| missense | non-CpG transition | - | 2,041,008 | 16,912,929 | 12.07 | 1,177,541 | 57.69 |
| missense | transversion | - | 1,788,005 | 44,340,017 | 4.03 | 1,120,233 | 62.65 |
| nonsense | CpG | 0 | 375 | 1,997 | 18.78 | 204 | 54.40 |
| nonsense | CpG | 1 | 780 | 1,829 | 42.65 | 275 | 35.26 |
| nonsense | CpG | 2 | 28,594 | 54,975 | 52.01 | 9,785 | 34.22 |
| nonsense | non-CpG transition | - | 50,078 | 684,665 | 7.31 | 33,851 | 67.60 |
| nonsense | transversion | - | 53,192 | 2,802,136 | 1.90 | 37,165 | 69.87 |
| synonymous | CpG | 0 | 20,272 | 68,718 | 29.50 | 8,914 | 43.97 |
| synonymous | CpG | 1 | 16,563 | 31,021 | 53.39 | 4,116 | 24.85 |
| synonymous | CpG | 2 | 353,931 | 416,729 | 84.93 | 39,270 | 11.10 |
| synonymous | non-CpG transition | - | 1,314,239 | 8,937,863 | 14.70 | 718,666 | 54.68 |
| synonymous | transversion | - | 468,105 | 9,458,221 | 4.95 | 274,277 | 58.59 |
| frameshift | indel | - | 184,911 | - | - | 129,015 | 69.77 |
| inframe deletion | indel | - | 61,110 | - | - | 33,724 | 55.19 |
| inframe insertion | indel | - | 20,076 | - | - | 12,027 | 59.91 |

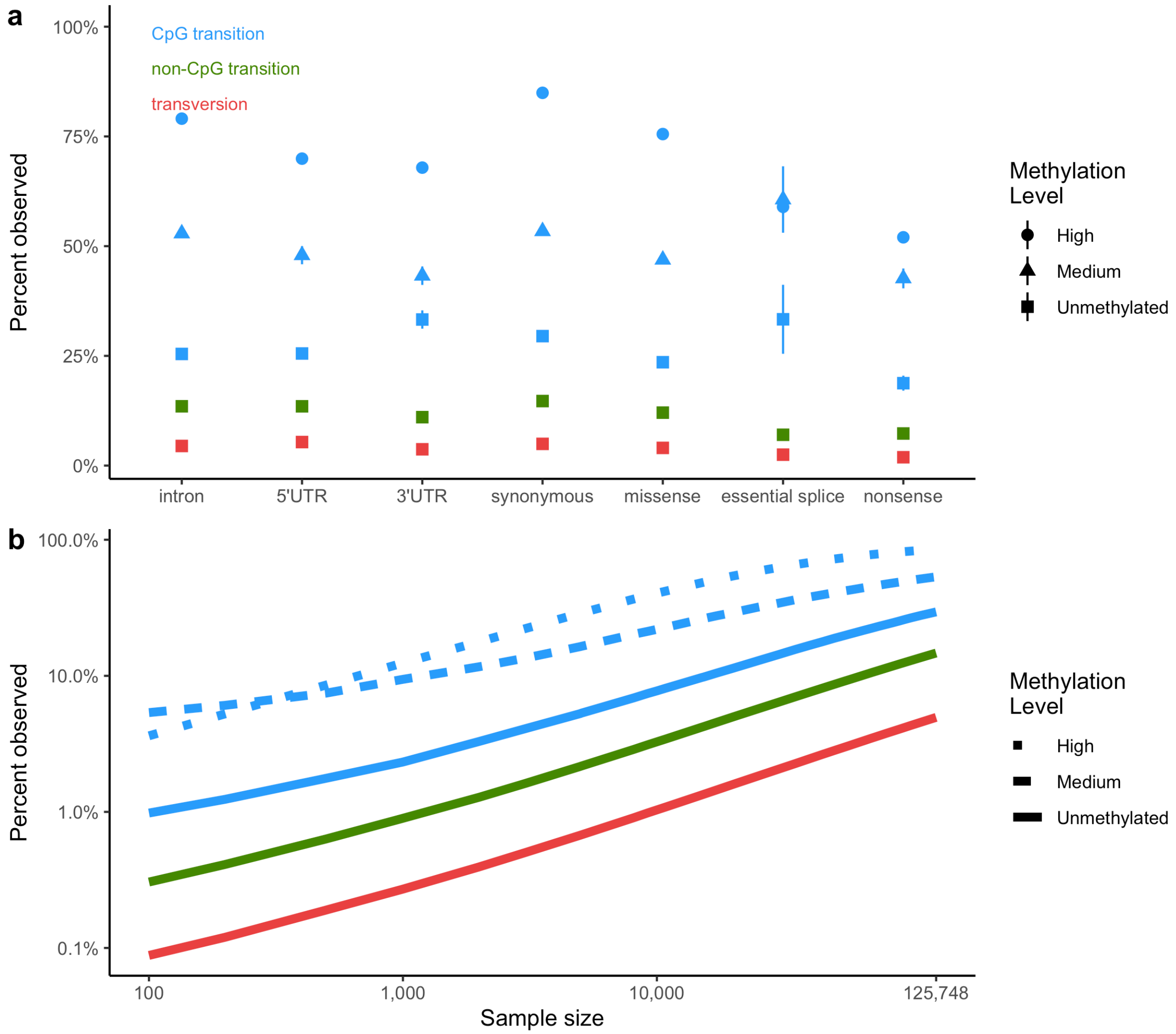


**Supplementary Figure 5 | Percent observed by methylation. a**, As in Fig. 1e, the proportion of possible variants observed for each functional class for each mutational type and methylation status in 125,748 exomes. **b**, As in Extended Data Fig. 4b, the proportion of possible variants observed as a function of sample size, broken down by variant class and methylation status. Colors are consistent in **a, b**.

#### *LOFTEE*

A number of challenges emerge when performing large-scale annotation of loss-of-function variants. Specifically, variants that are expected to have large effects on gene function will be depleted due to negative selection. However, error rates, including mapping, variant calling, and annotation errors, are relatively uniform[59]. Thus, this phenomenon results in an increased effective error rate at putative LoF variants[1,60].

To address these challenges, we implemented LOFTEE (the Loss-Of-Function Transcript Effect Estimator) as a plugin to VEP. LOFTEE utilizes the Ensembl API framework to annotate variant consequences based on properties of the genome and transcripts, and considers putative LoF (nonsense, splice-disrupting, and frameshift) variants in protein coding genes, filtering out variants with known annotation error modes (Extended Data Fig. 5a). Here, we use the MAPS metric to validate the expert-guided variant filters, and tune various parameters to ensure specificity. The putative LoF variants in these three classes of variation that are filtered out by LOFTEE exhibit MAPS scores near that of missense variants, suggesting that these filters are successfully filtering out annotation errors (Fig. 2b). The remaining variants have a MAPS score of 0.151, indicating a high level of deleteriousness.

One filter we specifically tuned based on this metric relates to variants near the end of a transcript (known as the END_TRUNC filter). Previously, we removed variants in the last 5% of a transcript, which was based on the enrichment of putative LoF variants in these regions[1,4]. With more fine-scaled metrics such as MAPS, we are now able to explore and optimize this filter. First, we implement the “50 base pair rule”, which removes any variants that are in the final exon or within 50 bp of the 3’ end of the penultimate exon[61]. This filter removes 39,072 (27.9%) of stop-gained variants in gnomAD, which collectively have a MAPS score of 0.106 (the remaining 100,626 variants maintain a very high MAPS score of 0.165). As terminal truncations may still be deleterious, we explored the properties of the variants that fail this filter based on the proportion of the transcript that is truncated, the number of base pairs they delete, and the number of base pairs deleted weighted by GERP score (Extended Data Fig. 5b), similar to the approach of Balasubramanian *et al.*[62]. Using the latter approach, we identify the top half of variants based on deleteriousness (cumulative GERP score of 180; MAPS = 0.141) while removing variants with MAPS = 0.06 (similar to missense variants).

Additionally, LOFTEE identifies additional putative LoFs in the form of non-canonical splice variants, including donor and acceptor disrupting variants, as well as variants that create donor splice sites. Briefly, we incorporate MaxEntScan[63] scores into a logistic regression model along with several other features related to splice site strength and evolutionary conservation. These are not used for assessment of constraint (see below), but are provided as other splice (OS) variants in the release file and have MAPS scores between those of missense and pLoF variants (Fig. 2b).

Crucially, LOFTEE favors a conservative approach to filtering variation with stringent filters to maximize specificity, as pLoF variants are enriched for error modes[1,60]. This results in a decreased sensitivity, as LOFTEE removes variants such as terminal truncations or at non-canonical splice sites, and does not consider other classes of LoFs such as missense variants that may affect structure or function or non-coding regulatory variants. Despite these caveats, after LOFTEE filtering, we discover 443,769 pLoF variants (Supplementary Table 15).

**Supplementary Table 15 | pLoF variants discovered in gnomAD**

| Filter applied (sequential) | Number of pLoF variants | | | |
|---|---|---|---|---|
| | Exomes | | Genomes | |
| Transcripts: | All | Canonical | All | Canonical |
| No filtering | 644,488 | 587,319 | 153,505 | 108,837 |
| High-quality variants | 515,326 | 470,169 | 128,275 | 108,762 |
| LOFTEE high-confidence (HC) | 443,769 | 413,097 | 101,288 | 91,033 |
| LOFTEE no flags | 385,842 | 364,759 | 85,051 | 78,559 |
| Call-rate filter (80%) | 345,458 | 332,495 | 83,919 | 77,551 |

*Genes affected by clonal hematopoiesis*

Clonal hematopoiesis of indeterminate potential (CHIP) is a phenomenon that is characterized by the accumulation of somatic mutations in mature blood cells derived from hematopoietic progenitors. The frequency of these mutations typically increases as an individual ages and complicate analyses of reference datasets[64]. To identify genes that show evidence of CHIP, we searched for canonical transcripts in which individuals carrying pLoF variants had a lower allele balance and greater age compared to those carrying synonymous variants. Cohorts vary in their reporting of age information. For example, some report age at diagnosis whereas others report the age at of the last patient visit. Age is therefore defined as the last known age of the individual and is not necessarily the age at sampling. We restricted our analysis to PASS-only variants in the exome dataset and required pLoF variants to be annotated as high confidence by LOFTEE. For each variant, the allele balance among individuals was represented as a frequency distribution with the values binned in increments of 0.05, ranging from 0 to 1.0. Age values were binned in increments of 5, ranging from 25 to 80, with values outside this range grouped into the respective flanking bin. Counts were summed across the frequency distributions for allele balance and age separately for pLoF and synonymous variants within a transcript in order to generate distributions for each gene, and the floor of the bin was used as the representative value of that bin. The Kolmogorov-Smironov test (KS test) was applied to determine if the allele balance and age distributions differed between pLoF and synonymous variants for each transcript. Mood's median test was also used to ensure that the medians of the distributions differed. We defined genes as having strong susceptibility to CHIP if the p-values of both the KS test and Mood's median test were below the Bonferroni-corrected p-value ($1.4 \times 10^{-6}$) and the median age of carriers of pLoF variants was at least 10% greater than the median age of those with synonymous variants.

Our analysis revealed evidence of CHIP in the canonical transcripts of three genes: *ASXL1*, *DNMT3A*, and *TET2* (Supplementary Table 16). These genes are among the most commonly reported CHIP-associated genes. Evidence of CHIP in these genes has been found in approximately 10% of individuals older than 65 years of age and 18% of individuals older than 90 years of age[65-67]. In support of this finding, we found that pLoF variants accumulate with age in these genes. For example, our analysis of *ASXL1* shows that the allele balance distribution is shifted left for pLoF variants, with a median of 0.25 compared to a median of 0.45 for synonymous variants, and the age distribution is shifted right for pLoF variants, with a median age of 70 as opposed to a median age of 55 for synonymous variants (Supplementary Fig. 6). The data for all genes with defined p-values for the allele balance and age KS and Mood's median tests can be found in Supplementary Dataset 6. The presence of somatic variants in such genes should be taken into account when interpreting the penetrance, pathogenicity, and frequency of potential germline variants. We focused our analysis on signals of pLoF variants though notably, CHIP can also be characterized by the accumulation of missense variants which would not have been revealed using our methods; future work to filter high-impact missense variants will enable a more complete understanding of CHIP.

**Supplementary Table 16 | Genes with evidence of CHIP**. Only genes with a p-value below the Bonferroni-corrected significance threshold of $1.4 \times 10^{-6}$ for both the two-sided KS test and Mood's median test, as well as a median age that is at least 10% greater for individuals with pLoF variants than those with synonymous variants, are shown. We define age as the last known age of the individual for 85,462 individuals.

| | **Gene** | ***ASXL1*** | ***DNMT3A*** | ***TET2*** |
|---|---|---|---|---|
| | **Canonical transcript** | ENST00000375687 | ENST00000264709 | ENST00000540549 |
| **Allele balance** | **Median value (pLoF)** | 0.25 | 0.25 | 0.30 |
| | **Median value (synonymous)** | 0.45 | 0.50 | 0.45 |
| | **KS test p-value** | 3.1E-69 | 1.3E-92 | 6.7E-109 |
| | **Mood's median test p-value** | 3.0E-19 | 1.7E-16 | 4.1E-24 |
| **Age** | **Median value (pLoF)** | 70 | 65 | 70 |
| | **Median value (synonymous)** | 55 | 55 | 50 |
| | **KS test p-value** | 2.0E-13 | 8.9E-11 | 1.3E-24 |
| | **Mood's median test p-value** | 2.8E-15 | 7.5E-12 | 3.1E-16 |

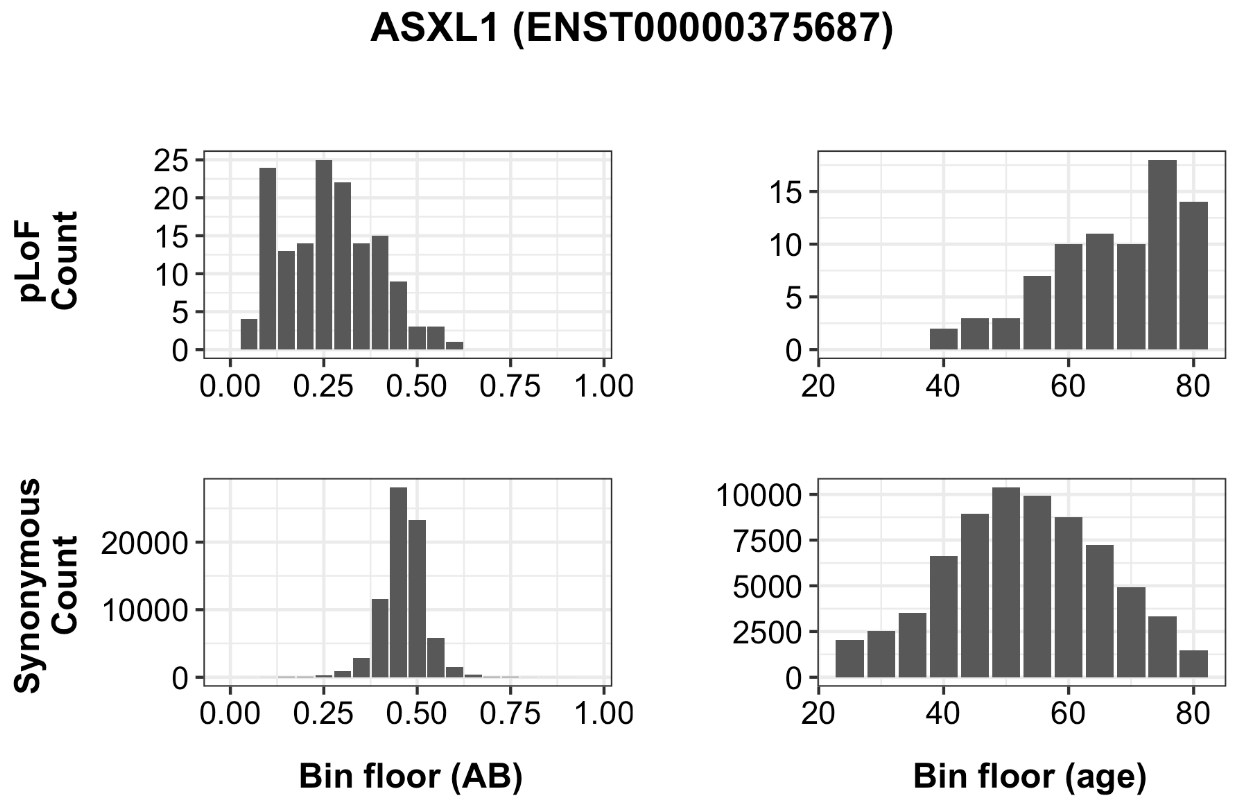


**Supplementary Figure 6 | Allele balance and age distribution for pLoF and synonymous ASXL1 variants.** Data for each distribution was binned, so the value of the bin floor was used as the representative bin value. All individuals less than 25 years old or greater than 80 years old were grouped into the 25 and 80 age bins, respectively. Information on age was available for 85,462 of the exome-sequenced individuals.

*Aggregate pLoF frequency*

In order to create a gene-level metric for the fraction of LoF haplotypes for each gene, we filtered the pLoF variants to those with high call-rate (80%), as variants with low call-rate inflate the allele frequency (e.g. a singleton where only 10 individuals were called will have a frequency of 5%, where the true frequency is likely lower), and used the most stringent set of LOFTEE criteria, which resulted in 345,458 variants. In order to avoid double counting LoF haplotypes harboring more than one pLoF, we group the data by gene and compute the fraction of individuals that have **no** pLoF variants, $q^2$. The fraction of haplotypes with a pLoF is then given by $p = 1 - \mathrm{sqrt}(q^2)$, the distribution of which across genes is shown in Extended Data Fig. 5c. This metric may be slightly affected by the undercalling of homozygotes issue as noted above, but it will only be slightly underestimated at genes with already high pLoF frequencies, as it primarily affects common variants.

*Homozygous variant curation*

From the 345,458 variants with the most stringent set of LOFTEE criteria (no filters or flags), we further filtered to 4,238 variants where at least one homozygous individual was observed. We subjected these variants to extensive manual curation, in order to filter technical errors commonly found in homozygous LoF variant prediction. These technical errors comprise three main groups: technical errors, rescue events, and transcript errors. Combinations of errors detected within these categories were used to determine if a variant was likely to ablate gene function. After reviewing each variant for technical artifacts, the variant was scored using a five point scale: not LoF, likely not LoF, uncertain, likely LoF, and LoF.

Technical errors included mapping errors and genotyping errors from sequencing issues, as well as misalignment of reads that could be detected in IGV and the UCSC genome browser, and errors within the reference sequence. Mapping errors are evident when reads around the variant harbored many other variants, especially those with abnormal allele balances. Furthermore, UCSC tracks for large segmental duplications, self chain alignments, and simple tandem repeats were utilized in determining mapping error status. Genotyping errors were partially eliminated by upstream filtering for read depth, genotype quality, and allele balance (see above). Additional hallmarks for genotyping errors included homopolymer repeats (defined as an insertion or deletion within or directly neighboring a sequence of seven or more of the same nucleotide), GC rich regions, and repetitive regions in which sequencing errors would be more common.

Rescue events include multi-nucleotide variants (MNVs), frame-restoring indels, and essential splice site rescues. MNVs visually identified in IGV and resulting in incorrectly called stop-gained mutations were classified as not LoF. Frame-restoring indels were verified by counting the length of the insertions and deletions to determine if the resulting variation disrupted the frame of the gene. The window used to detect surrounding indels was approximately 80 bp in length. Lastly, splice site rescues were verified by visually inspecting the +/- 21 bp region for an inframe splice site that could rescue the essential splice site. All possible in-frame splice site rescues within 21 bp of the essential splice site were filtered using Alamut (v.2.11), an alternative splice site prediction tool. Splice sites were classified as rescues if the MaxEntScan score for the alternate sequence was >/= 50% of the reference sequence score.

Finally, transcript errors were described as variants that occur in an exon found in a minority of transcripts for that gene or that occur in a poorly conserved exon. The UCSC genome browser was used to detect both of these situational errors. These were further assessed using pext (proportion expressed across transcripts) scores[15] and only those with low overall expression relative to the gene were determined to be not LoF. For an exon to be considered in a minority of transcripts, it had to be present in 50% or fewer of that gene's coding Gencode v19 Basic transcripts. Exon conservation was determined by looking at the nucleotide bp conservation based on PhyloP.

In order for a variant to be considered as LoF, it had to have no major error modes selected (such as LoF rescue) or have error modes such as weak exon conservation and minority of transcripts with a maximum pext score for the gene. If a single minor error mode was noted for a variant, which include some genotyping or mapping errors, it would be classified as likely LoF. In contrast, rescue errors were automatically classified at likely not LoF or not LoF. Multiple error modes (>= 3) resulted in a "not LoF" curation of the variant. Variants in which there was inconclusive evidence supporting the variant as LoF or not LoF were curated as unknown.

This process resulted in 2,636 homozygous pLoF passing curation filters, resulting in a list of 1,815 genes where we observe at least one homozygous knockout individual. This list likely misses some genes due to the strictness of curation and slight undercalling of homozygotes[4], but also may overestimate the effect of some pLoFs due to rescue mechanisms, and thus represents the best current estimate of confidently LoF-tolerant genes based on the gnomAD dataset. The full list of genes is provided in Supplementary Dataset 7. Next, we computed the mean number of pLoF alleles per individual, which is shown in Supplementary Table 17, with more detail broken down by population and frequency bin in Supplementary Datasets 8 and 9.

**Supplementary Table 17 | pLoF alleles per individual.** Variants with >95% frequency were removed. Filters are applied sequentially, except for where the three types of pLoF variants are broken down for rare variants.

| **Filter applied (sequential)** | **Exomes** | | **Genomes** | |
|---|---|---|---|---|
| **Zygosity:** | **Heterozygous** | **Homozygous** | **Heterozygous** | **Homozygous** |
| High-quality + LOFTEE HC | 148.2 | 25.5 | 205.3 | 33.2 |
| LOFTEE no flags | 92.7 | 15.5 | 135.6 | 21.1 |
| LCRs removed | 85.3 | 13.4 | 122.4 | 17.9 |
| Manual curation | 63.3 | 9.12 | 86.4 | 12.5 |
| Rare (<= 1% frequency) | 14.2 | 0.094 | 16.3 | 0.081 |
| • *Stop-gained* | *5.04* | *0.033* | *5.56* | *0.026* |
| • *Essential splice* | *4.00* | *0.028* | *4.25* | *0.026* |
| • *Frameshift* | *5.15* | *0.033* | *6.51* | *0.028* |
| Singleton or doubleton | 2.72 | 0.003 | 4.78 | 0.003 |
| Singleton | 1.95 | - | 3.62 | - |

### Constraint modeling

Konrad J. Karczewski, Kaitlin E. Samocha, Daniel G. MacArthur, Benjamin M. Neale, Mark J. Daly

In order to compute which genes are depleted of genetic variation, we extend the models described previously[4,7,49]. Briefly, we estimate the mutation rate for each single nucleotide substitution with 1 base of context (e.g. ACG -> ATG) using non-coding regions of the genome. We calibrate this mutation rate against the proportion observed of each context at synonymous sites in the exome, with an adjustment for low coverage regions. We apply these models to other classes of variation to establish an expected number of variants.

In these analyses, we sought to correct for the effect of methylation on the mutation rate at CpG sites, which become saturated for mutation at sample sizes above approximately 10,000 genomes[4]. To this end, we obtained methylation data for 37 tissues from the Roadmap Epigenomics Consortium[54]. Across these tissues, we compute the mean proportion of whole genome bisulfite sequencing reads corresponding to the methylated allele at each CpG site (Extended Data Fig. 6a) and discretize this metric into >0.6, 0.2-0.6, <0.2 (or missing) as high, medium, and low methylation levels, which are then used for all future analyses.

##### *Mutational model*

To calculate the baseline mutation rate for each substitution and context (and for CpG sites, the methylation level), we count the instances of each trinucleotide context in the autosomes of the human genome where 1) the most severe annotation was intron_variant or intergenic_variant, 2) the GERP score was between the 5th and 95th percentile of the genome-wide distribution (between -3.9885 and 2.6607), and 3) the mean coverage in the gnomAD genomes was between 15X and 60X. As methylated CpG variants are saturated at sample sizes above ~10,000 genomes, we downsampled the dataset to 1,000 genomes for use in calculating the mutation rate. Further, sites were removed if they were found in the gnomAD dataset but filtered out due to low quality, or found in greater than 5 copies in the downsampled set. This resulted in 5,918,128,813 possible variants, at which 23,930,773 high-quality variants with 5 or fewer copies were observed in the downsampled set. From these values, we compute the proportion observed for each context, which represents the relative mutability of each variant class, and scale this factor so that the weighted genome-wide average is the human per-base, per-generation mutation rate ($1.2 \times 10^{-8}$) to calculate the absolute mutation rate. These mutation rate estimates are well-correlated with previous estimates at non-CpG sites, but crucially, now incorporate the effect of methylation on mutation rate (Extended Data Fig. 6b). These estimates are provided in Supplementary Dataset 10.

*Improvements to constraint model*

Using these mutation rate estimates, we compute the expectation for number of variants in a given functional class as follows. First, because the exome dataset has a substantially larger sample size, we calibrate the mutation rates to a relatively neutral class of variation, synonymous variants. For each possible site where the most severe consequence on a canonical transcript is synonymous_variant, we compute the proportion observed in a similar fashion as above: we remove possible variants where there was no coverage, a low-quality variant was observed, or a variant above 0.1% frequency was observed. For each substitution, context, and methylation level, further divided by median exome coverage (at integer values between 1-100), we compute the proportion observed of high-quality variants below 0.1% frequency. Considering only sites above a median coverage of 40, we correlate the proportion observed with the mutation rates previously obtained for each mutational class (Extended Data Fig. 6c). We fit two models, one for CpG transitions and one for the remainder of sites (transversions and non-CpG transitions), to calibrate from the mutation rate to proportion observed in 125,748 exomes (Extended Data Fig. 6d).

For sites with a median coverage of 1-39, we perform an additional coverage correction, in a similar fashion as previously described[4], but performed at base-level resolution rather than exon-level. First, we define a metric that represents the relative mutability of the exome, or the number of observed synonymous variants divided by the total number of possible synonymous variants times the mutation rate summed across all substitutions, contexts, and methylation level. We compute this metric for high coverage sites as a global scaling factor, and divide this metric at low coverage sites by this scaling factor to create an observed:expected ratio for a given coverage level (Extended Data Fig. 6e). We build a model of log10(coverage) to this scaled ratio as a correction factor for low coverage sites.

Using these models, we can compute the expected number of variants for an arbitrary set of substitutions, such as all the pLoF variants in a given transcript. To do so, for each substitution, context, methylation level, and median coverage, we sum the number of possible variants times the mutation rate for all variants in our class of interest, and apply the calibration model separately for CpG transitions and other sites. For sites with median coverage from 1-39, we multiply this value by the coverage correction factor; otherwise, we use the value as-is. These values are summed across the set of variants of interest to obtain the expected number of variants. After removing TTN, we observe a good fit for synonymous variants at $r = 0.98$ (Extended Data Fig. 6f) and depletion for missense and pLoF variants (Extended Data Fig. 6g-h), consistent with previous results[1]. 392 genes had a poor fit of synonymous variation ($z < -3.71$), which are enriched for mapping artifacts: approximately 32% of these (126/392) have a mappability score < 0.9 (as described above in “Comparison of whole-exome and whole-genome coverage in coding regions”), compared to 10% (1908/18839) of genes that are not outliers for number of synonymous variants. Other genes in this category include the highly-paralogous HIST1 complex, as well as genes notorious for mapping errors such as *FLG*, *AHNAK2*, and *MUC* genes.

This base-level calibration and coverage correction process enables the assessment of constraint against any arbitrary sets of variants. We apply this method for all pLoF variants labeled as high-confidence by LOFTEE. While not within the scope of the current paper, we additionally assess constraint against missense variants, and missense variants annotated as probably_damaging by PolyPhen2[68] for the benefit of users. We compute these metrics separately for all 80,950 transcripts, of which 79,174 have an expected number of pLoF variants > 0, which are provided in the release. For most downstream analyses, unless otherwise specified, we consider the canonical transcript of each gene and compute these metrics for 19,704 genes, of which 19,197 have an expected number of pLoF variants > 0.

For pLoF variants, we compute the pLI score as previously described [4], and we further compute the 90% confidence interval around the observed:expected ratio. Specifically, for a given pair of observed and expected values, we compute the density of the Poisson distribution with fixed k (the observed number of variants) over a range of lambda values, which are given by the expected number of variants times a varying parameter ranging between 0 and 2. The cumulative density function of this function is computed and the value of the varying parameter is extracted at points corresponding to 5% and 95% to indicate the bounds of the confidence interval. The upper bound of this interval is termed the LoF observed/expected upper bound fraction (LOEUF), and is used for most analyses in this manuscript. All constraint and summary metrics are provided in Supplementary Dataset 11.

*Summary of constraint metrics*

The distribution of the pLoF observed/expected ratio is shown in Extended Data Fig. 7a: the mean observed/expected value is 0.537 (median 0.482), with 1266 genes with a value of 0 (no pLoFs observed). However, as we expect fewer than 5 pLoF variants in 498 of these genes, we instead use the LOEUF score described above (Extended Data Fig. 7b), which has a mean of 0.952 (median 0.911). Binning LOEUF into deciles partitions all human genes by the number of observed and expected pLoFs (Extended Data Fig. 7c), resulting in ~1,920 genes per decile.

LOEUF is correlated with coding sequence length (beta = $-1.07 \times 10^{-4}$; $p < 10^{-100}$; Extended Data Fig. 7d): as a result, we have adjusted for gene length or removed genes with fewer than 10 expected pLoFs in all analyses. The most constrained decile has an aggregate pLoF observed/expected ratio of ~6%, and accordingly, other splice and missense variants are also depleted, to a lesser degree (Extended Data Fig. 7e). The genes previously described as high pLI genes[4] are more likely to fall in the most constrained deciles (Extended Data Fig. 7f), while unconstrained genes are more likely to harbor homozygous pLoFs (Extended Data Fig. 7g). Finally, LOEUF decile correlates with the aggregate pLoF frequency ($\rho = 0.157$; $p < 10^{-100}$; Extended Data Fig. 7h). A comparison of these metrics is shown in Supplementary Fig. 7.

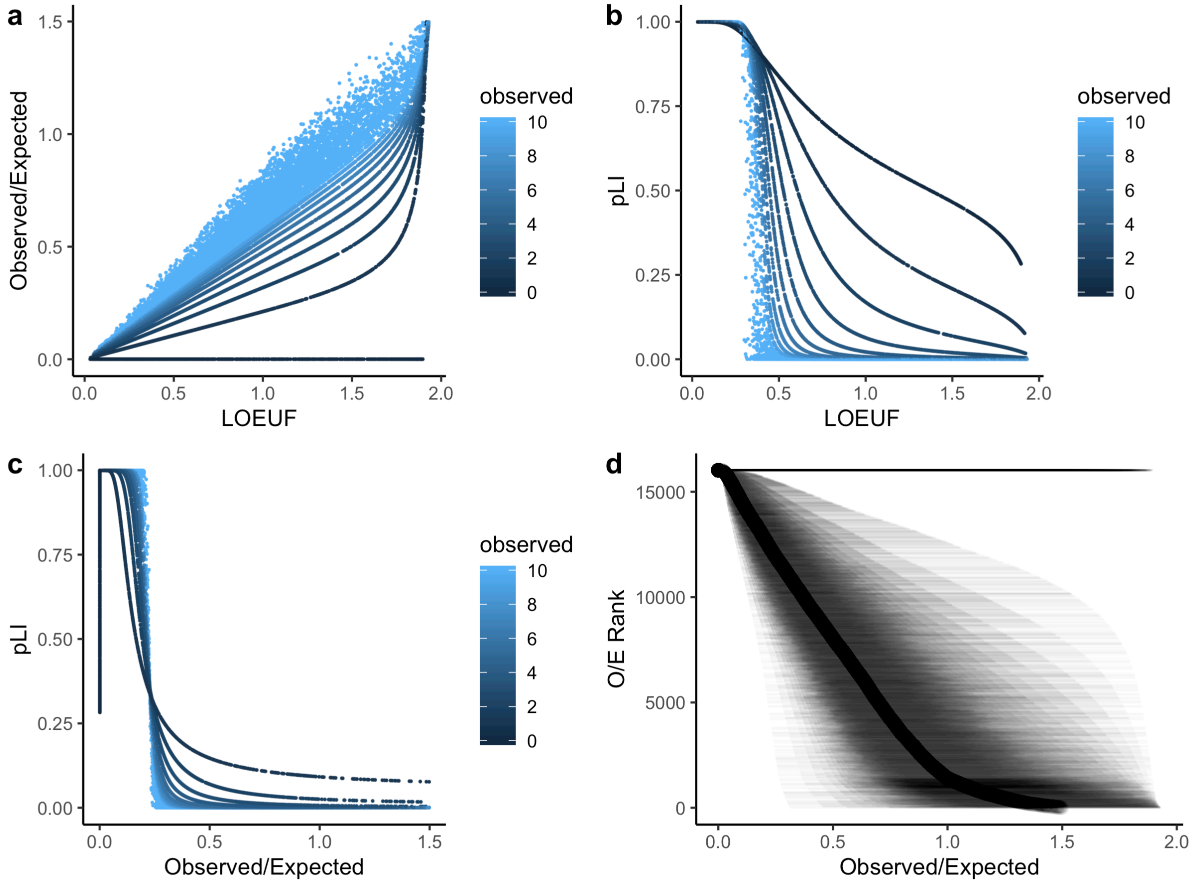


**Supplementary Figure 7 | LOEUF summaries.** Scatter plots of the observed/expected (o/e) ratio vs LOEUF (a), pLI vs LOEUF (b), and pLI vs observed/expected (c). d, The o/e ratio (dots) for each gene ranked by o/e, with confidence intervals as thin lines (LOEUF is the upper bound).

For certain analyses in this manuscript, we filter the dataset to genes where we expect over 10 pLoF variants. This cutoff was chosen as the minimum number of expected pLoF variants that can result in membership in the most constrained bin (11.1 expected) or pLI > 0.95 (9.43 expected). At present, 72.1% of genes (13841/19197) have > 10 pLoFs expected, including 86.5% of disease-associated genes from OMIM (2888/3340; OR = 0.45; Fisher's $p < 10^{-100}$). Of the 59 genes satisfying ACMG criteria for reporting of secondary findings, only five are underpowered, or have fewer than ten pLoFs expected (*SDHD*, *MYL3*, *VHL*, *MYL2*, *SDHAF2*).

We computed the expected number of variants for each gene, using the process described above, repeated for each downsampling of the exome dataset. For each gene, the number of individuals required to achieve a given expected number of variants is extrapolated using a linear model of log(number of expected variants) ~ log(number of individuals), which is available in Supplementary Dataset 12. The proportion of genes where at least a given number pLoF and missense variants are expected is plotted for a range of sample sizes in Supplementary Fig. 8.

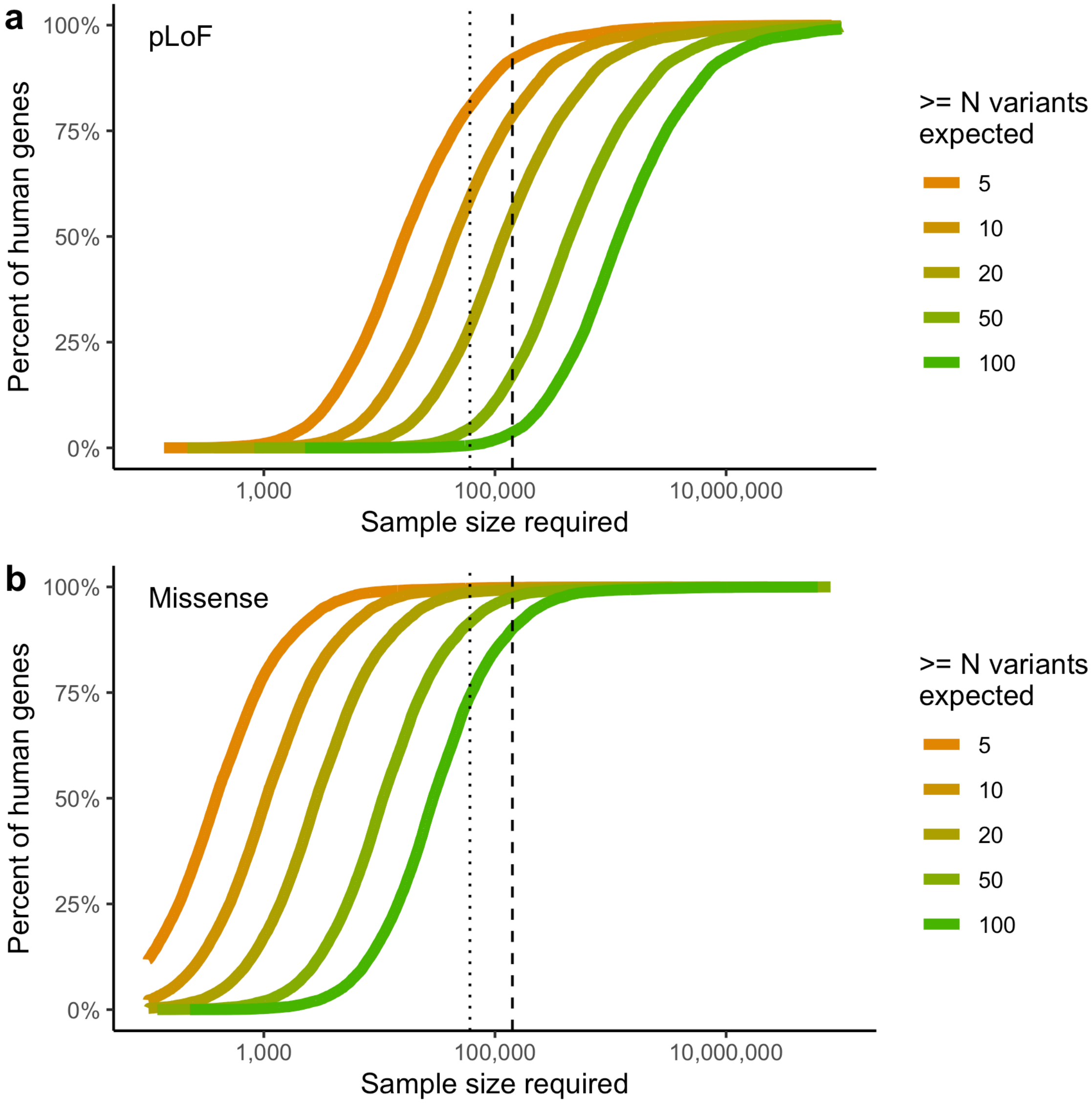


**Supplementary Figure 8 | The sample size required for well-powered constraint calculations.** The proportion of genes where a varying number of pLoF (a) and missense (b) variants would be expected (under neutrality) is shown as a function of (log-scaled) sample size.

### Constraint assessment and implications

Konrad J. Karczewski, Beryl B. Cummings, Daniel Rhodes, Qingbo Wang, Ryan L. Collins, Benjamin M. Neale, Mark J. Daly, Michael E. Talkowski, Daniel G. MacArthur

We assessed the LOEUF metric using a number of orthogonal metrics, including membership in known disease gene lists, comparisons to structural variant occurrence, and lethality in mouse orthologs and cellular knockouts. We explore the biological properties of constrained genes, as well as features of constraint across populations and subsets.

#### *Gene list comparisons*

First, we compared the LOEUF distribution of various established gene sets (https://github.com/macarthur-lab/gene_lists), as previously used and described[4]. In particular, we used a curated list of 189 known haploinsufficient disease genes, 709 autosomal dominant disease genes, 1183 autosomal recessive disease genes, and 360 olfactory receptors. For each gene list, we counted the number of genes in that gene list in each LOEUF decile (Supplementary Table 18) and normalized to the number of genes in the list (Fig. 3a).

**Supplementary Table 18 | Gene list membership by LOEUF decile**

| LOEUF decile | Haplo-insufficient | Autosomal Dominant | Autosomal Recessive | Olfactory Genes |
|---:|---:|---:|---:|---:|
| 0-10% | 104 | 140 | 36 | 0 |
| 10-20% | 47 | 128 | 72 | 1 |
| 20-30% | 17 | 86 | 112 | 0 |
| 30-40% | 8 | 80 | 173 | 4 |
| 40-50% | 7 | 65 | 206 | 8 |
| 50-60% | 4 | 54 | 207 | 6 |
| 60-70% | 0 | 46 | 154 | 18 |
| 70-80% | 2 | 49 | 120 | 49 |
| 80-90% | 0 | 34 | 58 | 96 |
| 90-100% | 0 | 26 | 40 | 174 |

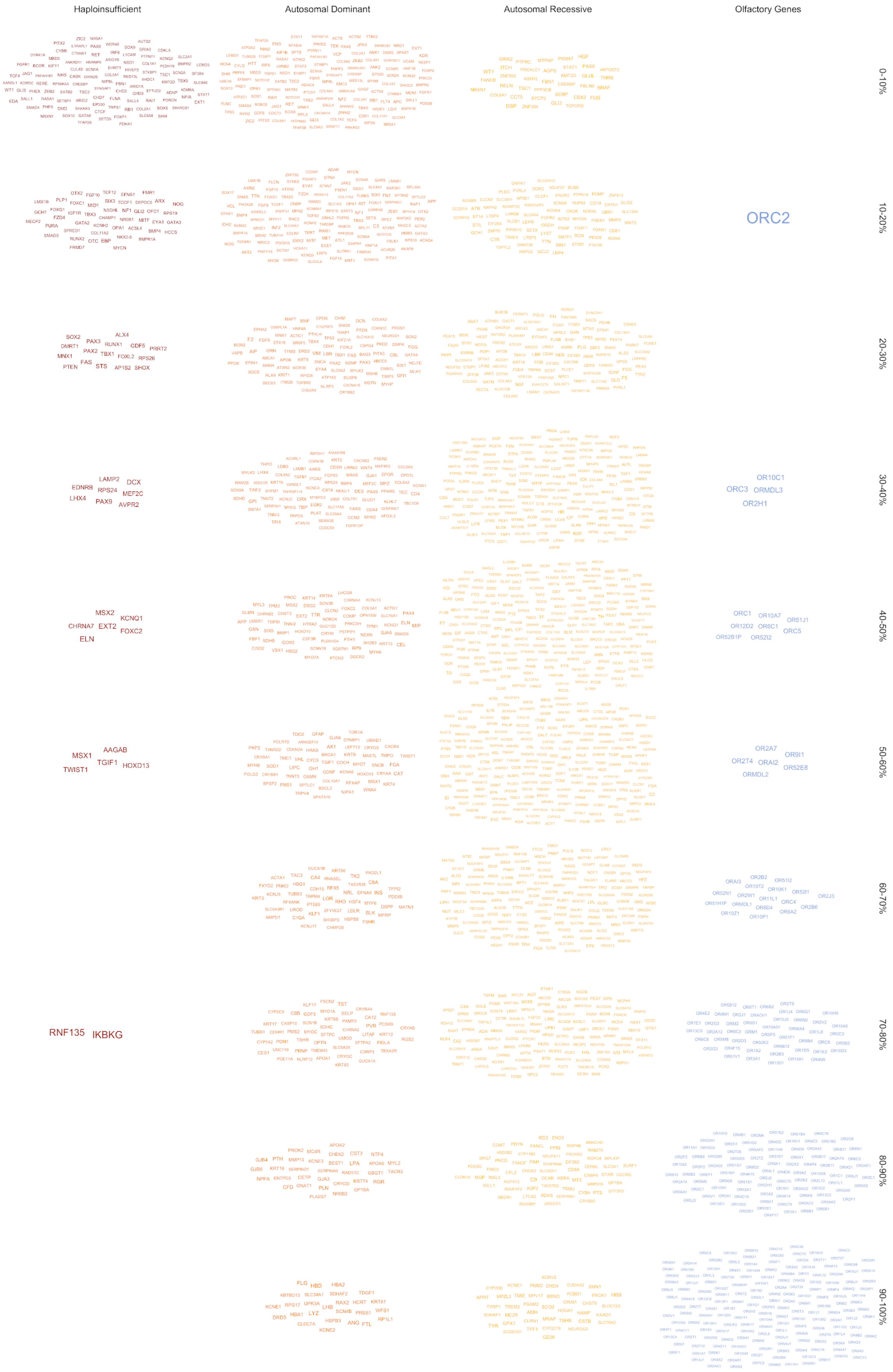

Haploinsufficient
Autosomal Dominant
Autosomal Recessive
Olfactory Genes
0-10%
10-20%
20-30%
30-40%
40-50%
50-60%
60-70%
70-80%
80-90%
90-100%
ORC2
RNF135
IKBKG

**Supplementary Figure 9 (previous page) | Genes within gene lists by LOEUF decile.** The genes represented in Supplementary Table 15 are shown by gene list and LOEUF decile.

Supplementary Fig. 9 enumerates all the genes in Supplementary Table 18, which are also available as Supplementary Dataset 13. Of the haploinsufficient genes, 80% were found in the two most constrained deciles of the genome. There were two genes that are in the haploinsufficient gene list, but with little evidence of constraint (in the 8th decile): *RNF135* (LOEUF = 1.44), which has limited support for pathogenicity[69]; and *IKBKG* (LOEUF = 1.37), which is poorly covered in gnomAD and whose first exon is lowly expressed, suggesting that the pLoFs in this gene are likely false positives. Membership in the haploinsufficient gene class is highly predicted by LOEUF (logistic regression beta = -4.3; p = $1.57 \times 10^{-33}$), even when adjusted for coding sequence length (p = 0.18 for the contribution of gene length in the joint model). Likewise, membership in the olfactory gene class is positively correlated with LOEUF (logistic regression beta = 3.4; p = $2.5 \times 10^{-85}$), even when adjusted for gene length (p = 0.023 for the contribution of gene length in the joint model).

*Structural variant comparisons*

We also compared constraint metrics to the relative enrichment or depletion of predicted LoF deletion SVs documented by a companion study[11]. We restricted this analysis to autosomal, biallelic, rare (AF<1%) deletions found in the 6,749 samples with WGS data in this manuscript that were predicted to overlap at least one nucleotide from a protein-coding exon in a gene's canonical transcript per Gencode v19[70], and retained all genes with matching gene symbols between the SNV and SV datasets (n=17,604). Despite the SV dataset being substantially sparser than SNVs and indels on a per-gene basis, we nevertheless fit a Poisson regression model to predict the number of expected rare biallelic LoF deletions per gene based on the following covariates: gene length, number of exons, median exon size, total number of nonredundant nucleotides in protein-coding exons, number of introns, median intron size, total number of nonredundant nucleotides in introns, and annotated overlap with segmental duplications. To prevent existing signatures of strong purifying selection confounding this null model of expected LoF SV per gene, we restricted the dataset to genes in the 5th-9th deciles based on observed:expected ratios for LoF SNVs when fitting the model. The SV-derived observed:expected ratios are correlated with LOEUF ($r = 0.13$; $p = 3.5 \times 10^{-71}$), after adjusting for gene length ($p = 7.5 \times 10^{-6}$ for the contribution of gene length). Finally, to compare these SV-derived observed:expected values to SNV constraint metrics, we first binned genes based on their LOEUF decile before summing observed and expected rare LoF deletion SV counts for all genes per decile and computing a decile-level LoF SV observed:expected ratio from these sums.

##### *Mouse and cell model comparisons*

We compared our constraint metrics with evidence of essentiality from mouse and human cell knockout experiments. For comparisons to mouse, human orthologs of mouse heterozygous lethal genes were defined by the following processes, based on curated experimental results from Mouse Genome Informatics (MGI, http://www.informatics.jax.org). First, we extracted all the mouse genes where lethality upon heterozygous knockout was ever reported in MGI, using the MouseMine web interface (http://mousemine.org/; accessed on 12 Feb 2017). Specifically, mouse heterozygous lethal genes were defined as those mouse genes containing the subset of the Mammalian Phenotype Term “abnormal survival” (MP: 0010769), “nervous system phenotype” (MP:0003631), and “embryo phenotype” (MP:0005380), which are highly associated with embryonic lethality (Supplementary Table 19) in the heterozygous knockout state, for any mouse strain. Next, we mapped human genes to their corresponding mouse orthologs by extracting the human-mouse ortholog correspondence table from MGI (http://www.informatics.jax.org/downloads/reports/HMD_HumanPhenotype.rpt). For each of the human genes, if any of the mouse homolog(s) was defined as heterozygous lethal, we defined the corresponding human gene as mouse heterozygous lethal, for a total of 389 genes.

**Supplementary Table 19 | Mammalian Phenotype Term lists**

| Mammalian Phenotype ID | Term | parent term | parent Mammalian Phenotype ID |
|---|---|---|---|
| MP:0011400 | lethality, complete penetrance | mortality/aging | MP:0010768 |
| MP:0010831 | lethality, incomplete penetrance | mortality/aging | MP:0010768 |
| MP:0002081 | perinatal lethality | mortality/aging | MP:0010768 |
| MP:0002082 | postnatal lethality | mortality/aging | MP:0010768 |
| MP:0002080 | prenatal lethality | mortality/aging | MP:0010768 |
| MP:0010770 | preweaning lethality | mortality/aging | MP:0010768 |
| MP:0008569 | lethality at weaning | mortality/aging | MP:0010768 |
| MP:0001890 | anencephaly | nervous system phenotype | MP:0003631 |
| MP:0000914 | exencephaly | nervous system phenotype | MP:0003631 |
| MP:0001730 | embryonic growth arrest | embryo phenotype | MP:0005380 |
| MP:0009657 | failure of chorioallantoic fusion | embryo phenotype | MP:0005380 |
| MP:0001683 | absent mesoderm | embryo phenotype | MP:0005380 |
| MP:0001696 | failure to gastrulate | embryo phenotype | MP:0005380 |
| MP:0001690 | failure of somite differentiation | embryo phenotype | MP:0005380 |
| MP:0004180 | failure of initiation of embryo turning | embryo phenotype | MP:0005380 |
| MP:0009331 | absent primitive node | embryo phenotype | MP:0005380 |
| MP:0011185 | absent primitive endoderm | embryo phenotype | MP:0005380 |
| MP:0000932 | absent notochord | embryo phenotype | MP:0005380 |
| MP:0004388 | absent prechordal plate | embryo phenotype | MP:0005380 |
| MP:0001693 | failure of primitive streak formation | embryo phenotype | MP:0005380 |

Next, in order to compare the constraint metrics with human cell essentiality inferred by pooled-library screening experiments using CRISPR/Cas genome engineering, we used a list of 684 genes deemed essential, and 927 genes deemed non-essential for cell viability in multiple cultured cell lines such as HEK293T cells and K562 cells[24]. Specifically, Hart *et. al* defined a set of essential genes using a strict Bayes Factor threshold, corresponding to the posterior probability of being >90% essential for more than six cell lines out of a minimum of 7 to a maximum of 12 different screens in different cancer and immortalized cell lines. Additionally, they defined nonessential genes based on low RNA expression level across 17 different cell lines, as well as curated shRNA screening results, and this was validated with CRISPR/Cas screening.

Genome engineering using a CRISPR/Cas system is known to typically induce a biallelic mutation at the targeted locus[71], resulting in a homozygous loss of function of the corresponding gene. As there is no corresponding dataset that differentiates heterozygous from homozygous lethality, these 684 genes likely represent a mixture of genes that are lethal when knocked out in heterozygous and/or homozygous states.

For each of the three essentiality categories (mouse heterozygous lethal, human cell essential, and human cell non-essential), we matched the canonical gene name by HUGO Gene Nomenclature Committee (HGNC) approved symbol, binned all the genes in the category based on their LOEUF decile, and normalized the number by dividing by the total number of genes in that specific category. The percentage of genes per decile is shown in Fig. 3c,d, providing evidence that the constraint metric correlates well with essentiality as measured by mouse and human cell experiments. Overlap with mouse heterozygous lethality was significantly associated with LOEUF (logistic regression beta = -2.27; p = $3.3 \times 10^{-52}$), even when adjusted for coding sequence length (beta = $3.3 \times 10^{-5}$; p = 0.028). LOEUF is also correlated with cell essentiality (logistic regression beta = -1.71; p = $1.7 \times 10^{-65}$; coding sequence length: beta = $2.5 \times 10^{-4}$; p = $2.4 \times 10^{-12}$) and non-essentiality (beta = 1.45; p = $3.8 \times 10^{-71}$; coding sequence length: beta = $-5.9 \times 10^{-6}$; p = 0.84). Note that the stronger skew towards lower deciles in the mouse data is likely due to more specific targeting of heterozygous lethality, compared to the CRISPR screens.

Further, we defined mouse homozygous-lethal knockout genes in a similar fashion as described above for heterozygous genes. As expected, the genes that were tolerant of homozygous knockout in humans (see above: Homozygous Variant Curation) had a significantly lower probability of being lethal when knocked out in mouse or essential in human cells, and were on average less constrained (i.e. higher LOEUF scores; Supplementary Table 20).

**Supplementary Table 20 | Comparison of genes we observe homozygous deletion in gnomAD population with other gene lists.** Fewer homozygous knockout tolerant genes are included in this comparison (n=1519 vs 1649) as 130 genes did not have a unique gene symbol approved by HGNC. Further, we filtered out genes from the mouse and cell comparison sets that did not have LOEUF score. For gene set comparisons, the p-value was computed using a Fisher's exact test (two-sided) and for LOEUF comparisons, a t-test (two-sided) was used.

| | Mouse Heterozygous KO | | Mouse Homozygous KO | | Cell Essential | | |
|---|---|---|---|---|---|---|---|
| | Lethal | Others | Lethal | Others | Essential | Others | Mean LOEUF |
| Homozygous KO tolerant genes (n=1519) | 12 | 1507 | 87 | 1432 | 6 | 1513 | 1.26 |
| Remaining genes (n=17675) | 383 | 17292 | 3647 | 14028 | 677 | 16998 | 0.91 |
| Odds Ratio | | 0.36 | | 0.23 | | 0.10 | n/a |
| p-value | | $6.8 \times 10^{-5}$ | | $9.1 \times 10^{-57}$ | | $1.5 \times 10^{-17}$ | $< 10^{-100}$ |

#### *Functional categorization*

We assessed the correlation between the LOEUF metric and a proxy measure for biological knowledge, the target development level (TDL) from the Pharos database[72]. A full definition of the TDL and the associated categories can be found at https://pharos.nih.gov. In brief, gene-products can be categorised into one of four categories based on the drugs and small molecules that target them, Tclin - targets with approved drugs; Tchem - targets with drug activities in ChEMBL that are not approved for market; Tbio - targets with weaker drug activities that do not meet the required activity thresholds to be classified as Tchem; Tdark - targets about which little is known. Each of these categories is significantly correlated with LOEUF in a joint logistic regression model with coding sequence length: Tclin (beta = -0.78; $p = 4 \times 10^{-18}$; cds length: beta = $2 \times 10^{-6}$; p = 0.89), Tchem (beta = -0.63; $p = 8 \times 10^{-30}$; cds length: beta = $5 \times 10^{-6}$; p = 0.68), Tbio (beta = -0.99; $p < 10^{-100}$; cds length: beta = $1.6 \times 10^{-5}$; p = 0.07), Tdark (beta =

1.17; $p < 10^{-100}$; cds length: beta = $2.7 \times 10^{-5}$; p = 0.009). For each class, we counted the number of genes in the list in each LOEUF decile and normalized according to the number of genes in the list (Extended Data Fig. 8a).

*Network analysis*

Protein-protein interaction networks were used to compare the LOEUF metric to gene-product functional importance. The STRING database[73] was queried using the R API (STRINGdb, v1.22.0) for the protein-protein interactions of all genes with at least 10 expected pLoFs. We then filtered interactions based on their combined scores[74] such that only high confidence interactions (score > 0.7) remained. From this, we generated a directed acyclic graph and kept the largest component resulting in a protein-protein interaction network of 14,955 nodes (proteins) and 315,217 edges (interactions). We then calculated the degree (the number of nodes a node is connected to) of each node. Lastly, we binned proteins based on their gene LOEUF deciles and computed the within decile mean degree with 95% confidence intervals (Fig. 4a).

*Expression*

The GTEx v7 gene and isoform expression data were downloaded from dbGaP (https://www.ncbi.nlm.nih.gov/gap) from accession phs000424.v7.p2.c999. The GTEx pipeline for isoform quantification is available publically (https://github.com/broadinstitute/gtex-pipeline/) and briefly involves 2-pass alignment with STAR v2.4.2a and isoform quantification with RSEM v1.2.22. We calculated the median isoform expression (measured as transcripts per million, or TPM) across individuals for all GTEx tissues, except reproduction-associated GTEx tissues (endocervix, ectocervix, fallopian tube, prostate, uterus, ovary, testes, vagina), cell lines (transformed fibroblasts, transformed lymphocytes) and any tissue with less than one hundred samples (bladder, brain cervicalc-1 spinal cord, brain substantia nigra, kidney cortex, minor salivary gland), resulting in the use of 38 GTEx tissues, which was used for subsequent downstream analysis.

We then computed the number of tissues (up to the 38 tissues as described above) where the transcript is expressed (defined as TPM > 0.3). We show the distribution of number of tissues where the canonical transcript is expressed in each (gene-based) LOEUF decile (Fig. 4b). Overall, the number of tissues in which a canonical transcript is expressed is correlated with LOEUF (linear regression beta = -1.07; $p < 10^{-100}$) when adjusted for gene length (beta = -9.9 x $10^{-4}$; $p = 10^{-53}$ for the contribution of gene length). Additionally, we merged these values with the per-transcript LOEUF table, where we computed the LOEUF transcript decile by binning the 79,174 transcripts with expected pLoF variants > 0 into 10 bins. We show the number of tissues where each transcript is expressed by its LOEUF transcript decile, broken down by canonical transcript and all transcripts (Extended Data Fig. 8b). Similarly, the number of tissues in which a transcript is expressed is correlated with the transcript's LOEUF (linear regression beta = -5.2; $p < 10^{-100}$) when adjusted for gene length (beta = -9.4 x $10^{-5}$; p = 0.01 for the contribution of gene length).

In order to investigate differential constraint within a gene, we identified 1,790 genes containing at least one constrained (transcripts belonging to the first decile) and one unconstrained transcript (transcripts belonging to any other decile). We divided the sum of the expression of constrained transcripts by the sum expression of all transcripts in the gene, and show that the constrained transcript accounts for most of a gene's expression (Fig 4c).

Finally, we considered whether the most expressed transcript in a disease-relevant tissue was also the most constrained for each gene, using a data set of gene to tissue mappings based on disease annotations[25]. After collapsing GTEx tissues to match with the gene-tissue dataset (taking the max expression across GTEx tissues where multiple tissues matched), we determined the transcript with the highest expression for each gene. For each gene, we filtered the expression data to only include the tissues where at least one disease was identified[25]. In this dataset, transcripts that were the most expressed in these tissues were also be the most constrained transcript for that gene in 55.3% of genes (a 5.4-fold enrichment over background). As an enrichment is expected by chance (e.g. due to single transcript genes always being the most constrained and most expressed), we performed a permutation test with 10000 replicates, sampling which transcript was most constrained with replacement, which resulted in a background mean of 28.8% (1.91-fold enrichment; Extended Data Fig. 8c).

*Population-specific constraint modeling*

The constraint modeling process was repeated for each of the down-samplings (described above in Frequency and context annotation) for the full dataset and each population in gnomAD, only considering variants present in each population with frequency < 0.1%, in order to compute the number of observed and expected pLoF variants for each sample of individuals (downsampling analysis shown in Fig. 2c,d).

In order to compare across populations, we considered only the results for 8,128 individuals from each population (corresponding to the sample size of the smallest sampled population, African/African-American individuals). We compared the LOEUF scores from non-Finnish European individuals to those from African/African-American individuals for 927 genes where the expected number of pLoF variants was at least 10 in each population (Extended Data Fig. 8d). We find that on average, the LOEUF score is lower (more constrained) in the African/African-American population than in the non-Finnish European population (0.488 vs. 0.617; t-test $p = 4 \times 10^{-14}$), but the two are highly correlated ($r = 0.78$; $p < 10^{-100}$). In 865 genes where the expected number of pLoF variants is at least 10 in all 5 major continental populations, we find a dependence between population and mean LOEUF score (Extended Data Fig. 8e).

*Comparison to previous metrics of essentiality*

We compared LOEUF to previous metrics of genic essentiality, including pLI and RVIS. pLI was computed on the gnomAD exome variants in this manuscript as described previously[4] and RVIS[8] scores for gnomAD were downloaded from http://genic-intolerance.org/ (RVIS_Unpublished_ExACv2_March2017.txt downloaded on July 15, 2019). We selected two gold standard datasets for comparison: 1) the haploinsufficient gene list described in “Gene list comparisons”, and 2) a union of the mouse heterozygous lethal and “cell essential” gene lists described in “Mouse and cell model comparisons.” Using these genes as “true positives” and all other genes as “negatives,” we performed logistic regression and created receiver operator characteristic (ROC) curves for each method and computed the area under the curve (AUC) as a performance assessment. In the logistic regression, LOEUF is highly correlated with membership in the haploinsufficient (beta = -2.6; p = 4 x $10^{-5}$) and essential (beta = -1.4; p = 1.9 x $10^{-25}$) gene lists, in a joint model with pLI (beta = 1.5; p = 3 x $10^{-4}$ and beta = 0.17; p = 0.15, respectively), RVIS (beta = -0.18; p = 0.05, and beta = -0.19; p = 1.5 x $10^{-5}$, respectively), and coding sequence length (beta = 4 x $10^{-6}$; p = 0.92 and beta = -8 x $10^{-5}$; p = 7 x $10^{-4}$, respectively). LOEUF substantially outperforms RVIS for both gold standard sets, and performs similarly to pLI for identifying haploinsufficient genes and outperforms pLI for essential genes (Supplementary Fig. 10).

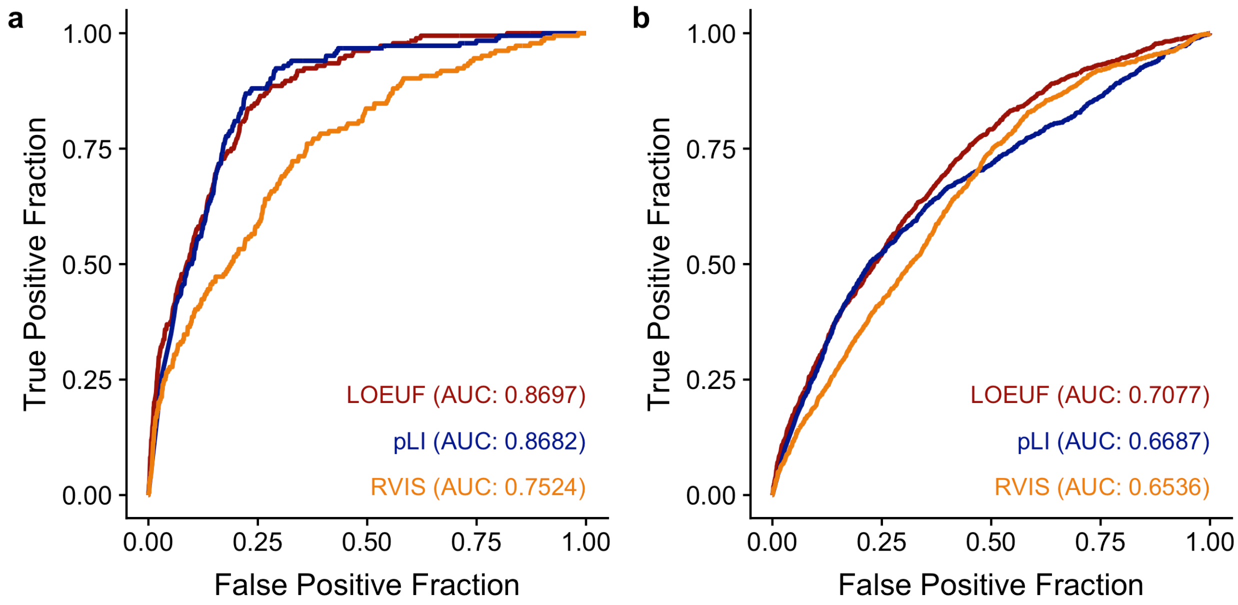

**Supplementary Figure 10 | Comparison to other gene essentiality metrics.** ROC curves for each gene essentiality metric, for discerning 184 haploinsufficient genes from 16,714 background genes **(a)** or 1,019 mouse heterozygous lethal or cell essential genes from 15,879 background genes **(b)**.

*Performance as a function of sample size*

We repeat the ROC process described above for each of the computed LOEUF scores for each downsampling of gnomAD and find that the performance of LOEUF is dependent on sample size and not yet saturated for identifying haploinsufficient genes (Supplementary Fig. 11).

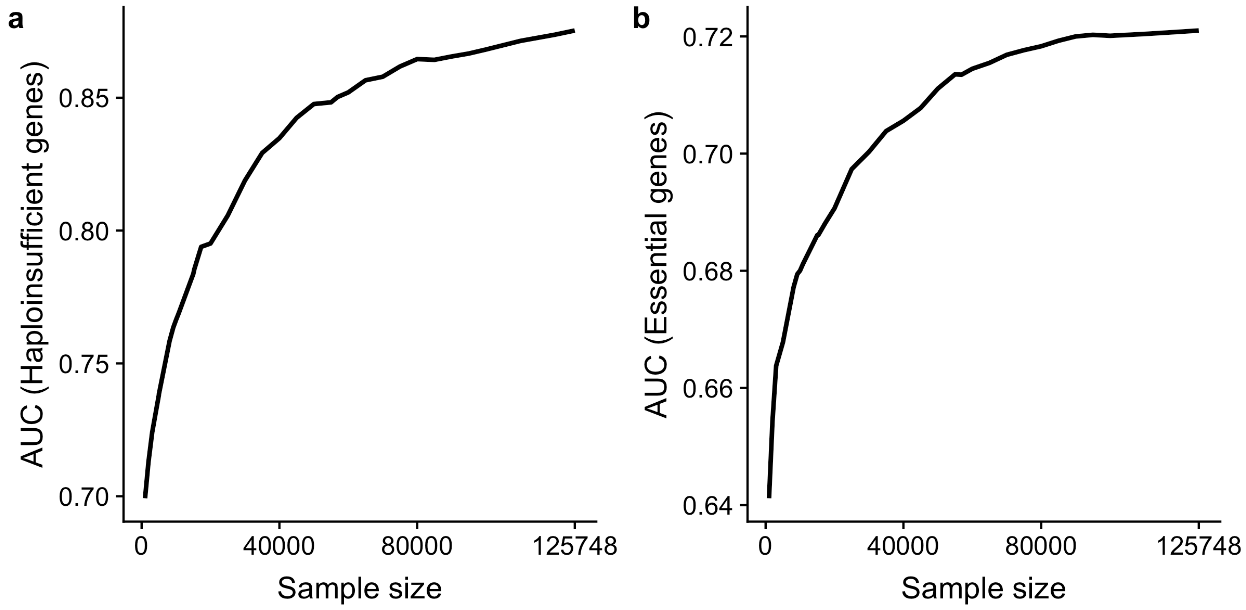


**Supplementary Figure 11 | Performance of LOEUF by sample size.** Area under the ROC curve (AUC) for LOEUF computed for various downsamplings of gnomAD, for discerning 184 haploinsufficient genes from 16,714 background genes **(a)** or 1,019 mouse heterozygous lethal or cell essential genes from 15,879 background genes **(b)**.

### Disease analysis

Andrea Ganna, Raymond Walters, Konrad J. Karczewski, Beryl B. Cummings, Jack A. Kosmicki, Jessica X. Chong, Daniel G. MacArthur, Mark J. Daly, Benjamin M. Neale

#### *Rare disease*

To investigate the constraint spectrum of disease genes, we observed the proportion of genes that are found in the Online Mendelian Inheritance in Man (OMIM, accessed on October 9, 2018; Extended Data Fig. 9a) and find a lower LOEUF in disease genes (mean 0.762 vs 0.992; t-test p-value < $10^{-100}$). In a logistic regression model with coding sequence length as a covariate, LOEUF is correlated with OMIM status (beta = -0.69; p = 4 x $10^{-61}$; gene length beta = 1.3 x $10^{-4}$; p = 1.2 x $10^{-33}$). These genes were further filtered to those underlying monogenic conditions and divided (as described in Chong et al., 2015[5]) into those discovered by whole-exome/whole-genome sequencing (WES/WGS) or previous techniques, such as mapping using linkage or large recurrent chromosomal microduplication/microdeletions, followed by candidate gene sequencing. We find that genes discovered by WES/WGS have lower LOEUF scores than those discovered using conventional linkage techniques, suggesting that the high-throughput technologies are more effective in identifying highly deleterious *de novo* variants in disease genes under extreme constraint against pLoF variation, compared to linkage approaches that are more powerful for identifying inherited variation in moderately constrained genes (Extended Data Fig. 9b). Within OMIM genes, LOEUF is correlated with discovery by WES/WGS compared to conventional approaches (beta = -0.69; p = 2 x $10^{-14}$) when adjusting for coding sequence length (beta = 7.8 x $10^{-6}$; p = 0.54 for the contribution of gene length).

To explore the impact of pLoF variants in constrained genes on neurodevelopmental phenotypes, we collated *de novo* variants from previously published studies, including data from 5,305 probands with intellectual disability / developmental disorders (Hamdam *et al*[75]: n = 41, de

Ligt *et al*[76]: n = 100, Rauch *et al*[77]: n = 51, DDD[78]: n = 4,293, Lelieveld *et al*[79]: n = 820), as well as 6,430 ASD probands and 2,179 unaffected controls[80]. All variants were annotated with VEP v85 against Gencode v19 (identically to the full gnomAD dataset as described above) and annotated with the worst consequence across all affected transcripts, and merged with per-gene LOEUF scores. Genes were filtered to those with at least 10 expected pLoF variants.

To obtain case-control rate ratios and 95% confidence intervals, we calculated the number of pLoF (stop gained, splice donor, splice acceptor, and frameshift) and synonymous variants passing VQSR filters per LOEUF decile in cases and controls and used the estimate from a two-sided Poisson exact test on counts, separately for intellectual disability / developmental disorders (Fig. 5a) and autism (Extended Data Fig. 9c).

To evaluate the utility of LOEUF at partitioning effect sizes for schizophrenia, we utilized a previously published Swedish case control cohort containing 4,133 schizophrenia cases and 9,274 controls[28]. We counted the number of variants seen only once in the dataset (allele count = 1) per LOEUF decile for synonymous and pLoF variants per individual, using the worst consequence of variant accross transcripts. We then performed a logistic regression on case/control status and the number of pLoF and synonymous variants (where the genotype of the individual is non-reference) per LOEUF decile, adjusting for sex, batch, the first 5 principal components, and the total number of variants identified in the sample. The effect size was calculated as the exponent of the beta coefficient from the logistic regression with a 95% confidence interval (Extended Data Fig. 9d).

*Common disease*

Heterozygous pLoF variants in constrained genes are expected to confer some non-trivial survival or reproductive disadvantage[4]. This has been shown empirically, as individuals with diseases characterized by lower reproductive rates carry an excess of rare and *de novo* pLoF variants in constrained genes compared to the general population, which has been shown for disorders such as autism[7], schizophrenia[48], and intellectual disability[78]. Additionally, this class of genetic variation, as a whole, also exerts an effect in the general population beyond the effect on severe neurodevelopmental or psychiatric phenotypes: in particular, individuals that carry a higher number of rare pLoF variants in constrained genes tend to achieve a lower education level on average that those with fewer of these mutations[30].

Following this observation, Ganna and colleagues have explored the impact of rare pLoF variants in constrained genes on 13 quantitative traits and 10 diseases using exome sequencing data from more than 100,000 individuals[29]. As expected, there was no direct association with later onset disorders, which are not characterized by reduced reproductive rates. However, they identified an association with measures of overall health and survival, such as the number of hospitalizations and age at enrollment, suggesting that this class of variants can potentially exert pleiotropic effects on many traits. However, the lack of large exome and genome sequencing studies with many measured phenotypes limits our ability to explore this aspect.

One possibility would be to use common variants surrounding or overlapping constrained genes to understand the impact of these genes on many phenotypes. This can be done in large biobank-scale datasets where common variants have been measured using genotyping technology. However, is it not clear the extent to which inferences drawn from high-impact coding mutations, as discovered by the constraint process, will be reflected in common, typically non-coding, variants. A recent paper suggests a convergence between the impact of rare coding and common variants in constrained genes: Pardiñas *et al.* found that common variants associated with schizophrenia tend to be enriched around genes that are under strong selective pressure[81], similarly to what has been previously shown for rare coding variants[31]. A similar enrichment of both common and rare coding variants has been observed in attention deficit/hyperactivity disorder[35,82]. In this analysis, we seek to explore the contribution of common variants within and near constrained genes across 657 phenotypes.

Specifically, we evaluate enrichment of common variant associations in and near constraint genes using stratified LD Score regression[83]. Briefly, it can be shown that under a basic polygenic model we expect the GWAS $\chi^2$ statistics for SNP *j* to be:

$$E\left[{\chi^2}_j\right] = N\sum_c \tau_c l(j,c) + 1$$

where *N* is the sample size, *c* is the index for the annotation category, $l(j,c)$ is the LD score of SNP *j* with respect to category $C_c$, and $\tau_c$ is the average per-SNP contribution to heritability of category $C_c$. That is, the $\chi^2$ statistic of SNP *j* is expected to be a function of the total sample *N*, how much the SNP tags each category $C_c$ (quantified by $l(j,c)$, the sum of the squared correlation coefficient of SNP *j* with each other SNP in a 1 cM window that is annotated as part of category $C_c$) and $\tau_c$, the effect size of the tagged SNPs.

With this model, LD Score regression allows estimation of each $\tau_c$. Each $\hat{\tau}_c$ is the contribution of category $C_c$ after controlling for all other categories in the model and can be interpreted similarly to a coefficient from a linear regression. Testing for significance of $\hat{\tau}_c$ is useful because it indicates whether the per-SNP contribution to heritability of category *C* is significant after accounting for all the other annotations in the model. However, the scaling of $\hat{\tau}_c$ is also a function of the total $\hat{h}_g^2$ (SNP-based heritability estimate) for the trait, such that an annotation with the same proportional enrichment of signal will have higher $\tau_c$ for traits with higher total $\hat{h}_g^2$. Therefore, to compare the $\hat{\tau}_c$ coefficients across traits (given the same baseline model), we instead use $\hat{\tau}_c^* = \frac{\hat{\tau}_c}{\hat{h}_g^2}$.

In addition to considering the conditional contribution of category $C_c$ with $\hat{\tau}_c^*$, the total marginal heritability explained by SNPs in category $C_c$, denoted $\hat{h}_g^2(C_c)$, is given by

$$\hat{h}_g^2(C_c) = \sum_{C:j\in C_c}\sum_{c':j\in C_{c'}} \hat{\tau}_{c'}$$

In other words, the heritability in category $C_c$ is the sum of the average per-SNP heritability for all SNPs in $C_c$, including contributions to per-SNP heritability from other annotations $c'$ that overlap with category $C_c$ (as indicated by the terms of the inner sum where $c' \neq c$). Importantly, $\hat{h}_g^2(C_c)$ does not depend on the categories chosen to be in the model and provides an easier interpretation.

In this analysis, we will focus first on the *heritability enrichment for category $C_c$* (here, LOEUF decile) which can be obtained by dividing the proportion of total $\hat{h}_g^2$ in $\hat{h}_g^2(C_c)$ to the proportion of all SNPs in $C_c$. An enrichment > 1 indicates that SNPs in category $C_c$ have, on average, a higher per-SNP heritability for a given trait than the average SNP genome-wide.

We applied stratified LD Score regression to summary statistics generated from the Neale lab (https://www.nealelab.is/uk-biobank/) for 4,203 unique phenotypes measured on the UK Biobank. Among these phenotypes, we selected 650 phenotypes that have a significant heritability, as defined by having a heritability p-value < 0.05 after Bonferroni correction for multiple testing. We also included 8 additional sets of summary statistics from large GWAS of cardiovascular diseases, educational attainment, neuroticism and 5 psychiatric/neurodevelopmental disorders (ADHD[82], autism spectrum disorder[84], bipolar disorder[85], major depressive disorder[86] and schizophrenia[81]).

The baseline-LD model[87] for the LD Score regression included 74 annotations that capture different genomic properties including conservation, epigenetic markers, coding regions and LD structure. We also included an annotation that comprised HapMap SNPs 100 kb upstream and downstream of each gene included in the analysis. None of the baseline annotations were multicollinear with the constraint metric, LOEUF. The strongest correlations observed included a metric of background selection ($r^2 = 0.3$)[88] and with regions of high CpG content ($r^2 = 0.37$).

In Fig. 5b, we compared 10 annotations, each defined as a LOEUF decile, and built the corresponding annotation for LD Score regression by considering all the common HapMap variants that were 100 kb upstream and downstream of the start and end of each gene, respectively. Each decile was fit in a separate model, so that we could calculate the enrichment compared to all the other deciles: a SNP was assigned an indicator variable indicating if any gene in the window fell into the given decile. In order to avoid computing biased estimates of the meta-analysis standard errors due to the correlation structure between traits, we identified "independent" traits using the *findCorrelation* function in the R package *caret*, imposing a maximum correlation between variables of 0.2, resulting in 276 independent traits. For each annotation, we report the mean and corresponding standard error of the mean, of a random-effect meta-analysis of the enrichment across the 276 independent traits.

Overall, we observed that annotations containing highly constrained genes are significantly more enriched for heritability across traits than unconstrained genes. In Fig. 5b, we have focused on the enrichment for each annotation across the 276 traits, as this metric is more straightforward to interpret. In Extended Data Fig. 10a, we report the $\hat{\tau}_c^*$ coefficient, which shows a similar trend. Note that positive values of $\hat{\tau}_c^*$ indicate greater per-SNP heritability from SNPs in the annotated category than would be expected based on the other annotations in the baseline-LD model, while negative values indicate depleted per-SNP heritability compared to that baseline expectation.

Next, we verified that our results were robust to changes in the window size around each gene (Extended Data Fig. 10b) and compared windows of 10 kb, 50 kb and 100 kb. The trend in enrichment across LOEUF deciles was consistent across different window sizes, although a proportionally larger enrichment in heritability across traits was observed for 10 kb windows in highly constrained genes as compared to 100 kb windows, whereas the opposite was true for unconstrained genes. This is not unexpected given that a larger window around unconstrained genes is more likely to overlap higher constraint genes than a smaller window, while a smaller window around constrained genes provides a more specific signal around these genes.

We further evaluated whether the heritability enrichment among highly constrained genes could be explained by such genes having higher expression in the brain or being longer than unconstrained genes. We added three additional annotations to the baseline model: 1) the log(TPM) for brain expression from GTEx[14], 2) the number of exons, and 3) the total gene length. Each SNP was annotated with the mean value of each of these annotations for genes within a 100 kb window. In Extended Data Fig. 10c, we report the results for the $\hat{\tau}_c^*$ coefficient across the 10 deciles of LOEUF, after adjusting for these additional annotations, and observed consistent results, indicating that brain expression, gene length, and exon count do not account for the observed heritability enrichments.

Finally, we sought to identify which among the 657 traits showed the strongest constraint enrichment (Fig. 5c). For this analysis, we used LOEUF as a continuous metric in the LD score regression model, and each SNP was annotated with the mean LOEUF score of all genes within a 100 kb window. Because the marginal enrichment values for continuous annotations are not readily interpretable, we instead consider the p-value for the $\hat{\tau}_c^*$ coefficient. This tests whether per-SNP heritability is further enriched proportional to LOEUF beyond the enrichment explained by the baseline annotations. The strongest associations were observed for schizophrenia, educational attainment, and a cognitive test for reaction time (Fig. 5c; Supplementary Table 21).

**Supplementary Table 21 | Phenotypes with association between heritability and constraint.** Only phenotypes with $p < 10^{-4}$ for enrichment based on LD score regression are shown. For all phenotypes, see Supplementary Dataset 14.

| | Enrichment | | | Heritability | |
|---|---|---|---|---|---|
| Description | p-value | Estimate | SE | $h^2$ | p-value |
| Schizophrenia | 1.90E-14 | 0.899 | -0.021 | 0.770 | 1.90E-14 |
| Qualifications: College or University degree | 3.75E-09 | 0.860 | -0.019 | 0.286 | 3.75E-09 |
| Duration to first press of snap-button in each round | 6.51E-08 | 0.893 | -0.022 | 0.081 | 6.51E-08 |
| Educational attainment | 4.32E-07 | 0.885 | -0.016 | 0.115 | 4.32E-07 |
| Bipolar | 4.87E-07 | 0.918 | -0.033 | 0.542 | 4.87E-07 |
| Mean time to correctly identify matches | 8.00E-07 | 0.905 | -0.022 | 0.078 | 8.00E-07 |
| Average total household income before tax | 1.04E-06 | 0.856 | -0.023 | 0.093 | 1.04E-06 |
| Time spend outdoors in summer | 1.08E-06 | 0.856 | -0.029 | 0.068 | 1.08E-06 |
| Variation in diet | 1.54E-06 | 0.886 | -0.025 | 0.048 | 1.54E-06 |
| Answered sexual history questions | 7.27E-06 | 0.756 | -0.051 | 0.081 | 7.27E-06 |
| Time spent outdoors in winter | 3.39E-05 | 0.852 | -0.038 | 0.044 | 3.39E-05 |
| Drive faster than motorway speed limit | 5.00E-05 | 0.846 | -0.030 | 0.058 | 5.00E-05 |
| Frequency of tenseness / restlessness in last 2 weeks | 5.38E-05 | 0.818 | -0.037 | 0.040 | 5.38E-05 |
| Qualifications: None of the above | 6.57E-05 | 0.873 | -0.021 | 0.229 | 6.57E-05 |
| Frequency of tiredness / lethargy in last 2 weeks | 7.24E-05 | 0.876 | -0.025 | 0.061 | 7.24E-05 |
| Alcohol intake versus 10 years previously | 8.38E-05 | 0.849 | -0.041 | 0.031 | 8.38E-05 |

Overall, these results suggest that LOEUF captures common-variant heritability enrichment across many traits independently of existing genomic annotations. Some of the traits, including schizophrenia and educational attainment, that show the strongest enrichment here have also been previously characterized by having a similar enrichment for rare coding variants[29-31]. These results indicate that metrics of gene constraint might provide useful biological information for common diseases and traits.

### Data Availability

Matthew Solomonson, Nick Watts, Konrad J. Karczewski, Grace Tiao, Laurent C. Francioli, Qingbo Wang, Ben Weisburd, Daniel G. MacArthur

##### *Release files*

The gnomAD 2.1 dataset is available for download at our website, http://gnomad.broadinstitute.org. The exome and genome variant datasets are released as sites-level Variant Call Format (VCF) files as well as Hail tables. Per-base coverage summaries are also provided as .txt files and Hail tables.

Additionally, we provide 1,792,248 multi-nucleotide variants (MNVs), which we define as two or more variants within 2 bp existing on the same haplotype in an individual, as .tsv files and Hail tables. These are discussed in detail in a companion manuscript[16]. Within the exome, we created a separate file of 31,575 MNVs existing within a codon reading frame, annotated the functional impact of MNVs on the protein product, which could be different from either or both of the two constituent variants, and displayed these in the browser (see below).

##### *Code availability*

All code to perform quality control, as described in detail above, is provided at https://github.com/macarthur-lab/gnomad_qc: the Sample QC and Annotations code requires access to individual level data, and thus cannot be run directly and is provided for reference only. The code to perform all analyses and regenerate all the figures in this manuscript is provided at https://github.com/macarthur-lab/gnomad_lof. All analyses were done using R 3.6.1 with packages including tidyverse[89], broom[90], magrittr[91], readxl[92], plotROC[93], meta[94], STRINGdb[95], and tidygraph[96]. All visualizations were plotted in ggplot2[97], and aided by scales[98], ggridges[99], egg[100], ggpubr[101], ggrastr[102], cowplot[103], ggrepel[104], and ggwordcloud[105]. LOFTEE is available at https://github.com/konradjk/loftee. All code and software to reproduce figures are available in a Docker image at konradjk/gnomad_lof_paper:0.2.

*The gnomAD Browser*

We have developed a browser to help researchers and clinicians use gnomAD to interpret the impact of genetic variation on gene function. The gnomAD browser's user interface was heavily inspired by the ExAC browser[106], and was engineered using technologies that address the new requirements brought about by the increased size and complexity of gnomAD. Specifically, we have added support for loading, storage, and combining data derived from both whole exome and whole genome sequencing methods, as well as user interface elements for displaying average base pair coverage, quality metrics scores, MNV annotations, and variant summary statistics for WES/WGS data. To accommodate users that require particular sets of samples be excluded from summary statistic calculations, the browser has an option for viewing the prepared subsets of gnomAD (Supplementary Table 11). These gnomAD subsets currently include “controls only”, “non-cancer”, “non-neuro”, or “non-TOPMed”. Nearly all of the data shown in the browser can be queried and downloaded through a GraphQL API (http://gnomad-api.broadinstitute.org), which for some users, obviates the need to transfer, store, and parse very large VCF files. We have organized the codebase with the goal of maximizing code reuse in other genomic data sharing initiatives and to support ongoing development of new features that will aid users in variant interpretation, which will be discussed in more detail in a forthcoming publication. The source code for the browser is available at https://github.com/macarthur-lab/gnomadjs.

This variant data, coverage data, and quality metric data, as well as most orthogonal datasets shown in the browser, were shaped and loaded into an Elasticsearch database (https://www.elastic.co/products/elasticsearch) using Hail (https://hail.is/) running on a Google Cloud Platform Dataproc cluster (https://cloud.google.com/dataproc/). The relevant scripts can be found at https://github.com/macarthur-lab/gnomadjs/tree/master/projects/gnomad/data and https://github.com/macarthur-lab/hail-elasticsearch-pipelines. The front-end code was bundled with webpack (https://webpack.js.org/) and transpiled with Babel (https://babeljs.io/). The view layer of the application was built using React (https://reactjs.org/). Application state is managed in part by Redux (https://redux.js.org/). Visualizations were developed with D3 (https://d3js.org/), HTML Canvas, and scalable vector graphics (SVG). The variant table was rendered with react-virtualized (https://github.com/bvaughn/react-virtualized). Web page elements were styled using styled components (https://www.styled-components.com/). The web server, API, and frontend build processes are run using Node.js (https://nodejs.org/en/). Separate docker images (https://www.docker.com/) were built for each of the gnomAD browser services (web server, API, and databases). The images are deployed and managed by the container orchestration engine Kubernetes (https://kubernetes.io/) running on Google Cloud Platform Kubernetes Engine (https://cloud.google.com/kubernetes-engine/).

##### *Supplementary Datasets*

**Supplementary Dataset 1 | Coverage and mappability summary for each gene by platform.** Well-covered bases are defined as those where at least 80% of the samples sequenced were covered at a depth of at least 20x (10x for males on non-pseudoautosomal regions of the X chromosome). Platforms for exomes are computationally determined as described in “Platform imputation for exomes.”

**Supplementary Dataset 2 | SNV counts in gnomAD exomes.** Number of observed and possible SNVs in the gnomAD exomes, broken down by functional class, variant type (context, reference, and alternative alleles), and median coverage, for each of the computed downsamplings.

**Supplementary Dataset 3 | SNV counts in gnomAD genomes.** Number of observed and possible SNVs in the gnomAD genomes, broken down by functional class, variant type (context, reference, and alternative alleles), and median coverage, for each of the computed downsamplings.

**Supplementary Dataset 4 | Indel counts in gnomAD exomes.** Number of observed indels and positions where such an indel is possible (`coverage_real_estate`) in the gnomAD exomes, broken down by functional class, length, and median coverage, for each of the computed downsamplings.

**Supplementary Dataset 5 | Indel counts in gnomAD genomes.** Number of observed indels and positions where such an indel is possible (`coverage_real_estate`) in the gnomAD genomes, broken down by functional class, length, and median coverage, for each of the computed downsamplings.

**Supplementary Dataset 6 | Results for the Kolmogorov-Smirnov test and Mood's median test on the allele balance and age distributions for each transcript.** The uncorrected p-values are shown for each test (two-sided KS test and Mood’s median test), based on 85,462 of the exome-sequenced individuals. Transcripts were omitted from the results if data were insufficient to run the analyses and resulted in p-values of "NA" for the Kolmogorov-Smirnov tests for both allele balance and age. Summed histograms representing the total counts of each bin value for pLoF (`summed_ab_hists_lof` and `summed_age_hists_lof`) and synonymous (`summed_ab_hists_synonymous` and `summed_age_hists_synonymous`) variants within the transcript and the median value are also reported. The allele balance bins range from 0 to 1.0 in increments of 0.05. Age bins range from 25 to 80 in increments of 5 with values <25 and >80 grouped into the first and last bin, respectively.

**Supplementary Dataset 7 | List of genes where at least one confident homozygous pLoF genotype was observed.** See “Homozygous variant curation” for more details.

**Supplementary Dataset 8 | Total number of alleles observed by population by functional class in the gnomAD exomes.** The aggregate number of alleles (`total`) and homozygous individuals (`total hom`) are shown.

**Supplementary Dataset 9 | Total number of alleles observed by population by functional class in the gnomAD genomes.** The aggregate number of alleles (`total`) and homozygous individuals (`total hom`) are shown.

**Supplementary Dataset 10 | Mutation rates.** The estimated mutation rate for each context, reference, and alternate allele, split by methylation status for CpG transitions.

**Supplementary Dataset 11 | Constraint metrics.** Note that this file contains all transcripts: for gene-based analyses, the file should be filtered to canonical transcripts (`canonical == true`), and LOEUF decile bin (`oe_lof_upper_bin`) recomputed for each gene. The most commonly used metrics in this manuscript are `oe_lof_upper`, `oe_lof_upper_bin`, and `p`. Columns are:

- gene: Gene name
- transcript: Ensembl transcript ID (Gencode v19)
- canonical: Boolean indicator as to whether the transcript is the canonical transcript for the gene
- obs_mis: Number of observed missense variants in transcript
- exp_mis: Number of expected missense variants in transcript
- oe_mis: Observed over expected ratio for missense variants in transcript (obs_mis divided by exp_mis)
- mu_mis: Mutation rate summed across all possible missense variants in transcript
- possible_mis: Number of possible missense variants in transcript
- obs_mis_pphen: Number of observed missense variants in transcript predicted "probably damaging" by PolyPhen-2
- exp_mis_pphen: Number of expected missense variants in transcript predicted "probably damaging" by PolyPhen-2
- oe_mis_pphen: Observed over expected ratio for PolyPhen-2 predicted "probably damaging" missense variants in transcript (obs_mis_pphen divided by exp_mis_pphen)
- possible_mis_pphen: Number of possible missense variants in transcript that are predicted "probably damaging" by PolyPhen-2
- obs_syn: Number of observed synonymous variants in transcript

- exp_syn: Number of expected synonymous variants in transcript
- oe_syn: Observed over expected ratio for missense variants in transcript (obs_syn divided by exp_syn)
- mu_syn: Mutation rate summed across all synonymous variants in transcript
- possible_syn: Number of possible synonymous variants in transcript
- obs_lof: Number of observed predicted loss-of-function (pLoF) variants in transcript
- mu_lof: Mutation rate summed across all possible pLoF variants in transcript
- possible_lof: Number of possible pLoF variants in transcript
- exp_lof: Number of expected pLoF variants in transcript
- pLI: Probability of loss-of-function intolerance; probability that transcript falls into distribution of haploinsufficient genes (~9% o/e pLoF ratio; computed from gnomAD data)
- pRec: Probability that transcript falls into distribution of recessive genes (~46% o/e pLoF ratio; computed from gnomAD data)
- pNull: Probability that transcript falls into distribution of unconstrained genes (~100% o/e pLoF ratio; computed from gnomAD data)
- oe_lof: Observed over expected ratio for pLoF variants in transcript (obs_lof divided by exp_lof)
- oe_syn_lower: Lower bound of 90% confidence interval for o/e ratio for synonymous variants
- oe_syn_upper: Upper bound of 90% confidence interval for o/e ratio for synonymous variants
- oe_mis_lower: Lower bound of 90% confidence interval for o/e ratio for missense variants
- oe_mis_upper: Upper bound of 90% confidence interval for o/e ratio for missense variants
- oe_lof_lower: Lower bound of 90% confidence interval for o/e ratio for pLoF variants
- oe_lof_upper: LOEUF: upper bound of 90% confidence interval for o/e ratio for pLoF variants (lower values indicate more constrained)
- constraint_flag: Reason gene does not have constraint metrics. One of:
    - no variants: Zero observed synonymous, missense, pLoF variants
    - no_exp_syn: Zero expected synonymous variants
    - no_exp_mis: Zero expected missense variants
    - no_exp_lof: Zero expected pLoF variants
    - syn_outlier: Too many or too few synonymous variants; synonymous z score < -5 or synonymous z score > 5
    - mis_too_many: Too many missense variants; missense z score < -5
    - lof_too_many: Too many pLoF variants; pLoF z score < -5
- syn_z: Z score for synonymous variants in gene. Higher (more positive) Z scores indicate that the transcript is more intolerant of variation (more constrained). Extreme values of syn_z indicate likely data quality issues
- mis_z: Z score for missense variants in gene. Higher (more positive) Z scores indicate that the transcript is more intolerant of variation (more constrained)
- lof_z: Z score for pLoF variants in gene. Higher (more positive) Z scores indicate that the transcript is more intolerant of variation (more constrained)
- oe_lof_upper_rank: Transcript's rank of LOEUF value compared to all transcripts (lower values indicate more constrained)
- oe_lof_upper_bin: Decile bin of LOEUF for given transcript (lower values indicate more constrained)
- oe_lof_upper_bin_6: Sextile bin of LOEUF for given transcript (lower values indicate more constrained)

- n_sites: Number of distinct pLoF variant sites in the transcript
- classic_caf: Sum of allele frequencies of pLoFs in the transcript
- max_af: Maximum allele frequency of any pLoF in the transcript
- no_lofs: The number of individuals with no observed pLoF variants in the transcript
- obs_het_lof: The number of individuals with at least one observed heterozygous pLoF variant, but no homozygous pLoF variants, in the transcript
- obs_hom_lof: The number of individuals with at least one observed homozygous pLoF in the transcript
- defined: The number of individuals where at least one high-quality genotype (including homozygous reference) is observed at a called site annotated as a pLoF variant
- p: The estimated proportion of haplotypes with a pLoF variant. Defined as: 1 - sqrt(no_lofs / defined)
- exp_hom_lof: The expected number of individuals with at least one homozygous pLoF variant based on the frequency of pLoF haplotypes. Defined as: defined * $p^2$
- classic_caf_afr: Sum of allele frequencies of pLoFs in the transcript among African/African-American individuals
- classic_caf_amr: Sum of allele frequencies of pLoFs in the transcript among Latino individuals
- classic_caf_asj: Sum of allele frequencies of pLoFs in the transcript among Ashkenazi Jewish individuals
- classic_caf_eas: Sum of allele frequencies of pLoFs in the transcript among East Asian individuals
- classic_caf_fin: Sum of allele frequencies of pLoFs in the transcript among Finnish individuals
- classic_caf_nfe: Sum of allele frequencies of pLoFs in the transcript among Non-Finnish European individuals
- classic_caf_oth: Sum of allele frequencies of pLoFs in the transcript among Other (uncharacterized ancestry) individuals
- classic_caf_sas: Sum of allele frequencies of pLoFs in the transcript among South Asian individuals
- p_afr: The computation of `p` repeated among only African/African-American individuals
- p_amr: The computation of `p` repeated among only Latino individuals
- p_asj: The computation of `p` repeated among only Ashkenazi Jewish individuals
- p_eas: The computation of `p` repeated among only East Asian individuals
- p_fin: The computation of `p` repeated among only Finnish individuals
- p_nfe: The computation of `p` repeated among only Non-Finnish European individuals
- p_oth: The computation of `p` repeated among only Other (uncharacterized ancestry) individuals
- p_sas: The computation of `p` repeated among only South Asian individuals
- transcript_type: Transcript biotype (https://www.gencodegenes.org/pages/biotypes.html)
- gene_id: Ensembl gene ID
- transcript_level: Transcript level from Gencode (https://www.gencodegenes.org/pages/data_format.html)
- cds_length: Length of coding sequence in gene
- num_coding_exons: Number of coding exons in gene
- gene_type: Gene biotype (https://www.gencodegenes.org/pages/biotypes.html)
- gene_length: Length of gene
- exac_pLI: pLI score calculated from ExAC
- exac_obs_lof: Number of observed pLoF variants in gene in ExAC
- exac_exp_lof: Number of expected pLoF variants in gene in ExAC
- exac_oe_lof: Observed to expected ratio of pLoF variants in ExAC

- brain_expression: Expression of gene in brain from GTEx data
- chromosome: Chromosome name
- start_position: Start position of gene
- end_position: End position of gene

**Supplementary Dataset 12 | Estimated number of individuals required to reach an expected number of variants for each gene.** See “Summary of constraint metrics.” The number of individuals required (`n_required`) in order to achieve an expected number of variants (`n_variants`) and its percentile rank (`rank`) is shown for each functional class (`variant_type`).

**Supplementary Dataset 13 | Gene lists.** Membership of each gene in haploinsufficient, autosomal recessive, and olfactory gene lists described in Fig. 3a, Supplementary Figure 9, and Supplementary Table 18.

**Supplementary Dataset 14 | Summary of enrichments by phenotype.** Table with summary statistics from partitioning heritability analysis with LOEUF as a covariate for 657 traits.

### Acknowledgments

ATVB & Precocious Coronary Artery Disease Study (PROCARDIS): Exome sequencing was supported by a grant from the NHGRI (5U54HG003067-11) to Drs. Gabriel and Lander. Bulgarian Trios: Medical Research Council (MRC) Centre (G0800509) and ProgramGrants (G0801418), the European Community's Seventh Framework Programme (HEALTH-F2-2010-241909 (Project EU-GEI)), and NIMH(2P50MH066392-05A1). GoT2D & T2DGENES: NHGRI ("Large Scale Sequencing and Analysis of Genomes" U54HG003067), NIDDK ("Multiethnic Study of Type 2 Diabetes Genes" U01DK085526), NIH ("LowF Pass Sequencing and High Density SNP Genotyping in Type 2 Diabetes" 1RC2DK088389), National Institutes of Health ("Multiethnic Study of Type 2 Diabetes Genes" U01s DK085526, DK085501, DK085524, DK085545, DK085584; "LowF Pass Sequencing and HighF Density SNP Genotyping for Type 2 Diabetes" DK088389). The German Center for Diabetes Research (DZD). National Institutes of Health (RC2F DK088389, DK085545, DK098032). Wellcome Trust (090532, 098381). National Institutes of Health (R01DK062370, R01DK098032, RC2DK088389). The Academy of Finland Center of Excellence for Complex Disease Genetics (grant number 312063, Sigrid Jusélius Foundation). METSIM: Academy of Finland and the Finnish Cardiovascular Research Foundation. Inflammatory Bowel Disease: The Helmsley Trust Foundation, #2015PG-IBD001, Large Scale Sequencing and Analysis of Genomes Grant (NHGRI), 5 U54 HG003067-13, National Institute of Diabetes and Digestive and Kidney Diseases (NIDDK; DK062432), Canada Research Chair (#230625). Jackson Heart Study: We thank the Jackson Heart Study (JHS) participants and staff for their contributions to this work. The JHS is supported by contracts HHSN268201300046C, HHSN268201300047C, HHSN268201300048C, HHSN268201300049C, HHSN268201300050C from the National Heart, Lung, and Blood Institute and the National Institute on Minority Health and Health Disparities. Ottawa Genomics Heart Study: Canadian Institutes of Health Research MOP136936; MOP82810, MOP77682,

Canadian Foundation for Innovation 11966, Heart & Stroke Foundation of Canada T-7268. Exome sequencing was supported by a grant from the NHGRI (5U54HG003067-11) to Drs. Gabriel and Lander. Pakistan Risk of Myocardial Infarction Study (PROMIS): Exome sequencing was supported by a grant from the NHGRI (5U54HG003067-11) to Drs. Gabriel and Lander. Fieldwork in the study has been supported through funds available to investigators at the Center for Non-Communicable Diseases, Pakistan and the University of Cambridge, UK. Registre Gironi del COR (REGICOR): Spanish Ministry of Economy and Innovation through the Carlos III Health Institute [Red HERACLES RD12/0042, CIBER Epidemiología y Salud Pública, PI12/00232, PI09/90506, PI08/1327, PI05/1251, PI05/1297], European Funds for Development (ERDF-FEDER), and by the Catalan Research and Technology Innovation Interdepartmental Commission [SGR 1195]. Swedish Schizophrenia & Bipolar Studies: National Institutes of Health (NIH)/National Institute of Mental Health (NIMH) ARRA Grand Opportunity grant NIMHRC2MH089905, the Sylvan Herman Foundation, the Stanley Center for Psychiatric Research, the Stanley Medical Research Institute, NIH/National Human GenomeResearch Institute (NHGRI) grant U54HG003067. SIGMA-T2D: The work was conducted as part of the Slim Initiative for Genomic Medicine, a project funded by the Carlos Slim Health Institute in Mexico. The UNAM/INCMNSZ Diabetes Study was supported by Consejo Nacional de Ciencia y Tecnologıía grants 138826, 128877, CONACT- SALUD 2009-01-115250, and a grant from Dirección General de Asuntos del Personal Académico, UNAM, IT 214711. The Diabetes in Mexico Study was supported by Consejo Nacional de Ciencia y Tecnología grant 86867 and by Instituto Carlos Slim de la Salud, A.C. The Mexico City Diabetes Study was supported by National Institutes of Health (NIH) grant R01HL24799 and by the Consejo Nacional de Ciencia y Tenologia grants 2092, M9303, F677-M9407, 251M, and 2005-C01-14502, SALUD 2010-2-151165. Schizophrenia Trios from Taiwan: NIH/NIMH grant R01MH085560 and R01MH085521. Tourette Syndrome Association International Consortium for Genomics (TSAICG): NIH/NINDS U01 NS40024-09S1. Leicester Exome Sequencing Cohort: British Heart Foundation (grant

CS/14/2/30841). Framingham Heart Study: National Heart, Lung and Blood Institute's Framingham Heart Study Contract (HHSN268201500001I); National Institute for Diabetes and Digestive and Kidney Diseases (NIDDK) R01 DK078616. SISU: Academy of Finland (grants 263278, 265240, 286500, 293404, 301220, 307866, 308248, 312073), Sigrid Jusélius Foundation, the Strategic Neuroscience Funding of the University of Eastern Finland, the Academy of Finland Center of Excellence for Complex Disease Genetics (grant number 312074)
